# Replication dynamics of individual loci in single living cells reveal variation of stochasticity

**DOI:** 10.1101/485987

**Authors:** Bénédicte Duriez, Sabarinadh Chilaka, Jean-François Bercher, Eslande Hercul, Nicole Boggetto, Marie-Noëlle Prioleau

## Abstract

Eukaryotic genomes are replicated under the control of a highly sophisticated program during the restricted time period corresponding to S-phase. The most widely used replication timing assays, which are performed on populations of millions of cells, suggest that most of the genome is synchronously replicated on homologous chromosomes. We investigated the stochastic nature of this temporal program, by comparing the precise replication times of allelic loci within single vertebrate cells progressing through S-phase at six loci replicated from very early to very late. We show that replication timing is strictly controlled for the three loci replicated in the first half of S-phase. Out of the three loci replicated in the second part of S-phase, two present a significantly more stochastic pattern. Surprisingly, we find that the locus replicated at the very end of S-phase, presents stochasticity similar to those replicated in early S-phase. We suggest that the richness of loci in efficient origins of replication, which decreases from early-to late-replicating regions, may underlie the variation of timing control during S-phase.

## Introduction

At each cell division, the genome must be entirely and faithfully duplicated during the short time period corresponding to S-phase. DNA replication errors, such as genomic rearrangements, may have damaging consequences, leading to cell death or tumorigenesis. Intensive research on the DNA replication program has revealed that is subject to a highly sophisticated process tightly regulating its execution in space and time (Fragkos *et al*, 2015). DNA replication is initiated at a large number of sites, known as origins of replication, on the chromosomes of eukaryotic cells (Méchali, 2010; Prioleau & MacAlpine, 2016). The number of potential origins licensed in G1 phase is larger than the number of origins activated in S-phase in each cell. This is thought to reflect flexible origin choice and to be directly related to the stochastic nature of the eukaryotic replication program. Several factors, such as primary sequence, chromatin landscape and *trans*-factors may influence the activation of origins (Besnard *et al*, 2012; Cadoret *et al*, 2008; Cayrou *et al*, 2011, 2012; Costas *et al*, 2011; Picard *et al*, 2014; Sequeira-Mendes *et al*, 2009; Valton *et al*, 2014).

Not only do eukaryotic origins of replication display spatial organization and flexible activation during the cell cycle, but their activation is also subject to temporal regulation. This programmed timing of replication results in a succession of domains of hundreds of kilobases to megabases in length, located along the entire chromosome, being replicated at about the same time (Renard-Guillet *et al*, 2014). These domains replicate in early, mid or late S-phase, and about 50% are cell type-specific in metazoans (Rhind & Gilbert, 2013; Ryba *et al*, 2010). The factors responsible for the establishment, regulation and maintenance of these domains throughout the cell cycle remain largely unknown. As a general rule, GC-rich euchromatin regions with a high density of active genes replicate early in S phase, whereas heterochromatin and AT-rich regions replicate late in S-phase. Only three *trans*-factors (Rif1, Fkh1/2 and USF) have been directly implicated in controlling the timing of replication in yeast and vertebrate cells (Foti *et al*, 2016; Hassan-Zadeh *et al*, 2012; Knott *et al*, 2012; Peace *et al*, 2016; Reinapae *et al*, 2017; Yamazaki *et al*, 2013).

Two genome-wide studies in the haploid yeast *S. cerevisiae* have assessed the impact of stochastic origin activation on replication dynamics (Hawkins *et al*, 2013; McGuffee *et al*, 2013). Replication initiation was found to be stochastic, as different cohorts of origins initiated DNA replication in different cells. Not only was the choice of origins stochastic, but so was the timing of their activation, resulting in significant cell-to-cell variability in genome replication (Hawkins *et al*, 2013; McGuffee *et al*, 2013). As previously suggested by modeling data (Yang *et al*, 2010), there is a positive correlation between median origin activation time and range of activation time, consistent with greater variability of activation timing for origins activated later in the cell cycle. Thus, late origins tend to fire over larger time windows than early origins (Hawkins *et al*, 2013). The measurement of replication time windows in diploid vertebrate cells, through comparisons of replication timing for allelic loci, can be used to determine whether replication dynamics follows the same rules in vertebrates. The chief advantage of this approach is that it prevents bias due to slight differences in cell synchronization, supposing that comparisons are made within single cells.

Most timing analyses performed in vertebrates to date have measured the average timing of the two alleles of individual loci in a cell population (Ryba *et al*, 2011). However, three recent genome-wide studies established allele-specific replication timing maps in humans (Koren & McCarroll, 2014; Mukhopadhyay *et al*, 2014; Koren *et al*, 2014) and in mouse (Rivera-Mulia *et al*, 2018). They demonstrated a high degree of similarity in autosome replication profiles between individuals or clones and experimental replicates (Koren & McCarroll, 2014; Mukhopadhyay *et al*, 2014; Rivera-Mulia *et al*, 2018). Mukhopadhyay *et al.* reported that human chromosome homologs replicated highly synchronously, within less than 48 minutes of each other, over about 88% of the genome. The remaining 12% of the genome could be divided into about 600 regions with less synchronous replication, with an average time lag in firing of 50 min to 150 min. The authors suggested that these regions might be associated with large structural variants and that most asynchronous regions were enriched in imprinted genes (Mukhopadhyay *et al*, 2014). Among six hybrid mESC clones, with different combinations of three different genomes, only cell lines derived from rather distantly species contain regions with asynchronous replication between alleles (12% of the genome has a time lag in firing above 80 min). The only parameter that distinguishes these regions from the rest of the genome is their subspecies origin (Rivera-Mulia *et al*, 2018). Koren *et al.* also investigated possible changes in the control of replication timing during S-phase in human lymphoblastoid cell lines. They observed a gradual loss of replication structure with the progression of S-phase (Koren & McCarroll, 2014), as previously reported for *S. cerevisiae* (Hawkins *et al*, 2013; McGuffee *et al*, 2013).

These allele-specific replication timing analyses were performed on millions of cells. They thus measured the average replication timing of million alleles but not the variation from allele to allele in individual cells (Koren & McCarroll, 2014; Mukhopadhyay *et al*, 2014; Rivera-Mulia *et al*, 2018). Only regions subject to imprinting or clearly devoid of a structured replication program would be recognized as asynchronously replicated regions in these conditions. This global method is therefore inappropriate for the evaluation of intrinsic parameters of the stochastic nature of replication timing. A recent study has addressed the question of the stochastic variation in mouse replication timing through the comparison of homologs in single cells (Dileep & Gilbert 2018). They found that replication timing domains in single cells are similar to the ones described in population-based assays, thereby highlighting the strong control of replication timing. They also reported that stochastic variation in replication timing is similar between cells and between homologs regardless of their replication timing. However, data from only 10 cells collected in mid S-phase were analyzed, therefore stochastic properties of replication timing throughout the S-phase could not be examined with a high resolution.

We decided to study replication dynamics in greater detail, by tracking the replication of the two alleles of individual loci in single cells progressing through S-phase. Real-time analyses of this type have a very high temporal resolution and are very suitable for evaluations of the stochastic properties of replication timing.

## Results

### Experimental design for analyzing the replication timing of the two alleles at a given locus in single-cells by real-time live-cell imaging

We evaluated the stochastic nature of replication timing at a single locus, by determining and comparing the replication timings of two alleles at the same locus in single cells by live-cell imaging. We used the DT40 chicken B-cell line, in which genetic manipulation is easier than in mammalian cell lines. The ratio of targeted to random DNA insertions in this cell line is indeed much higher than in mammalian cell lines, facilitating the site-specific insertion of transgenes for both alleles of the locus of interest. The DT40 cell line also has a short generation time and a high proportion of cells in S-phase, and is therefore highly suitable for analyses of the replication program. This cell line has the same origin properties and patterns of timing for DNA replication as reported for mammalian models. We and others have already successfully used this model to address questions concerning the DNA replication process in vertebrates (Valton *et al*, 2014; Hassan-Zadeh *et al*, 2012; Schiavone *et al*, 2016; Petermann *et al*, 2006).

We tracked the replication of single loci, by adding a fluorescent tag to the locus of interest and measuring the intensity of fluorescence over S-phase: fluorescence intensity doubles at the time point at which the locus is replicated. This approach has already been used in haploid or diploid yeast to examine relative replication time of two loci that are 20 to 200 kb apart, either on one chromosome or on homologous chromosomes (Kitamura *et al*, 2006; Saner *et al*, 2013; Dovrat *et al*, 2018). The tag used consisted of an array of 120 repeats of the bacterial tetracycline operator (TetO) bound to the tetracycline repressor fused to the EGFP protein (TetR-EGFP) (Fig. 1A).

**Figure 1.**
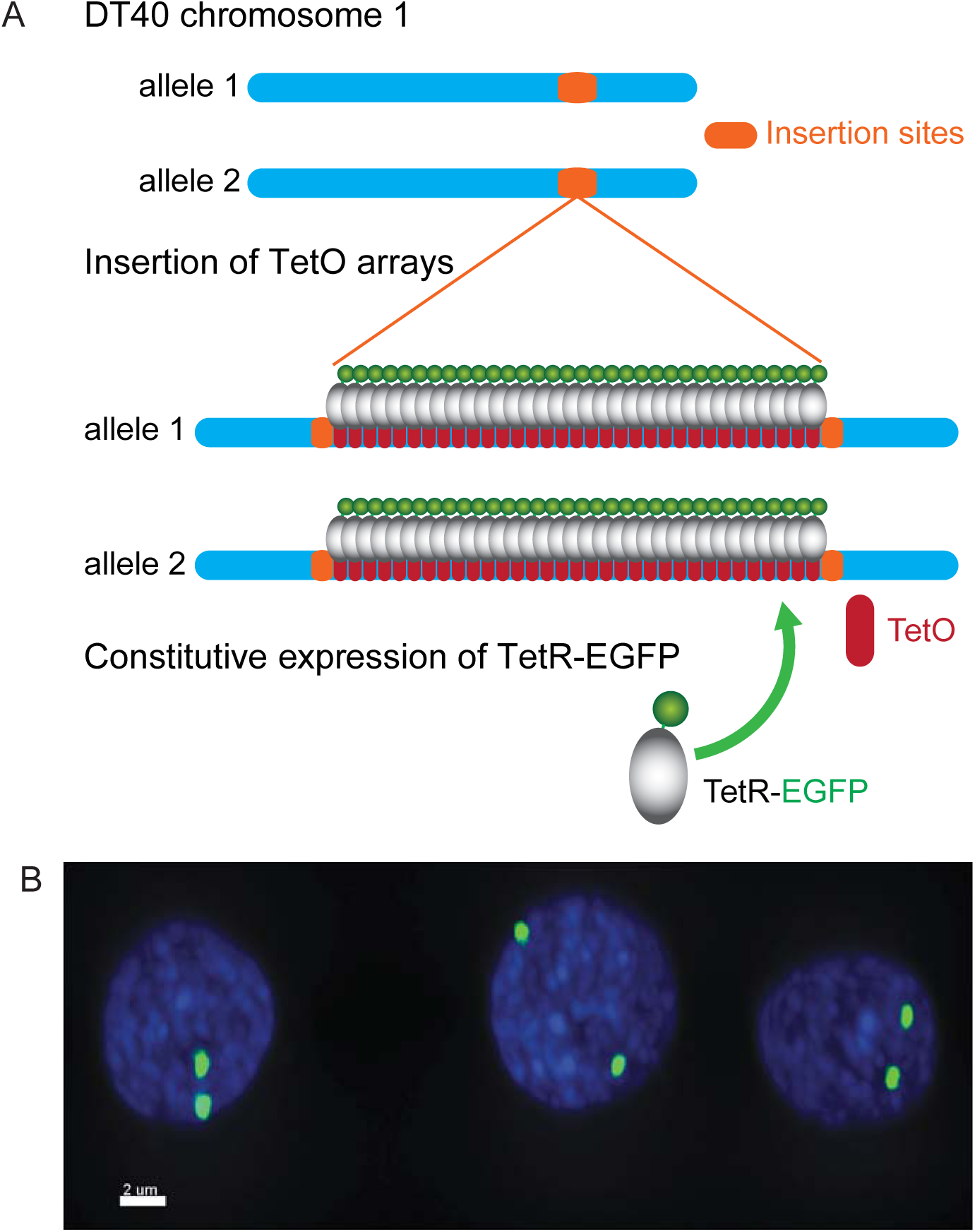
Construction of cell lines with a fluorescent tag on both alleles of a single locus. (A) Schematic representation of the insertion of an array of 120 repeats of the tetracycline operator (TetO) (red boxes) into both alleles of the targeted locus (orange box). The cell line constitutively expresses the TetO repressor fused to enhanced-GFP (TetR-EGFP) which binds to TetO arrays. (B) The visualization of two bright fluorescent spots in the nucleus of each cell of the constructed cell lines indicates the labeling of the two targeted alleles. Cells were imaged with a confocal microscope. The fluorescent EGFP spots are visualized in green, and the chromatin labeled with Hoechst 33342 is shown in blue (image of three cells from Early 2 cell line). Scale bar, 2 µm.

Six loci were selected, on the basis of the DT40 replication timing map established in our laboratory (Hassan-Zadeh *et al*, 2012), to illustrate the three different types of timing domains: early-, mid-late- and late-replicating domains (Fig. 2A and Supplementary Fig. 1). Moreover, a more precise RT analysis in wild-type cells showed that RT of these loci was distributed homogenously along S-phase from a very early profile (Early 1) to a very late profile (Late 2) (Fig. 2C). Six different cell lines were constructed. In each cell line, the TetO array was inserted into both alleles of the locus of interest in a cell line expressing TetR-EGFP (Supplementary Fig. 2). The cell lines were named “Early 1”, “Early 2”, “Mid-late 1”, “Mid-late 2”, “Late 1” and “Late 2” cell line, according to the replication timing of the locus tagged. A seventh cell line was established by inserting one TetO array into one allele of the Early 2 locus and a second TetO array into one allele of the Mid-late 2 locus. This cell line was named “Early 2 + Mid-late 2” cell line.

**Figure 2.**
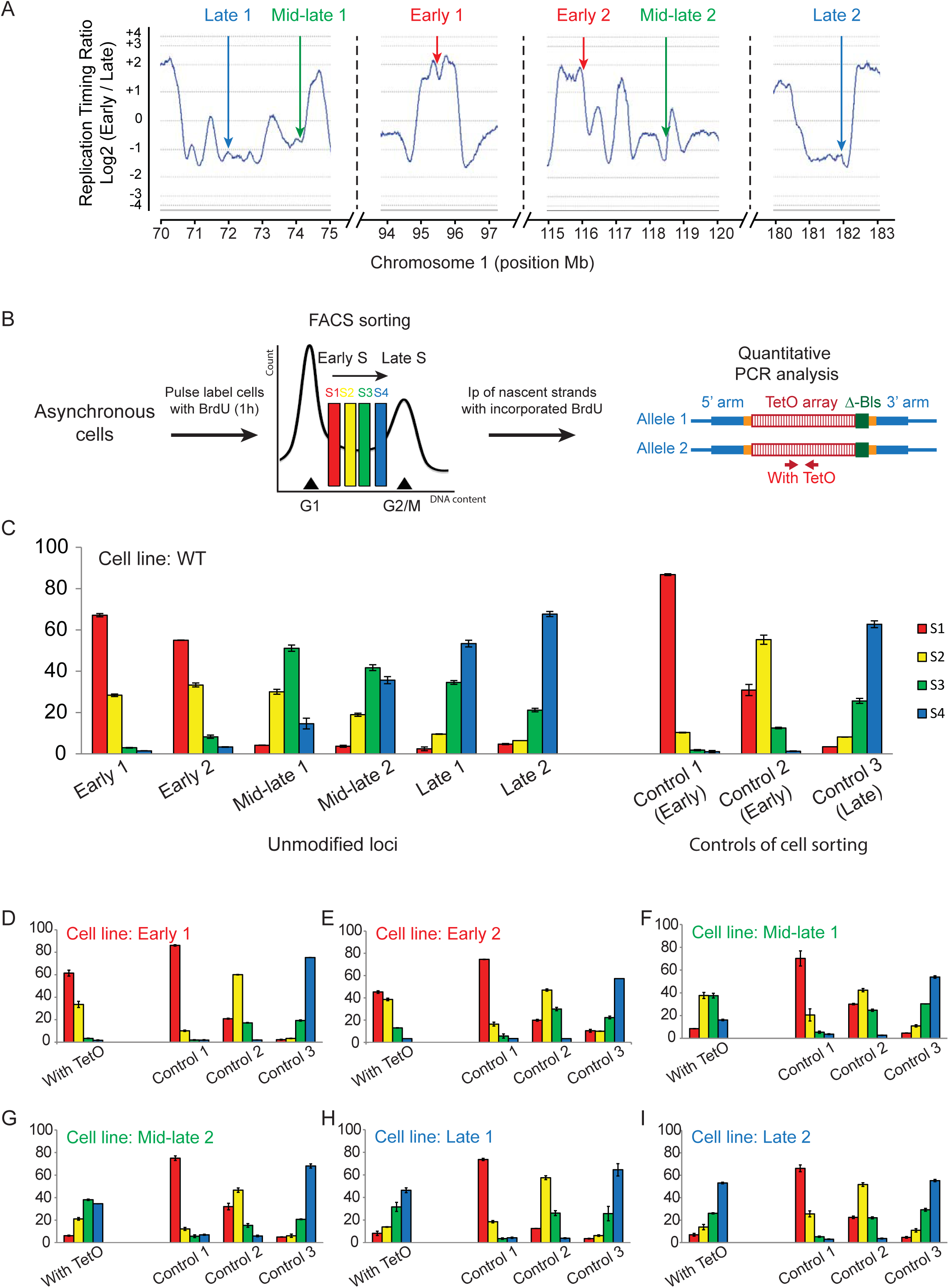
Domains with different replication timing profiles were investigated. (A) A portion of the DT40 chromosome 1 replication timing map, previously established in our laboratory (Hassan-Zadeh *et al*, 2012), is shown. Values of the log_2_-ratio (early/late) (blue curve) define the timing profile along the chromosome, with positions indicated on the *x* axis (Assembly WUGSC 2.1/galGal3 May 2006). Six domains with different timing profiles (early, mid-late and late) were selected and the chosen insertion sites are indicated (red, green and blue arrows). (B) The precise replication timing profiles for the six insertion loci were determined in quantitative timing assays. Asynchronously growing cells were subjected to BrdU pulse-labeling. Four S-phase fractions (fractions S1 to S4, covering the whole of S-phase, from early to late S-phase) were subjected to cell sorting (FACS analysis). Nascent strands were purified and quantified by real-time PCR with primers amplifying targeted sites. The replication timing profiles of three control loci (Control 1 (MED14), control 2 (Rho), and control 3 (UPDYS2)) were used to evaluate the sorting. (C) Replication timing profiles are shown in the WT DT40 cell line, for the six insertion sites (unmodified loci). (D-I) The replication timing profile of the modified locus was determined in each of the six homozygous cell lines with TetO inserted into both alleles. The replication of the modified Early 2 locus appeared to occur slightly later in the Early 2 cell line than that of the unmodified locus in the WT cell line, but this modified locus was still replicated early, as most of the nascent strands were found in the S1 and S2 fractions (84%). Consistent with these findings, the control 2 locus also displayed a slight delay in replication, confirming a slight difference in the cell sorting. A similar situation was observed with the Mid-late 1 locus in the Mid-late 1 cell line where the modified locus was mostly replicated during mid S-phase. The replication profiles of Early 1, Mid-late 2, Late 1 and Late 2 loci were very similar in the homozygous cell lines and the WT cell line. Error bars indicate standard deviation for qPCR duplicates.

We firstly checked that the replication timing of the tagged loci was not altered by the presence of the TetO arrays bound to TetR-EGFP. We compared the replication timing of the wild-type and TetO-tagged alleles in the six heterozygous cell lines. Timing experiments were performed as previously described (Hassan-Zadeh *et al*, 2012) (Supplementary Fig. 3A). We found that the two alleles had similar timing profiles, with a slight tendency towards a short delay for the tagged allele in the Mid-late 2 cell line (Supplementary Fig. 3E). We assumed that the insertion of exactly the same sequence into both alleles would have a similar slight impact on both alleles. Replication of early loci occurred mostly in the first part of S-phase (S1+S2= 95% and 88%, in the Early 1 locus and in the Early 2 locus, respectively). Replication of mid-late loci were more concentrated in the mid part of S-phase (S2+S3= 81% and 61% in the Mid-late 1 locus and in the Mid-late 2 locus, respectively). The late loci replicated mostly in the last part of S-phase (S3+S4= 88% and 89% in the Late 1 locus and in the Late 2 locus, respectively). We compared results obtained in wild-type cells with those obtained in each of the six homozygous cell lines with TetO arrays insertion on both alleles. We found that replication timing of homozygous tagged loci is highly similar to replication timing of corresponding loci in wild-type cell line (Fig. 2D-I). The replication timing profiles of the modified loci were, thus, preserved in each of the six cell lines. We then checked that the TetO arrays were correctly detected with TetR-EGFP proteins. The inserted arrays were visualized as two bright fluorescent spots in the nucleus of each cell of the different cell lines (illustrated for the Early 1 cell line in Fig.1B).

We determined the precise replication timing of the two alleles of the locus of interest, by tracking the two fluorescent spots within single cells as they progressed through S-phase, which lasts about seven hours. Briefly, cells were synchronized in G1 by elutriation without drug addition, to prevent possible disturbance of progression through the cell cycle (Fig. 3A). Early S-phase cells were cultured in dishes under a confocal microscope and images were acquired at five-minute intervals, providing a very high temporal resolution. The fluorescence intensity of each spot was measured at each time point to determine the respective replication timings of the two alleles on the basis of the time point at which fluorescence intensity doubled (Kitamura *et al*, 2006). We calculated the replication delay (RD) as the time interval separating the replication of the two alleles within a single cell (Fig. 3B-C and Supplementary Fig. 4 and Fig. 5).

**Figure 3.**
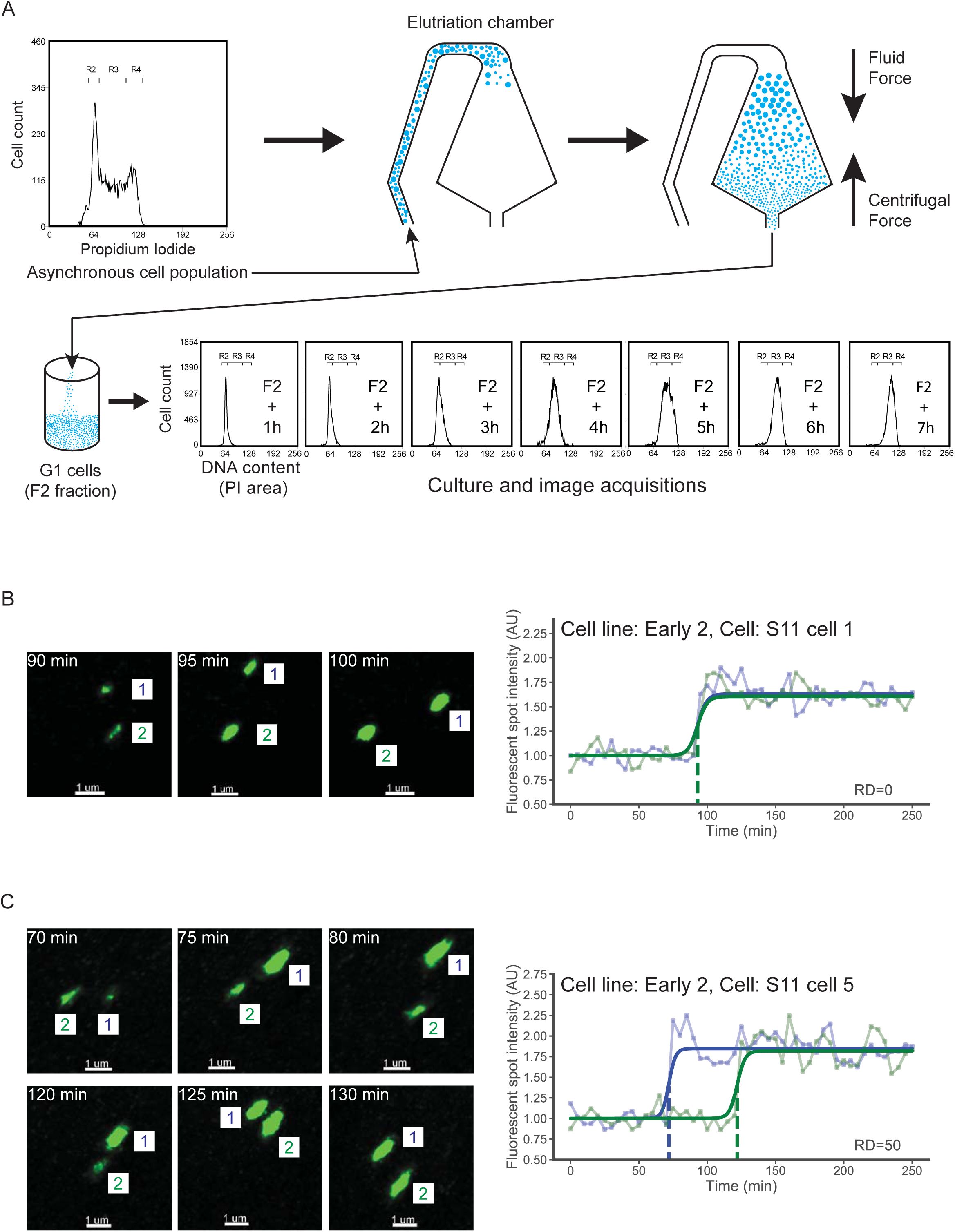
Measurement of replication delays between the two alleles in individual cells. (A) Schematic illustration of the elutriation process for cell separation on the basis of size through the opposition of fluid and centrifugal forces. The G1 cells, which are the smallest, were purified in this way and cultured in normal conditions for several hours. At one-hour intervals, the progression of the cultured cells through the cell cycle was evaluated by flow cytometry with PI incorporation for the measurement of DNA content. Seven successive profiles showed that S-phase took about seven hours to complete. (B) Real-time imaging of individuals cells and measurement of spot intensities: illustration of the synchronous replication of both alleles in one cell of the Early 2 cell line. G1 cells were cultured under an inverted confocal microscope and 3D images were acquired for the GFP channel, at five-minute intervals. The two spots are numbered 1 and 2 to identify the two alleles. Both alleles of the illustrated cell underwent replication between the image acquisition time points at 90 and 95 min. On the right, variation of the intensity of GFP fluorescence for the two spots (blue and green curves), together with replication time (blue and green dotted lines) is shown. This cell is illustrated in supplementary movie S1. Scale bars, 1 µm. (C) Illustration of the asynchronous replication of the two alleles in one cell of the Early 2 cell line. In this cell, one allele replicates between the 70 and 75 min time points, and the second allele replicates between the 120 and 125 min time points. The alleles are therefore duplicated with a RD of 50 min. This cell is illustrated in supplementary movie S2. Scale bars, 1 µm.

### A tight synchrony of duplication of the two alleles is observed for the three loci replicated in the first part of S-phase

We ensured that the results obtained were representative of the entire cell population, by analyzing at least 50 cells corresponding to 100 chromosome duplication events for each of the six constructed cell lines. Replication timing (RT) of the two TetO-tagged alleles was determined in each cell (RT1 and RT2 refer to the allele that replicates first and to the one that replicates secondarily, respectively). Correlation degrees between RT1 and RT2 (Fig. 4), together with RD values (Fig. 5), allowed us to evaluate the stochastic nature of the replication timing program for the six studied loci.

**Figure 4.**
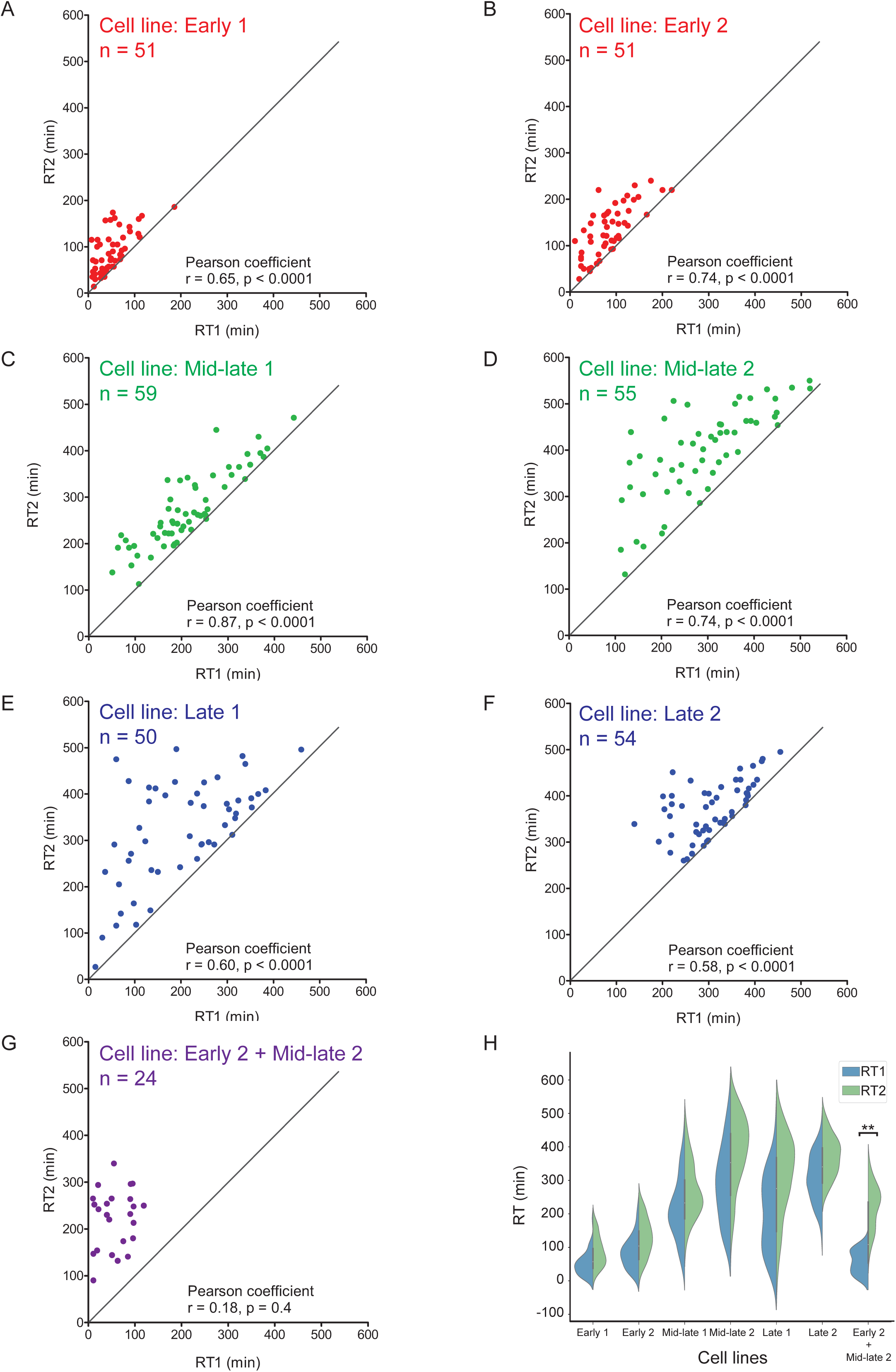
The stochasticity of replication timing increases with the progression of S phase. The timing of replication for the two alleles was determined for individual cells, by real-time imaging. (A-G) Replication timings (RT) of the two alleles in all cells analyzed are displayed on scatter plots. On the top of the graph the name of the cell line and the number of cells analyzed are indicated. RT1 is the RT of the allele that replicates first, and RT2 is the RT of the one that replicates secondarily. RT1 is on the *x*-axis and RT2 is on the *y*-axis. The Pearson coefficients are indicated. A gray line (y=x) is figured as an indicator of perfect synchronous replication of both alleles. (H) The probability density functions of the distributions of RT1 and RT2 for the seven cell lines are represented as violin plots. The violin plots use a kernel density estimation to show the distribution of values. In the six homozygous cell lines, the probability density functions of RT1 and RT2 are similar. In the Early 2 + Mid-late 2 cell line, the probability density functions RT1 and RT2 are significantly different (F-test, ** with *p*-value=0.0054).

**Figure 5.**
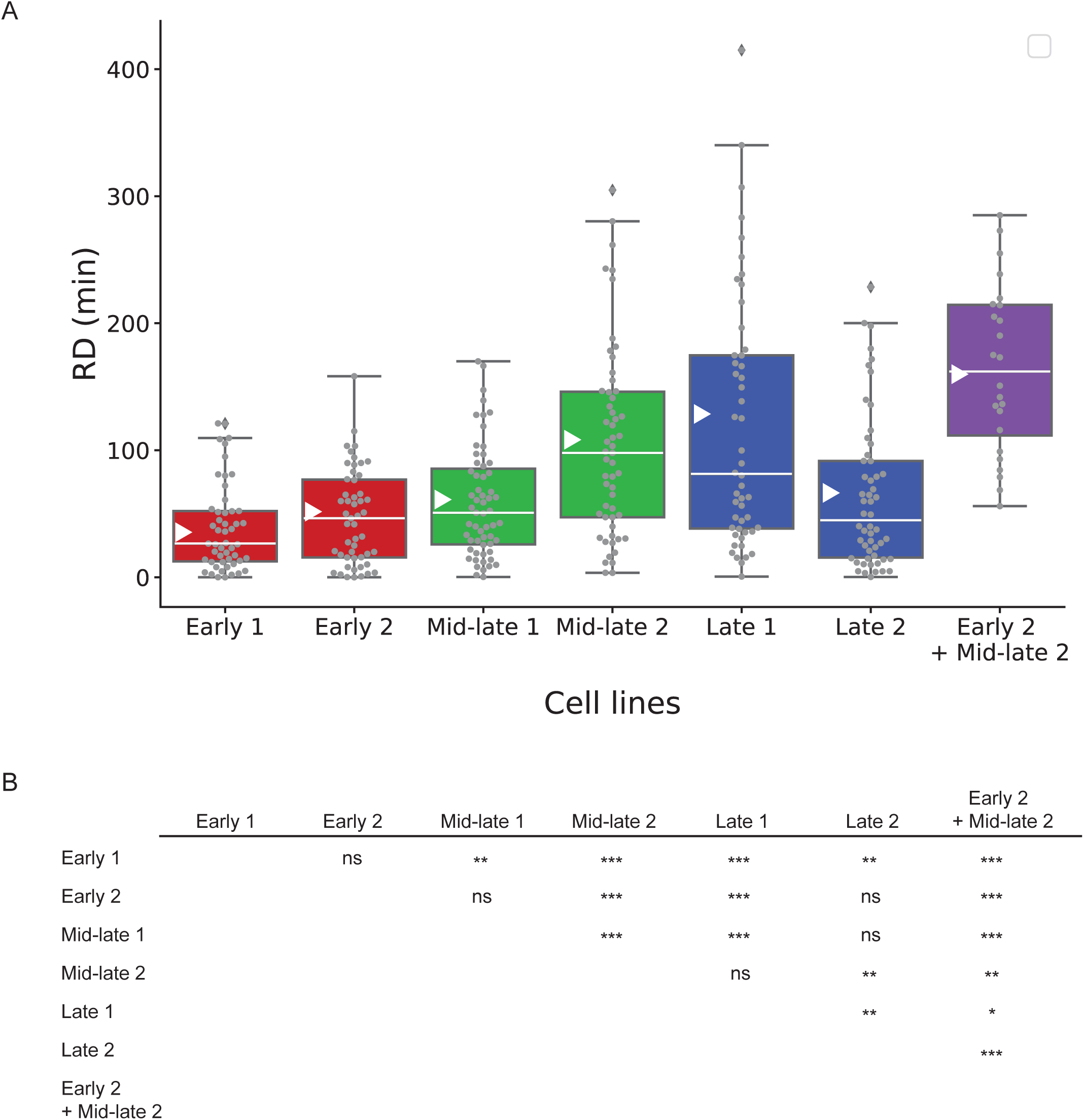
The replication delay between alleles varies as S-phase progresses. (A) Replication delays (RDs) for each of the seven cell lines are represented in box plots. Center white line is median; box limits are upper and lower quartiles; whiskers extend to points that lie within 1.5x interquartile range of the lower and upper quartiles; white triangle indicates mean value; dots are individual samples. (B) Comparisons of all the cell lines were performed with an unpaired t-test with Welch’s correction (*** with p-value<0.0001, two-tailed test).

In the Early 1, Early 2 and Mid-late 1 cell lines, the replication timings of the two alleles followed an almost linear distribution and were significantly correlated (Fig. 4A-C). The probability density functions of replication times were very similar for the two alleles at the same locus (Fig. 4H). In the Early 1 cell line, RD values ranged from 0 to 121 min with a median value of 26 min, and in the Early 2 cell line, RD values ranged from 0 to 158 min with a median value of 46 min (Fig. 5A). Both alleles were replicated in less than 60 min in 78% and 65% of the cells studied from Early 1 cell line and from Early 2 cell line, respectively. In the Mid-late 1 cell line, RD values ranged from 0 to 170 min with a median value of 51 min. RD values in the Mid-late 1 cell line are higher and significantly different from RD values in the Early 1 cell line, and they are similar to RD values in the Early 2 cell line (Fig. 5). Both alleles were replicated in less than 60 min in 56% of the cells from Mid-late 1 cell line. Replication timing of these three loci replicated in the first part of S-phase proved to be highly controlled in a narrow time window and is not much stochastic.

### Two out of three loci replicated in the second part of S-phase present a significant increase in stochasticity

The RT of the two alleles were also significantly correlated in Mid-late 2, Late 1 and Late 2 cell lines (Fig. 4D-F), with similar probability density functions of RT1 and RT2 (Fig. 4H), suggesting a tight control of their replication timing. In Mid-late 2 cell line, RD values ranged from 4 to 305 min with a median value of 98 min (Fig. 5A). Quite similarly, in Late 1 cell line, RD values ranged from 1 to 415 min, with a median value of 81 min (Fig. 5A). Both alleles were replicated in less than 60 min in 35% and 40% of the cells in the Mid-late2 and Late 1 cell lines, respectively. Distributions of RD values in these two cell lines are similar amongst themselves, but significantly different from the distributions of RD values from the three earliest loci (Fig. 5B). Some RDs obtained for the Late 1 cell line were even larger than those for the Mid-late 2 cell line, encompassing the entire S-phase in one of the 50 cells (RD=415 min) (Late 1 cell line, Cell: S17-cell 1 in Supplementary movie 3 and in Supplementary Fig. 5). Surprisingly, for a late locus, the first allele to be replicated in this cell was replicated at the start of S-phase, whereas the other allele was not replicated until the end of S-phase. However, as expected for a late locus, most of the alleles tended to replicate in the last part of S-phase, particularly in the cells with low RD values (Fig. 4E, Late 1 cell line, Cell: S12-cell 1 in Supplementary movie 4 and in Supplementary Fig. 5). This can also be seen for the probability density functions of individual alleles, with a large range of replication timings for the first allele to be replicated (RT1), covering most of S-phase, whereas the likelihood of the second allele being replicated (RT2) increases as S-phase progresses (Fig. 4H). The scattering of the replication of the first allele to be replicated over S-phase appeared to be even greater in the Late 1 cell line than in the Mid-late 2 cell line (Fig. 4H). We next analyzed RD values from 54 cells of the Late 2 cell line. In this cell line, RD values ranged from 0 to 229 min, with a median value of 44.5 min (Fig. 5A), and 57% of the cells replicate both alleles in less than 60 min. Surprisingly, the RD range and median value were lower than that of Mid-late 2 and Late 1 cell lines. As well, the probability density functions of RT1 and RT2 from Late 2 cell line were more restricted in the S-phase than those from Late 1 and Mid-late 2 cell lines (Fig. 4H). These observations indicated that the replication of this Late 2 locus seems to be restricted to a narrow time window at the end of the S-phase. RT profiles obtained with four S-phase sorted fractions showed that 68% of the replication events of the TetO array occurred during the last part of S-phase (S4 fraction) in the Late 2 cell line, compared to 53% in the Late 1 cell line (Fig. 2C) indicating that the Late 2 locus was replicated later than the Late 1 locus. We concluded that in this Late 2 locus, which replicates in the very last part of S-phase, the stochastic nature of replication timing is lower than in earlier replicated Mid-late 2 and Late 1 loci.

Overall, the measurement of RD values of six loci, whose replication timings are gradually distributed along the S-phase, shows that stochastic nature of the replication timing program is very low in the first part of S-phase, before it becomes more important in two loci replicated in the second part of S-phase. Surprisingly, stochasticity was found to be low in the very last part of S-phase for the studied locus that replicates last.

### Late 1 and Late 2 loci differently interact with nuclear lamina

To get some insight into what could explain the difference between the two late cell lines in terms of RD, we determined the nuclear position of the Late 1 and Late 2 loci, with respect to Lamin B1. Lamin B1 is one of the components of the nuclear lamina (Steensel & Belmont, 2017), a nuclear compartment strongly associated with late replication (Pope *et al*, 2014). We constitutively expressed the human Lamin B1 fused to DsRed1 (LMNB1-DsRed1) in the Late 1 and Late 2 cell lines. As expected, the DsRed1 fluorescence signal is detected at the nuclear periphery of all the cells (Fig. 6A). Distance between each of the two TetO/TetR-EGFP spots and LMNB1-DsRed1 was measured in 250 and 254 cells in Late 1 and Late 2 cell lines, respectively. The Late 2 locus interacted more frequently with the nuclear lamina than the Late 1 locus (median distance=0.19 µm and 0.57 µm for Late 2 and Late 1 cell lines, respectively) (Fig. 6B).

**Figure 6.**
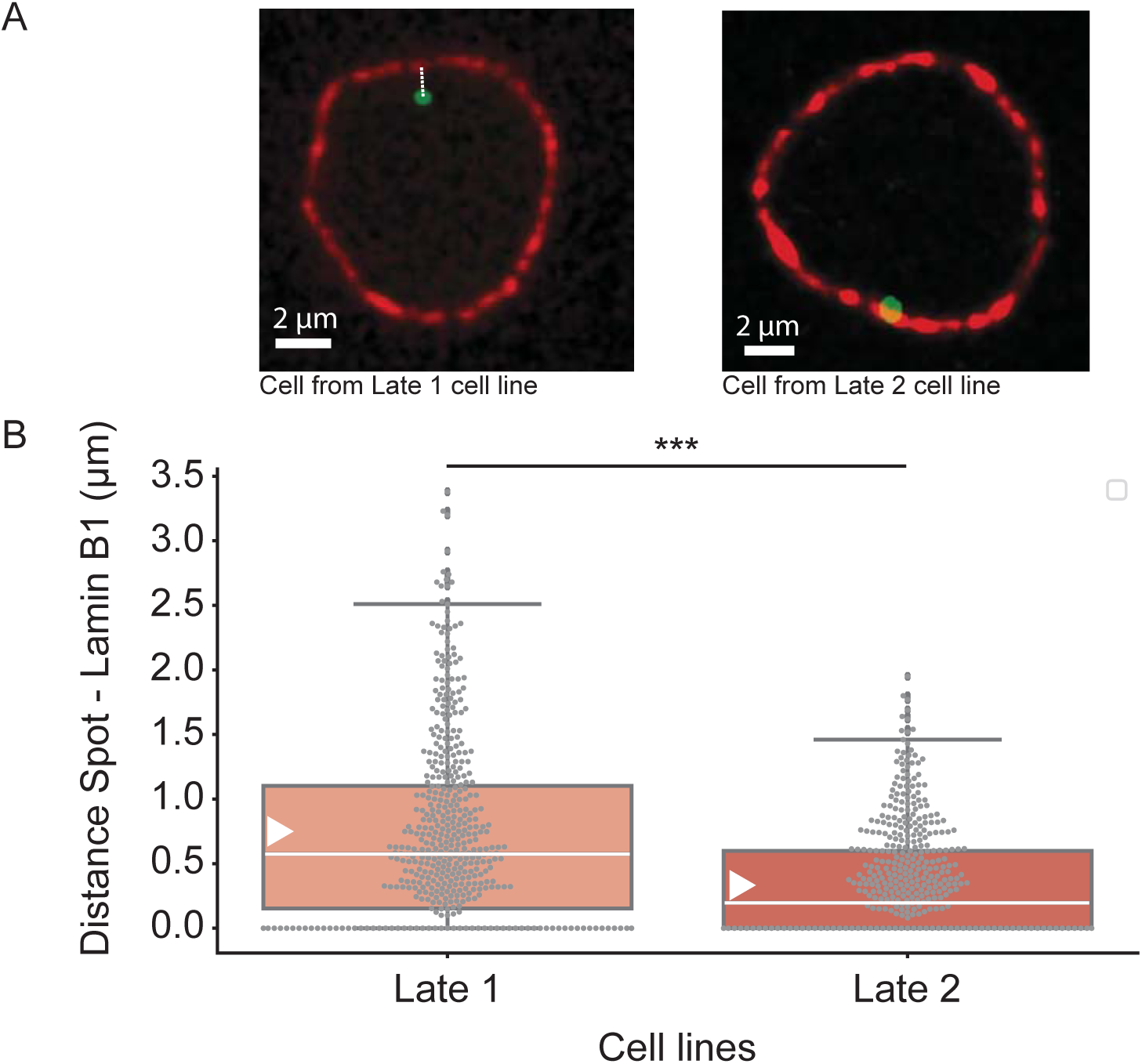
Localization of the TetO/TetR-EGFP spots with respect to Lamin B1. (A) The distances between the center of TetO/TetR-EGFP signal and the closest DsRed-LMNB1 signal (Lamin Associated Domain) were measured in a single focal plane (0.25 µm) of cells from Late 1 and Late 2 cell lines stably expressing LMNB1-DsRed. Scale bars, 2 µm. (B) TetO/TetR-EGFP spots in the Late 2 cell line interact significantly more frequently with Lamin B1 than in the Late 1 cell line. The results are presented in box plots. Center white line is median; box limits are upper and lower quartiles; whiskers extend to points that lie within 1.5x interquartile range of the lower and upper quartiles; white triangle indicates mean value; dots are individual samples. The numbers of nuclei analyzed were n=250 and n=254 in Late 1 cell line and Late 2 cell line, respectively. The number of TetO/TetR-EGFP analyzed spots were n=500 and n=508 for Late 1 cell line and Late 2 cell line, respectively, since each nuclei contains two TetO/TetR-EGFP spots in both cell lines. *P*-value was calculated using the unpaired t-test with Welch’s correction (*** with p-value<0.0001, two-tailed test).

### The single-cell approach can detect the asynchronous replication of an early locus and a mid-late locus

We investigated whether two independent loci with different replication timings in population-based assays were replicated in different time windows. We used our new single-cell approach to assess the replication timing of TetO arrays in individual cells for the Early 2 + Mid-late 2 cell line. We tracked and analyzed 24 cells. The replication timings of the two loci were found to be completely uncorrelated (Pearson coefficient *R*= 0.18, with *p*-value =0.4) (Fig. 4G). RD ranged from 56 min to 285 min, with a mean value of 164 min (Fig. 5A). Unlike the observations in the six homozygous cell lines, only one cell out of 24 cells (4%) replicated the two loci in less than 60 min. With this cell line, the probability density functions of RT1 and RT2 are significantly different, which is not the case for the six homozygous cell lines (Fig. 4H). Furthermore, the probability density function of RT1 was quite similar to that of the two alleles in the homozygous Early 2 cell line, whereas that for RT2 was broader, as for the two alleles of the Mid-late 2 cell line (Fig. 4H). We therefore hypothesized that the locus replicated first was the Early 2 locus. These observations indicate that the Early 2 and Mid-late 2 loci tend to replicate in separate time windows and further highlight the strong control of the timing program observed in the homozygous cell lines.

## Discussion

Our precise quantification of the replication delay between allelic loci in single cells revealed that alleles replicate synchronously, with a variation of the stochasticity of the replication timing program during the S phase. The six studied loci can be divided into two groups regarding the stochastic nature of their replication timing: (i) the three earliest loci (Early 1, Early 2 and Mid-late 1) together with the Late 2 locus and, (ii) the two late loci (Mid-late 2 and Late 1). Intriguingly, in the Late 2 cell line which replicates at the latest time point (Fig. 2C), alleles were replicated more synchronously than in the Late 1 cell line, breaking with the stochasticity increase observed in Mid-late 2 and Late 1 cell lines (Fig. 5A). Alleles were replicated quite asynchronously with RD values higher than 120 min in a significant proportion of cells found in the Mid-late 2 and Late 1 cell lines, 40 and 44% respectively. This high extent of stochasticity of replication timing has not been reported by previous allele-specific analyses. However, unlike our single-cells approach, previous allele-specific analyses were based on populations of millions of cells (Koren & McCarroll, 2014; Mukhopadhyay *et al*, 2014; Rivera-Mulia *et al*, 2018). This finding probably indicates a randomization of the stochastic nature of replication timing for the two alleles, with the first of the two alleles to be replicated differing between cells. Measurements of the mean replication timing of maternal and paternal alleles therefore leads to observations of synchronous replication over most of the genome (Mukhopadhyay *et al*, 2014). Our precise temporal measurements in individual cells indicated median time windows ranging from 26 min for an early region (Early 1 cell line) to 98 min for a late region (Mid-late 2 cell line). Koren *et al.* evaluated the structure of the replication timing program as a function of distance along the chromosome, by autocorrelation analyses. Their results suggest that the replication timing program may be less structured in late-replicating regions than in regions replicating earlier in S-phase (Koren & McCarroll, 2014).

The variable replication timing observed here for Mid-late 2 and Late 1 regions may be considered in light of the results obtained with haploid yeast cells. Indeed, modeling and measurements of origin efficiency and replication timing in *S. cerevisiae* have shown that earlier activation of an origin is associated with greater efficiency, with later activation of the origin associated with greater stochasticity of activation time (Hawkins *et al*, 2013; McGuffee *et al*, 2013; Moura *et al*, 2010; Yang *et al*, 2010). We think that comparison of haploid cells is a situation similar to the comparison of one chromosome with its homolog in single diploid cells from vertebrates. We found that replication of the Early 1, Early 2 and Mid-late 1 loci were highly synchronous on the two homologous chromosomes in most of cells, whereas the Mid-late 2 and Late 1 loci displayed less synchronous replication of the two alleles. Regarding the Late 2 locus, which displayed higher synchronous replication of both alleles, we hypothesized that its specific strong association with Lamin may underlie that situation (discussed below). In addition, a comparison of the replication timing of an early locus (Early 2) and a mid-late locus (Mid-late 2) within the same cells showed that the replication dynamics of these two loci were clearly separate in time, demonstrating accurate control of the progression of the timing program during S phase. Thus, replication dynamics, although controlled over time, appears to have stochastic properties that change during progression through the S phase, not only at the global scale of the cell population but also at the scale of individual cells.

Assuming that replication forks progress homogeneously, the synchronous replication of early, mid-late or late regions results from the synchronous activation of identical or closely spaced origins on the two chromosomes. Conversely, asynchronous replication of these regions, especially for late regions, may result from the asynchronous activation of different origins on the two chromosomes. The genome-wide detection of origins is mostly based on the identification of short nascent strands (SNSs) extracted from populations of millions of cells (Besnard *et al*, 2012; Cadoret *et al*, 2008; Cayrou *et al*, 2011; Langley *et al*, 2016; Picard *et al*, 2014; Sequeira-Mendes *et al*, 2009). Origin mapping in various species has shown that there are more origins in early-than in late-replicating regions, and that the origins in early-replicating regions are more efficient (Prioleau & MacAlpine, 2016). We hypothesize that, in early regions, efficient, abundant and physically grouped origins are synchronously activated on a large proportion of chromosomes, resulting in overall synchronous replication in individual cells. By contrast, in late regions, less efficient and more physically dispersed origins are activated with various degrees of synchrony, differing between chromosomes, resulting in less synchronous replication overall. The difference in synchrony between early-replicating regions (Early 1, Early 2 and Mid-late 1 loci) and late-replicating regions (Mid-late 2 and Late 1) observed here may, therefore, be directly related to origin abundance and strength, which were found to decrease from early-to late-replicating regions. A crucial step in replication initiation is the recruitment of firing factors to licensed origins, triggering replisome assembly and bidirectional replication. Limiting concentrations of these factors in the nucleus have been shown to be related to the correct course of the timing program (Douglas & Diffley, 2012; Mantiero *et al*, 2011; Tanaka *et al*, 2011; Wong *et al*, 2011). Efficient origins might have a high affinity for these limiting factors and recruit them as soon as S-phase starts, leading to their early and efficient firing on most chromosomes, thereby favoring synchrony or low stochasticity. Once replication is underway, limiting factors can be recycled to origins with lower affinities, until the entire replication program is completed. Activation time gradually becomes more stochastic as the affinity of the origin for limiting factors decreases. Here, such a situation in the Early 1, Early 2 and Mid-Late 1 cell lines may underlie the synchronous replication of the two alleles restricted to the first part of S-phase in all cells. In the Mid-late 2 cell line, this situation results in less synchronous replication in the second part of S-phase. The replication time of the Late 1 locus is even more variable, with one allele replicating in the first part of S-phase and the other replicated at the end of S-phase in some cells. We can speculate that, in a late region, which has origins of low efficiency with a low affinity for limiting factors, replication is initiated at different time points on the two chromosomes. As previously suggested, the probability of a late domain being replicated probably increases as S phase progresses (Yang *et al*, 2010). Consistent with this view, we found that a reasonably large proportion of cells (40%) replicate both alleles quite synchronously (RD≤60 min), mostly in the second part of S-phase. The synchronous replication in these cells may have arisen from the essentially synchronous activation of the same of nearby origins in both alleles. These origins may be among the late origins firing efficiently in narrow time windows in regions that have not yet been passively replicated, as described by McGuffee *et al*. (McGuffee *et al*, 2013). Likewise, replication of the Late 2 locus, which replication timing revealed to be more strictly controlled and displayed less stochasticity, may be initiated by such origins. We showed that this locus is tightly associated with nuclear lamina at the nuclear periphery. The Lamin-associated domains (LADs) have been reported to overlap with regions that replicate late during S-phase (Steensel & Belmont, 2017; Pope *et al*, 2014). A recent paper addressed genome-wide the causal relationships between nuclear lamina association and gene expression (Leemans *et al*, 2018). The authors observed important variation in reporter expression levels within LADs implying that they are heterogeneous structures. Moreover, the chromatin feature that was the most predictive of reporter expression in LADs was the frequency contact with the nuclear lamina. It suggests that reporters are repressed more efficiently when they are inserted in regions that are more stably associated with the nuclear lamina. Similarly, the strong interaction of the Late 2 locus with lamin B1 might have a robust repressive effect on origin firing until the end of S-phase. We speculate that replication of such late replicating domains occurred quickly and efficiently in the very last part of S-phase, as previously observed by single-molecule analyses (Guilbaud *et al*, 2011). Due to their particular nuclear localization, they may be inaccessible to the limiting firing factors before the end of S-phase thus efficiently preventing early firing. In very late S-phase, when most of the genome has already been replicated, all these factors can be concentrated in the nuclear lamina compartment leading to a very efficient firing of numerous origins, even though these origins are not as strong as those found in early replicating regions.

## Methods

### Plasmid construction

The coding sequence of TetR was amplified with the Herculase II Fusion DNA Polymerase (Agilent) from the pLAU53 vector (gift from DJ Sherratt), with the addition of two NLS sequences. The amplicon was inserted in-frame with the EGFP sequence in the pEGFPN1 vector (Clontech) and the resulting plasmid was named pEGFP-TetR. The MultiSite Gateway Pro Kit (Invitrogen) was used to generate targeting plasmids with the TetO array. Briefly, two homologous arms flanking the insertion site were amplified by PCR from DT40 genomic DNA and inserted into pDONR P1-P5 (5’ arm) and pDONR P3-P2 (3’ arm). An array of 4.4 kb with 120 TetO repeats was amplified by PCR from the pLAU44 vector (gift from DJ Sherratt) and inserted into pDONOR P5-P4. The primers used are described in Table S1. The blasticidin resistance cassette was previously inserted into pDONR P4r-P3r (Hassan-Zadeh *et al*, 2012). The final complete plasmids were constructed by ordered assembly of the 5’-arm, TetO array, blasticidin resistance cassette and 3’-arm.

### Cell culture and cell line construction

The DT40 cell lines were cultured in RPMI 1640 with Glutamax supplemented with 10% FBS, 1% chicken serum, 0.1 mM β-mercaptoethanol, and penicillin plus streptomycin, at 37°C, under an atmosphere containing 5% CO_2_. A first cell line expressing TetR-EGFP was established by co-electroporation with the pEGFP-TetR and pLoxBsr (Arakawa *et al*, 2001) plasmids. Electroporation and clone selection were as previously described (Hassan-Zadeh *et al*, 2012). One independent clone, DT40 TetR2, was selected on the basis of GFP fluorescence intensity in the cell nuclei and was used for the successive insertion of TetO arrays.

The TetO array was stably inserted into the first allele by electroporating 10^7^ DT40 TetR2 cells with 35 µg linearized final complete plasmid with TetO. Independent drug-selected clones were verified by PCR amplification with a primer pair combining one primer binding within the insert and another binding just downstream of the 3’ arm sequence (Bls-Forward and 3’-site-Reverse, Supplementary Fig. 1). The blasticidin resistance cassette is flanked with mutant LoxP RE and LoxP LE sites (Arakawa *et al*, 2001). This cassette was excised by culturing cells for 24 h in normal growth medium supplemented with 4-hydroxyl-tamoxifen (2 µM 4-OH tamoxifen), which induced the translocation to the nucleus of the Cre recombinase expressed in DT40 cells. Recombination at the two LoxP sites led to the excision of the blasticidin resistance cassette. Independent clones were obtained from single-cell limiting dilution cultures in normal medium. These clones were verified by PCR amplification with a primer pair flanking the blasticidin cassette (TetO-Forward and 3’-arm-Reverse, Supplementary Fig. 1), and by evaluation of their sensitivity to the drug. One of the selected clones was chosen at random for the insertion of a TetO array into the second allele. The transfection and selection protocol was identical to that for modification of the first allele, except for the last step (checking for blasticidin resistance cassette excision). As the DNA sequences of the two alleles were indistinguishable at this step, verification was based exclusively on sensitivity to the drug. Construction of homozygous cell lines proved to be more challenging for late loci. For the establishment of the Late 1 cell line, we indeed got very few clones with site-specific insertion of TetO array on the second chromosome. Construction of the Late 2 cell line was even more difficult and despite multiple experimental attempts we could not obtain any clone having a homozygous insertion of TetO arrays (276 negative clones issued from 9 electroporation experiments). We overcame this difficulty by inserting the second TetO array on the homologous chromosome in a site nearby the first site of insertion (5.6 kb upstream). Such a short distance between these two sites should not impact significantly on the RD between the two TetO arrays since average fork speed in DT40 is ~1.2 kb/min (Petermann *et al*, 2006). We therefore assumed that the Late 2 cell line constructed in this way mimics a genuine homozygous cell line. Finally, the two expected fluorescent spots in the nucleus of each cell were visualized by confocal microscopy (see below). One clone was chosen at random for further experiments.

We constructed two additional cell lines expressing the Lamin B1 fused to DsRed protein (LMNB1-DsRed). The Late 1 and Late 2 cell lines were co-electroporated with the plasmids pDsRed1-LMNB1 (gift from B. Buendia (Delbarre *et al*, 2006)) and pLoxBsr. For each cell line, one independent drug-resistant clone was selected on the basis of DsRed fluorescence intensity in the cell nuclei.

### Cell-cycle synchronization by centrifugal elutriation

Centrifugal elutriation was performed as previously described (Gillespie and Henriques 2006), in a Beckman Coulter Avanti^®^ J-26 XP with a JE-5.0 rotor and the 4 ml chamber. Briefly, 10^8^ asynchronous cells were harvested and resuspended in 100 ml of elutriation buffer (1X DPBS, 1% FBS, 1 mM EDTA). Cells were loaded into the chamber at room temperature, with a constant flow rate (40 ml/min) and a constant rotor speed (3800 rpm). After 15 minutes, cells with different sedimentation velocities were in equilibrium at different radial positions in the chamber. Seven 150 ml fractions (F1 to F7) were then collected by slowing the rotor speed, in 200 rpm steps, to 2400 rpm. Cells from with the F2 fraction containing G1 cells were harvested and resuspended in normal growth medium at a density of 10^6^ cells/ml. These cells were then used in live-cell imaging experiments.

### Timing assays

Timing assays were performed as previously described (Hassan-Zadeh *et al*, 2012). Briefly, BrdU (3 mM) was incubated with 10^7^ asynchronously growing cells for one hour. Cells were fixed in 70% ethanol and subjected to FACS on a BD INFLUX 500 (BD BioSciences), on the basis of DNA content, as assessed by propidium iodide staining. Four S-phase fractions (S1 to S4) of 50,000 cells each were collected. DNA was extracted from each fraction and sonicated to generate fragments of 500 to 1000 bp. DNA fragments that had incorporated BrdU were immunoprecipitated with an antibody against BrdU (BD Biosciences) and finally resuspended in TE buffer. The relative amounts of DNA in the four fractions were estimated by the qPCR quantification of mitochondrial DNA (Prime Pro 48 instrument from Techne). Relative quantification of specific DNA fragments in each of the four fractions was performed with qPCR using specific primer pairs (Table S1), with adjustment for mitochondrial DNA content.

### Live-cell imaging

G1 cells from elutriated fraction F2 were cultured at a density of 2 × 10^6^ cells/ml in a µ-Slide VI 0.4 ibiTreat (Biovalley) with normal growth medium (except that we used RPMI without phenol red) under an inverted confocal microscope (Leica DMI 6000 equipped with a Yokogawa spinning disk, with a QuantEM EMCCD camera). Cells were cultured at 37°C, under an atmosphere containing 5% CO_2_, throughout the experiment. All images were acquired under excitation with a 491 nm laser, with a Plan APO 100 X/NA 1.4 objective, an exposure time of 40 ms and 28 z-stacks (0.5 µm), with MetaMorph software (Molecular Devices). Phototoxicity was minimized by ensuring that the total exposure time did not exceed 120 s. The total acquisition periods were 250 minutes for the Early 2 cell line, 360 minutes for the Early 2 + Mid-late 2 cell line, and 500 minutes for the other cell lines, with images obtained at five-minute intervals.

### Image analyses and graph construction

We first removed the background from each image with ImageJ software (subtract background function with a rolling ball radius of 18 pixels). All images were then analyzed with Imaris software (versions 7.7 and 8.3.1, BitPlane, Oxford Instruments) software for 3D/4D visualization and analysis. Cells were individually tracked over the entire acquisition period. At each time point, the two nuclear spots were detected on the 3D image with the spot module of Imaris, and manually adjusted if necessary, and their fluorescence intensity was measured. Each spot was individually tracked over time with the ImarisTrack option, and manually corrected if necessary (supplementary Fig. 4). Data were then analyzed with a dedicated algorithm that determines RT values, calculates RD value, corrects photobleaching and constructs intensity graph. Briefly, the RT was identified through a change detection algorithm. This algorithm modeled the data as a unit step function, and accounted for the fact that the observations were corrupted by a linear drift due to photobleaching. Two linear regression models were computed before and after each possible change point and the best change point, *i.e*. the RT value, was automatically selected as the one with minimum mean square residuals. As a by-product, this approach identified the slope of amplitude declination and allowed correction of the photobleaching. Finally, we computed the parameters of a sigmoid function that fits the data. For a more convenient visualization of graph representations for the seven cell lines, times were adjusted by adding 70 minutes and 10 minutes to the coordinates of the Mid-late 2 cell line and the Late 1 cell line, respectively, because cells of these two cell lines have been cultured 130 minutes (Mid-late 2 cell line) and 70 minutes (Late 1 cell line) before the start of acquisition, while the other cell lines have been cultured 60 min before the start of acquisition. Box plots and violin plots were constructed with Python tools.

## Supporting information

## Data availability

All relevant data are available from the authors.

## Author Contributions

M.-N.P. and B.D. conceived, designed and interpreted the experiments. B.D., S.C., J.-F.B. E.H. and N.B. performed the experiments. M.-N.P. and B.D. wrote the manuscript.

## Acknowledgments

The authors thank the members of the laboratory of M-N.P. for useful insights and discussions. We acknowledge the ImagoSeine core facility of the Institut Jacques Monod, member of IBiSA and France-BioImaging (ANR-10-INBS-04) infrastructures, in particular Xavier Baudin for help with photonic microscopy and Vincent Contremoulins for advice on image analysis with Imaris software. We thank Dr. David .J. Sherrat for providing the pLAU53 and pLAU44 plasmids, and Brigitte Buendia for providing pLaminB1-DsRed1 plasmid. This work was supported by grants from the Association pour la Recherche sur le Cancer (Equipe Labellisée), the Agence Nationale pour la Recherche (ANR-15-CE12-0004-01), and the IdEx Paris Sorbonne. B.D. and M-N.P. are supported by Inserm.

