## Supplementary material for "Replication dynamics of individual loci in single living cells reveal variation of stochasticity"

**Duriez et.al**

**Supplementary Information**

**Supplementary Figure Legends**

**Supplementary Figure 1.** UCSC genome browser visualization (Assembly Gallus_gallus-5.0/galGal5 Dec 2015) of 100 kb windows centered on the six sites (arrow) where TetO arrays were inserted. Tracks showing CG percentage, annotated genes and CpG Islands are represented.

**Supplementary Figure 2.** Construction of cell lines with the insertion of TetO arrays into both alleles of a locus. (A) Schematic representation of the insertion of TetO arrays into both alleles of the selected locus. The TetO array, together with the blasticidin resistance gene (Bls), was integrated into the selected locus by homologous recombination involving the accompanying 5' and 3' arm sequences, which flank the insertion site. We checked the targeted insertion of the TetO array into the first chromosome, using the [Bls Forward]/[3' site Reverse] primer pair. The blasticidin resistance gene was then excised. Correct excision was verified by amplifying with the [TetO Forward]/[3' arm Reverse] primer pair and by determining the sensitivity of the cultured cells to the drug. A second TetO array was then inserted into the second chromosome, and its targeted insertion was checked with the [Bls Forward]/[3' site Reverse] primer pair. Finally, the blasticidin resistance gene was excised and correct excision was verified by checking the sensitivity of the cultured cells to the drug. Insertion sites: orange line, homology arms: thick blue lines, TetO array: red hatched box, blasticidin resistance cassette: green box, primer: thin arrow. (B) PCR amplification products were subjected to electrophoresis in an agarose gel. Names of the cell lines were indicated on the top of the gels, and primer pairs used for PCR amplifications are indicated on the bottom of the gels. In the Late 2 cell line, as two different insertion sites (site 1 and site 2) were used (see Methods), the insertion on the second allele in site 2 was checked with primers specific to site 2. The 1 kb Plus DNA ladder was used (L).

**Supplementary Figure 3.** Insertion of a TetO array into one allele does not affect its replication timing in heterozygous cell lines. (A) Schematic illustration of the experiment. Asynchronous cells were pulse-labeled with BrdU for one hour and fixed before sorting by FACS into four S-phase fractions (S1 to S4). Nascent DNA strands were purified by immunoprecipitation for BrdU. Quantitative PCR analyses were performed on the nascent DNA strands purified from each of the four S-phase fractions. Three primer pairs were used: the primer pair “With TetO” is specific for the allele with a TetO array (red arrows), the primer pair “Without” is specific for the wild-type allele (orange arrows) and the primer pair “Both” amplifies both alleles (blue arrows). Quantification with the “Both” primer pair should give an average of the amplification achieved with the two allele-specific primer pairs and was used as a control in the quantitative PCR experiment. The mode of calculation for shift in replication timing between the two alleles is indicated (ΔL: Late shift, ΔE: Early shift). (B-G) Insertion of the TetO array into the six loci does not change replication timing, even though insertion of the TetO array into the Mid-late 2 region slightly affects replication timing, shifting replication to later time points ($\Delta L$=6.91). However, this timing shift is not significant relative to the results of our previous analyses [11,16]. Quantitative PCR was performed on three independent loci representative of early- (Controls 1 and 2) and late- (Control 3) replicating regions. Quantitative PCR on these three control regions was used to evaluate and validate FACS results (the results are shown on the right-hand part of the histograms). Error bars indicate standard deviation for qPCR duplicates.

**Supplementary Figure 4.** Image analyses**.**

During the time-lapse, serial Z-plane frames (28 frames with a depth of 0.5 µm) were acquired at different positions every five minutes. (A) Background was subtracted from these images with the *subtract background* function of ImageJ software (the *Rolling Ball radius* parameter was fixed to 18 pixels). (B) Each z-stack acquisition was then analyzed as a 3D image with Imaris software (Imaris 7.7 and 8.3.1, Bitplane, Oxford intruments). Analyses were performed on individual cells through the establishment of a region of interest (ROI) which includes only the cell to be analyzed (cube in dotted lines, illustrated here for the cell 4 in position S19 for the Early 2 cell line). (C) The two TetO arrays bound to TetR-EGFP were visualized as two fluorescent bright spots within the nucleus. (D) Measurement of the fluorescence intensity of the spots was performed with the Spot function of Imaris software. Briefly, an ellipsoid (dimension: x,y=1.49 µm and z=3 µm) was centered on the maximum of intensity of the spot, and the sum of the fluorescence intensity within that ellipsoid was measured as the spot intensity. (E) Each spot was tracked over time with the spot tracking function of Imaris software, and manually adjusted if necessary. (F) Replication of both alleles in cell 4 from position S19 of the Early 2 cell line. On four successive images, the fluorescence intensity of each allele is indicated. Allele 1 was replicated between 55 and 60 min, and allele 2 between 60 and 65 min. (G) Left graph, variation of the fluorescence intensity of the two spots was reported on a graph. Right graph, we have developed a dedicated algorithm that fit the data on sigmoid curves and determined the replication time (RT) for each allele, thereby enabling calculation of the RD value.

**Supplementary Figure 5.** Fluorescence intensity variation curves for all analyzed cells. The variation of the fluorescence intensity of the two TetO/TetR-EGFP spots through the time-lapse is figured on a graph. Identification of each cell refers to the cell line, the position under the microscope (S-) and the number of the cell (Cell -). The curve of the allele that replicates first is in blue, and the curve of the other allele is in green. Photobleaching has been corrected. Replication times are indicated with blue and green dotted lines, and RD value is indicated on the right part of the graph.

**Supplementary Movies**

**Supplementary Movie 1** Cell from the Early 2 cell line (S11, Cell 1) in which the two alleles are replicated at the same time (RT1 = RT2 = 93 min). This cell was imaged over 250 minutes, at five-minute time intervals, in the GFP channel. This cell is the same as that shown in Figure 3b.

**Supplementary Movie 2** Cell from the Early 2 cell line (S11, Cell 5) in which the two alleles are replicated 50 minutes apart (RT1= 72 min and RT2= 122 min). This cell was imaged over 250 minutes, at five-minute intervals, in the GFP channel. This cell is the same as that shown in Figure 3c.

**Supplementary Movie 3** Cell from the Late 1 cell line (S17, Cell 1) in which the two alleles are replicated 415 minutes apart (RT1= 50 min and RT2= 465 min). This cell was imaged over 500 minutes, at five-minute intervals, in the GFP channel. The variation of GFP spot intensities is shown in Supplementary Fig.5.

**Supplementary Movie 4** Cell from the Late 1 cell line (S12, Cell 1) in which the two alleles are replicated at the same time (RT1= 301 min and RT2= 302 min). This cell was imaged over 500 minutes, at five-minute intervals, in the GFP channel. The variation of GFP spot intensities is shown in Supplementary Fig. 5.

**Supplementary Movie 5** Replication of the two alleles followed by cell division. Cell from the Mid-late 2 cell line (S25, Cell 2) in which the two alleles are replicated 31 minutes apart (RT1= 91 min and RT2= 122 min) and the nucleus divides 280 minutes later. This cell was imaged over 500 minutes, at five-minute intervals, in the GFP channel. The variation of GFP spot intensities is shown in Supplementary Fig. 5.

**Supplementary Table**

**Supplementary Table 1**

Primer sequences

Supplementary Figure 1

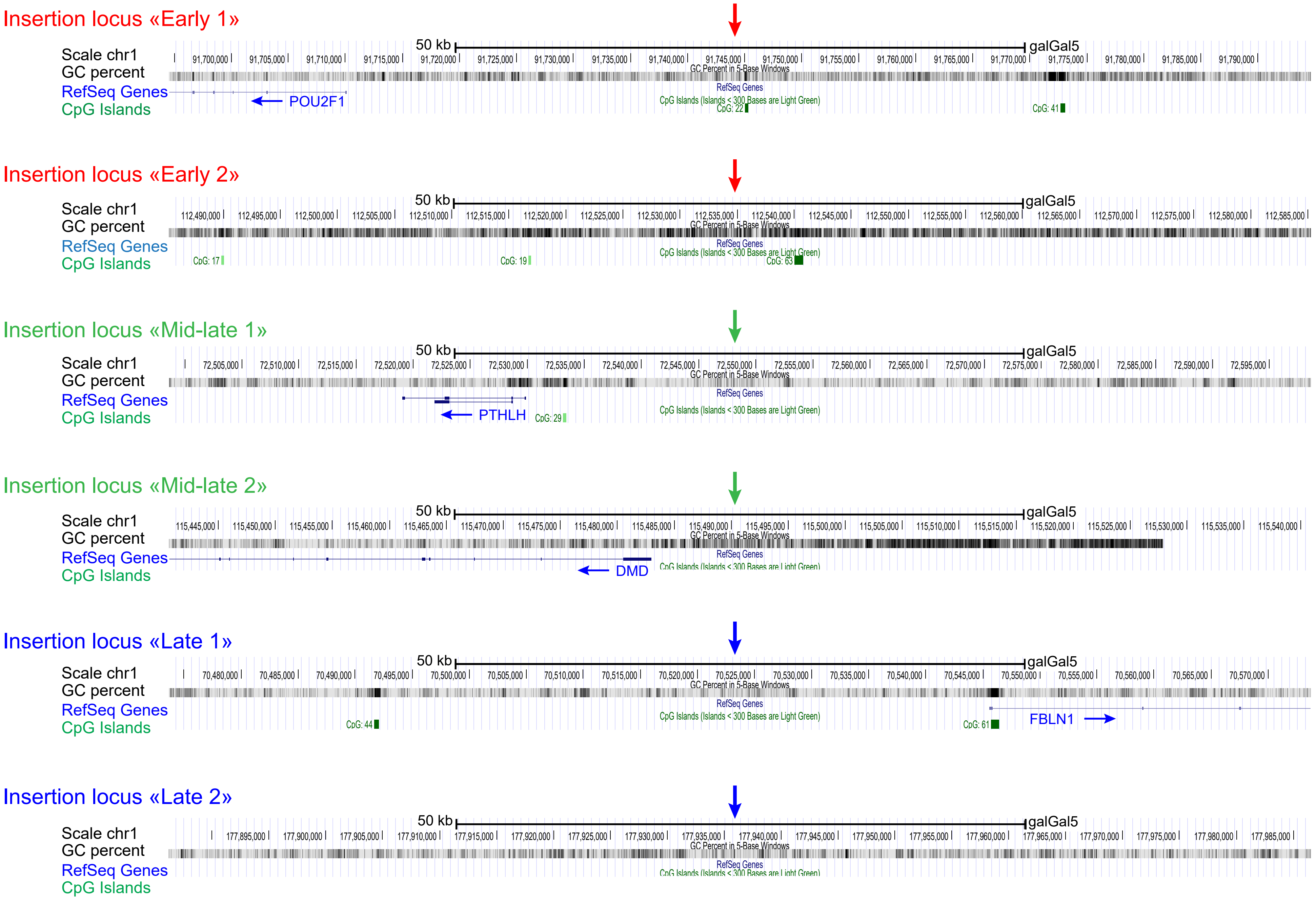

Supplementary Figure 2

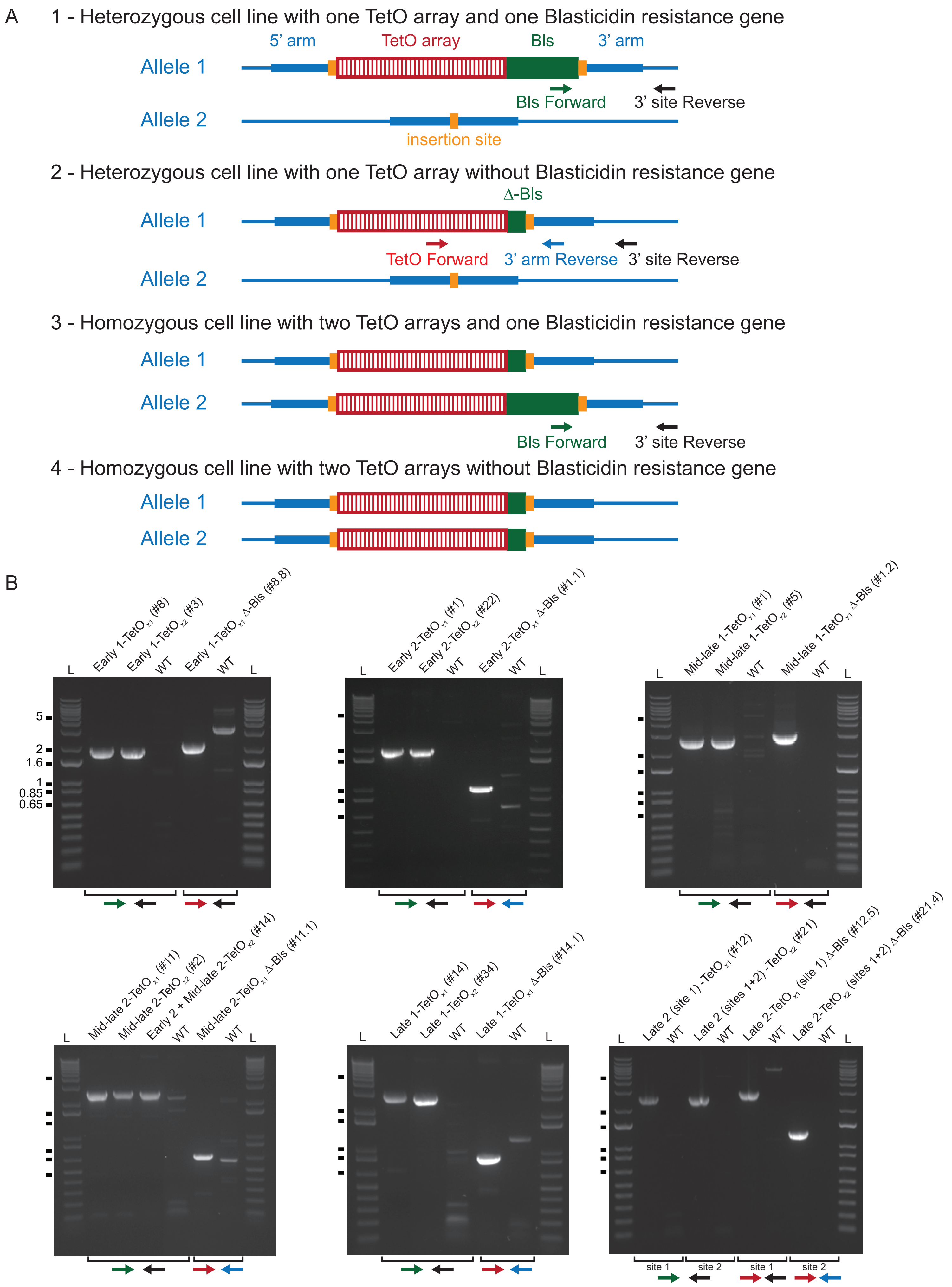

Supplementary Figure 3

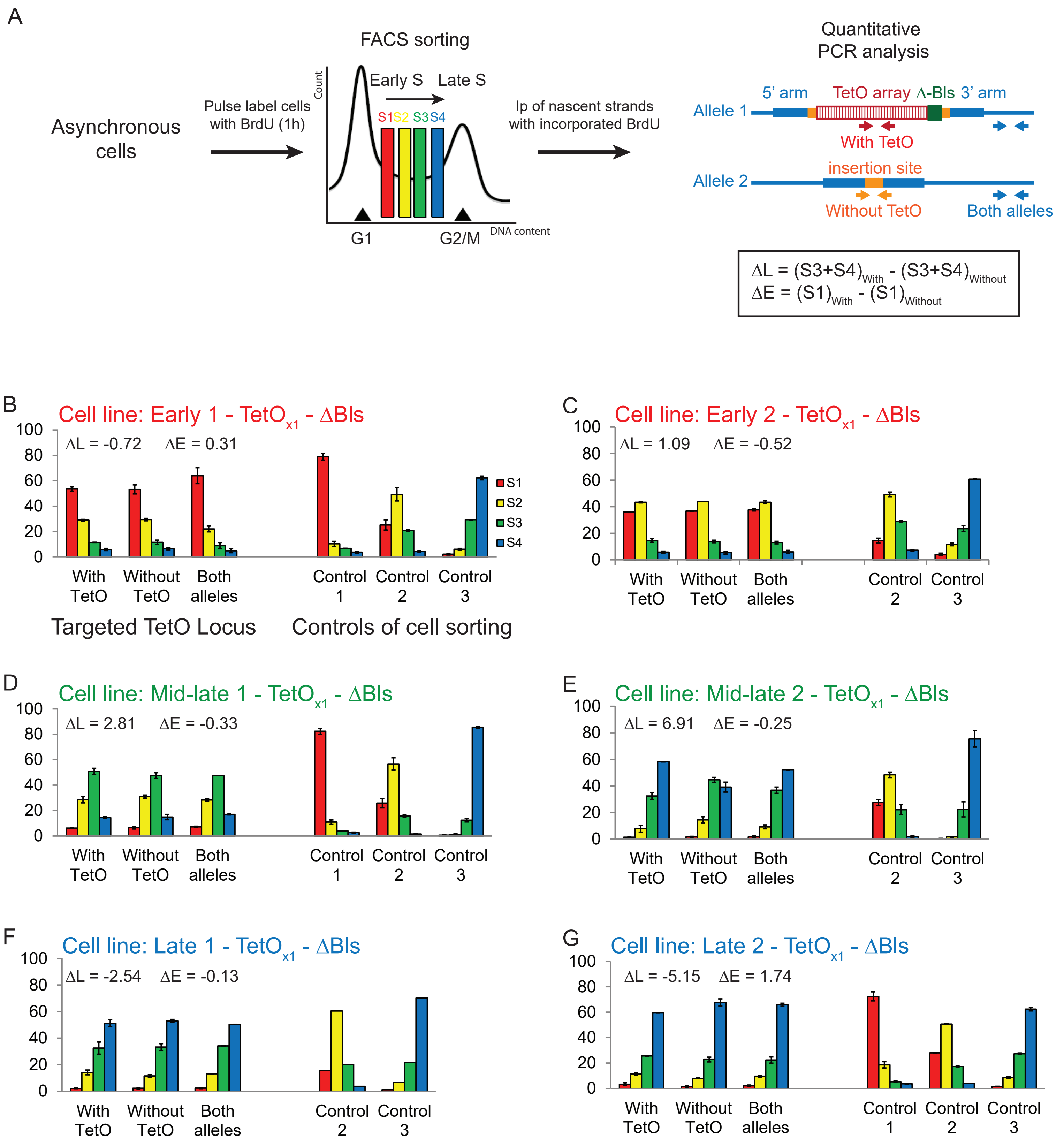

### Supplementary Figure 4

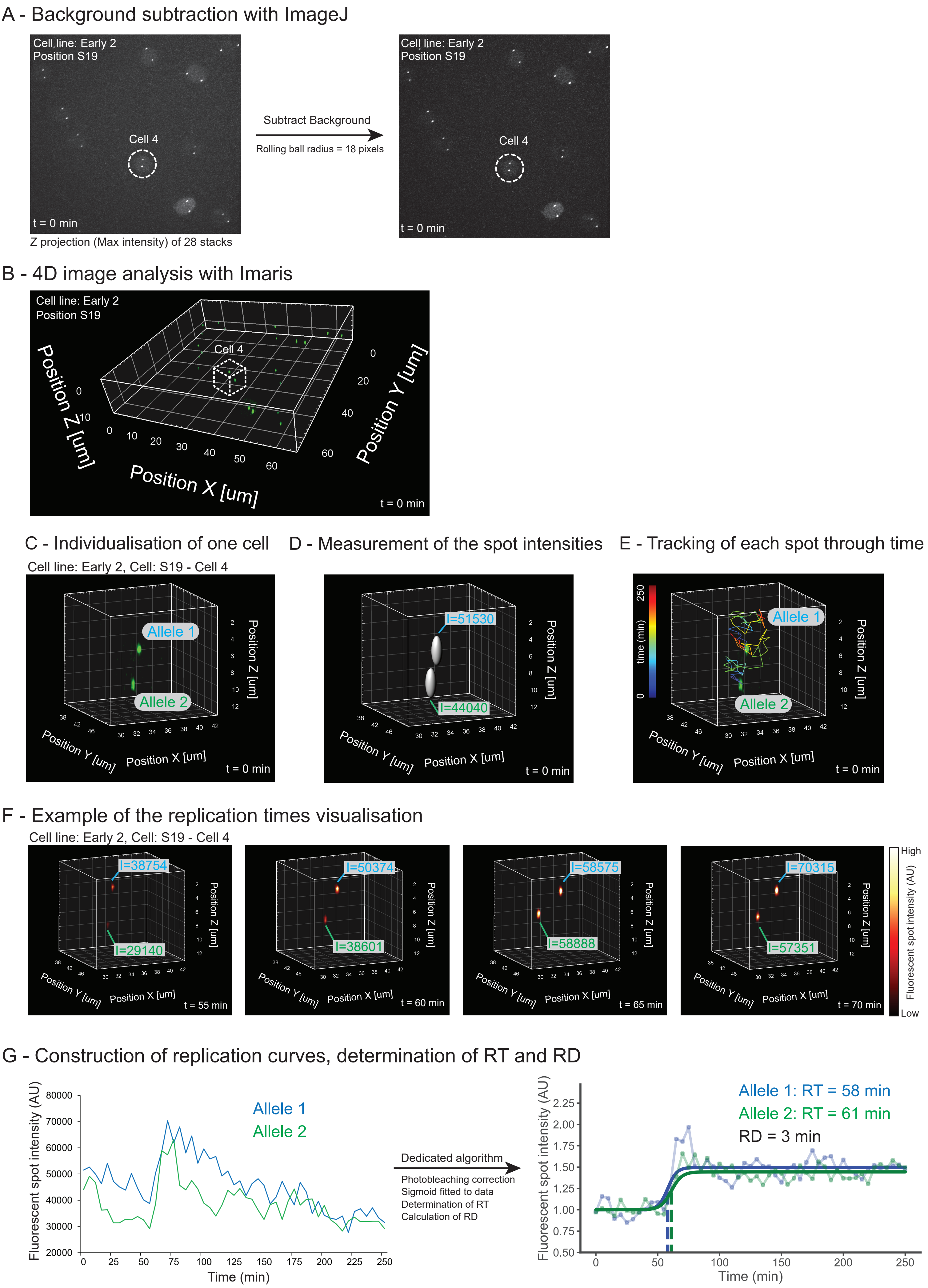

Supplementary figure 5

### 1 Early 1

Cell line: Early 1, cell: S1 cell 1

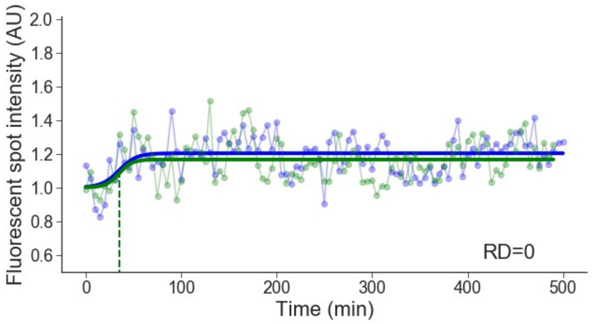


Cell line: Early 1, cell: S1 cell 2

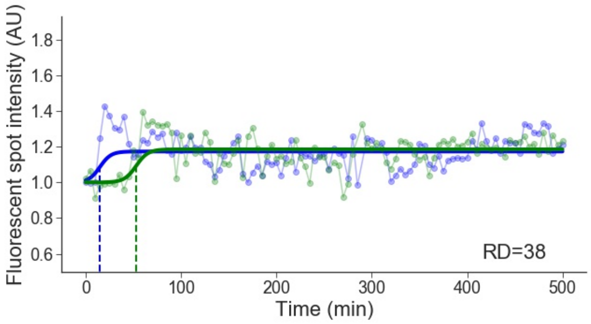


Cell line: Early 1, cell: S1 cell 3

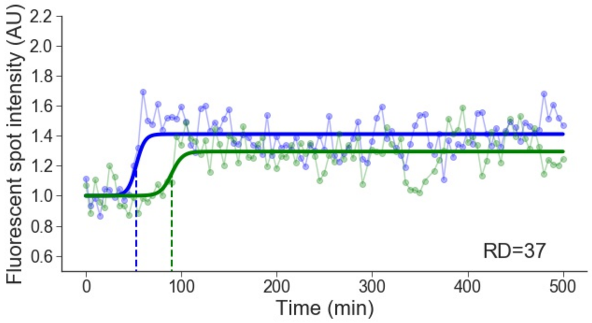


Cell line: Early 1, cell: S2 cell 2

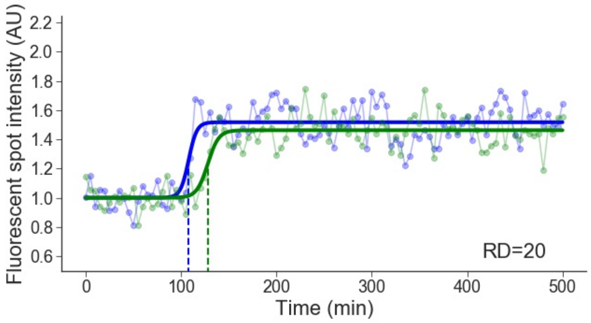


Cell line: Early 1, cell: S2 cell 3

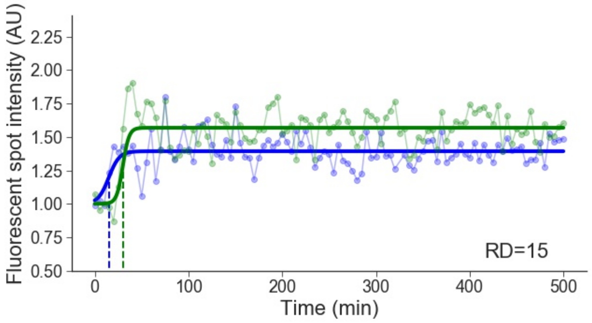


Cell line: Early 1, cell: S3 cell 1

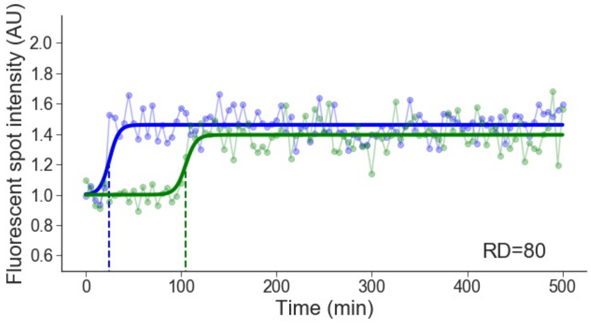


Cell line: Early 1, cell: S3 cell 3

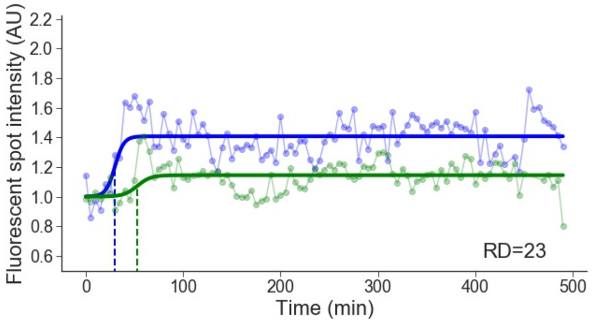


Cell line: Early 1, cell: S4 cell 1

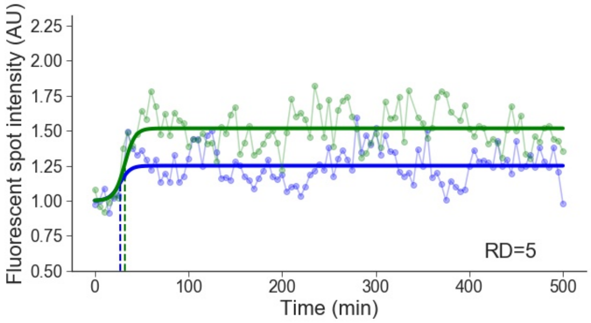

Cell line: Early 1, cell: S4 cell 2

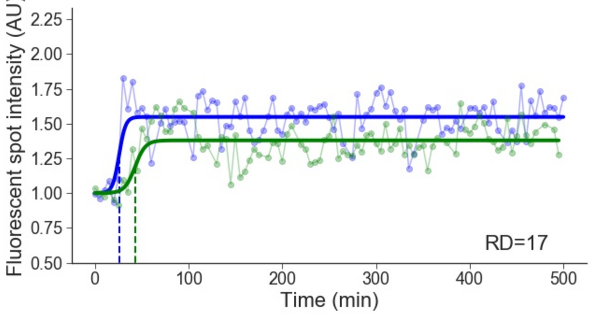


Cell line: Early 1, cell: S4 cell 3

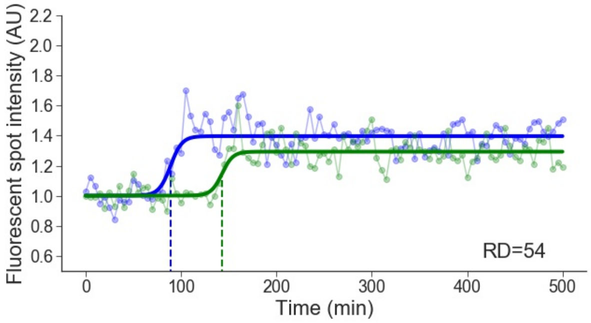


Cell line: Early 1, cell: S4 cell 4

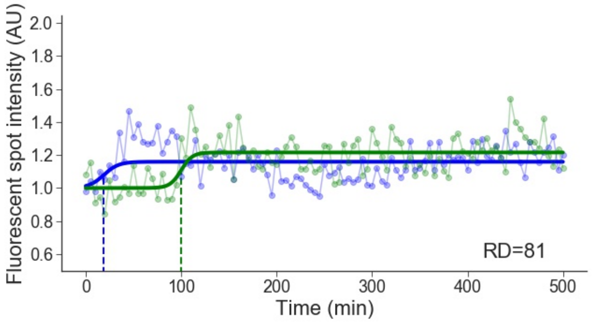


Cell line: Early 1, cell: S5 cell 2

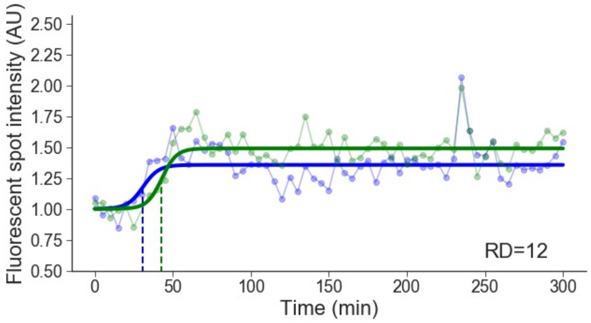


Cell line: Early 1, cell: S5 cell 3

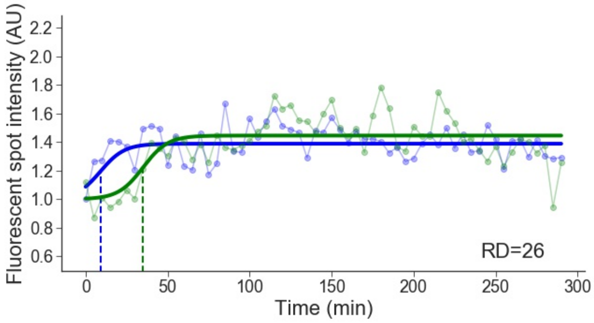


Cell line: Early 1, cell: S5 cell 4

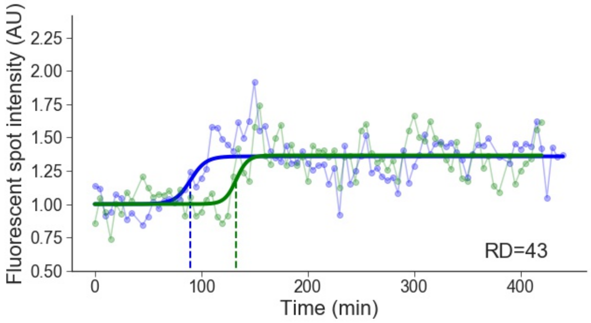


Cell line: Early 1, cell: S6 cell 1

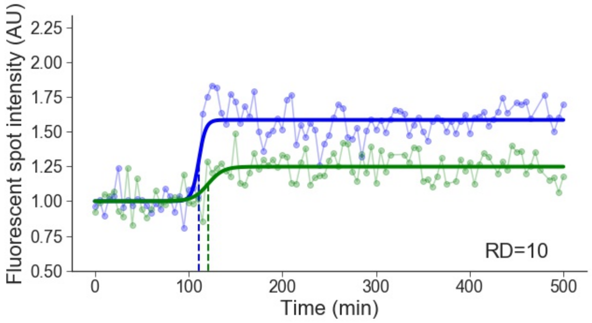


Cell line: Early 1, cell: S6 cell 3

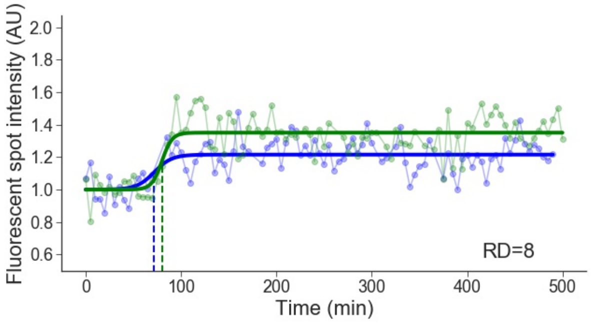

Cell line: Early 1, cell: S7 cell 1

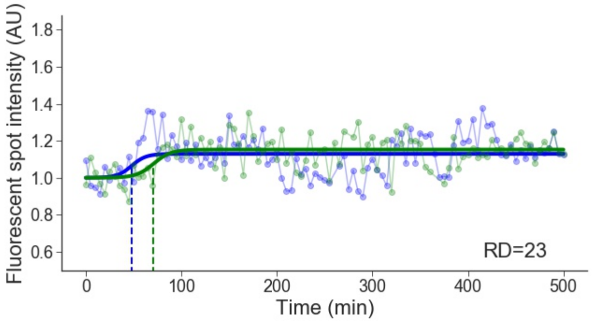


Cell line: Early 1, cell: S7 cell 2

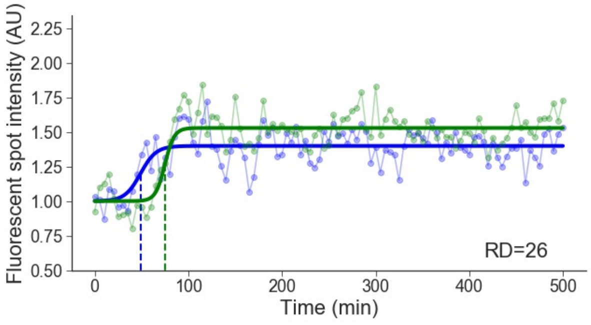


Cell line: Early 1, cell: S7 cell 3

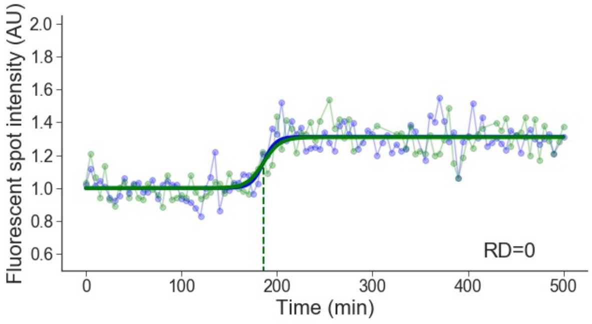


Cell line: Early 1, cell: S8 cell 1

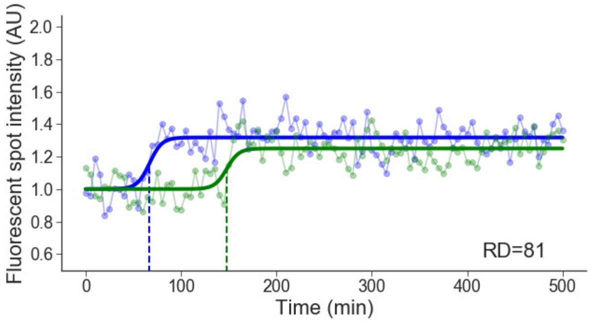


Cell line: Early 1, cell: S8 cell 2

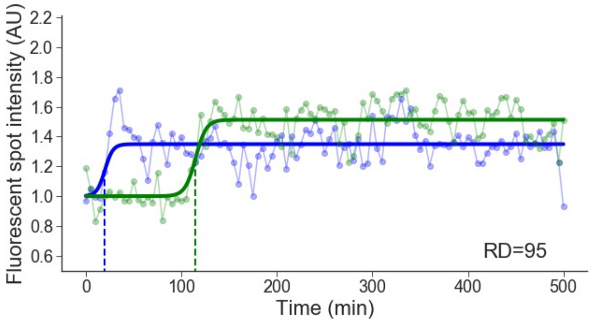


Cell line: Early 1, cell: S8 cell 3

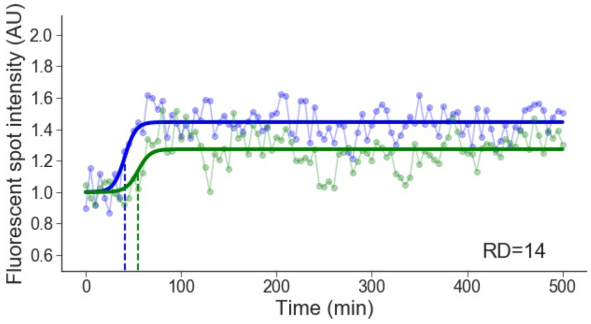


Cell line: Early 1, cell: S8 cell 4

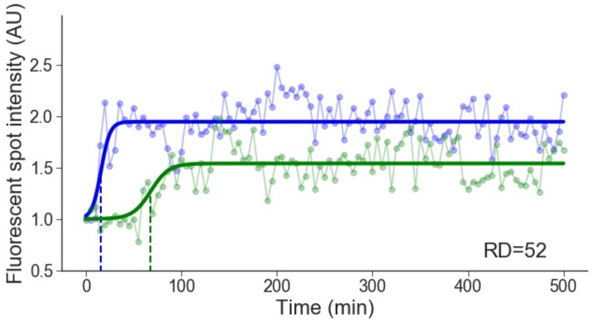


Cell line: Early 1, cell: S9 cell 1

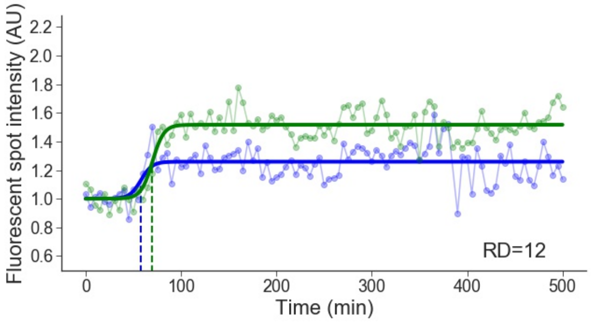

Cell line: Early 1, cell: S9 cell 3

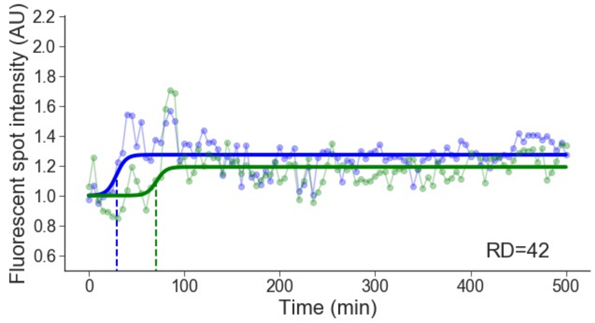


Cell line: Early 1, cell: S9 cell 4

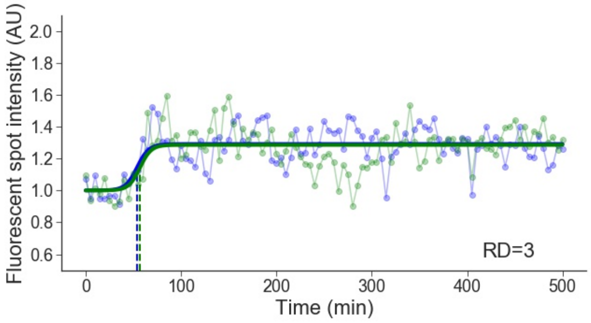


Cell line: Early 1, cell: S9 cell 5

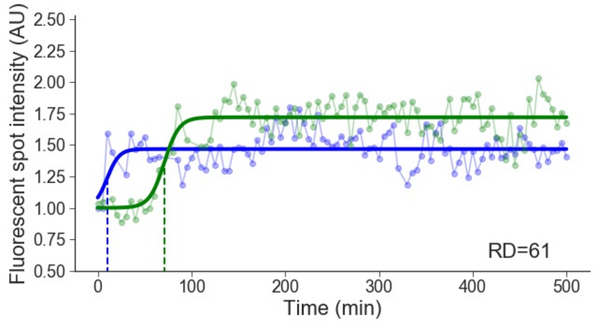


Cell line: Early 1, cell: S10 cell 1

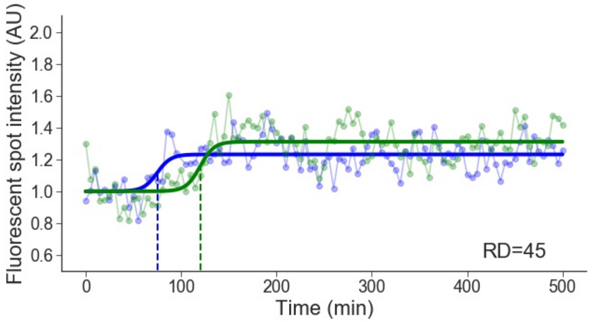


Cell line: Early 1, cell: S10 cell 2

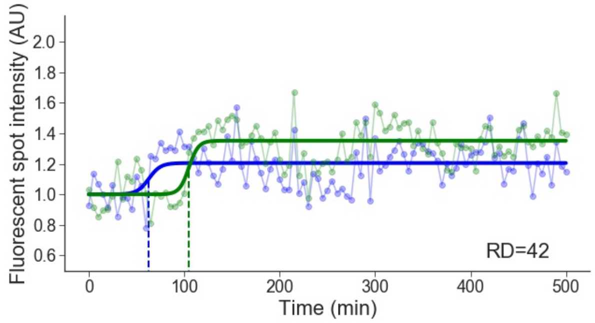


Cell line: Early 1, cell: S10 cell 5

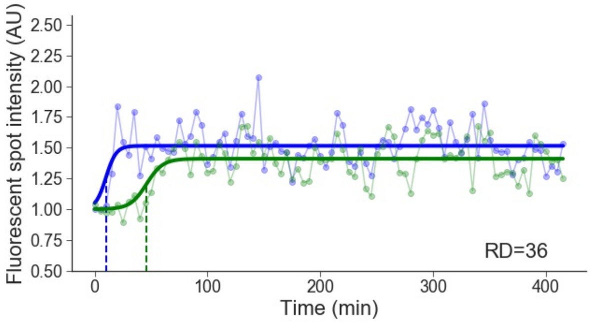


Cell line: Early 1, cell: S10 cell 7

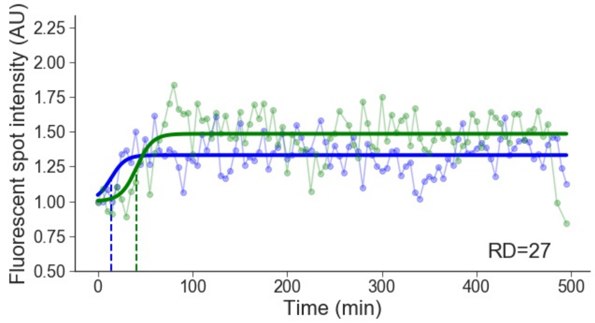


Cell line: Early 1, cell: S10 cell 8

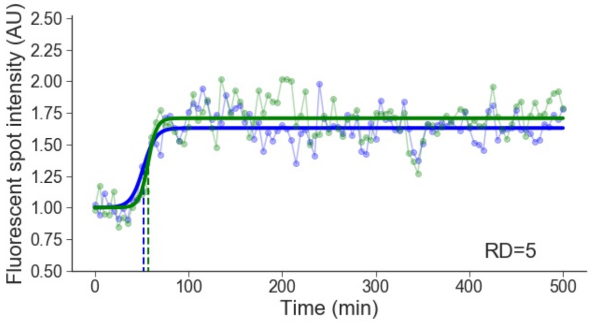

Cell line: Early 1, cell: S10 cell 9

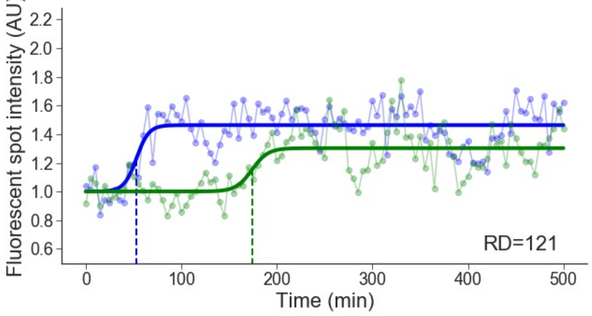


Cell line: Early 1, cell: S10 cell 11

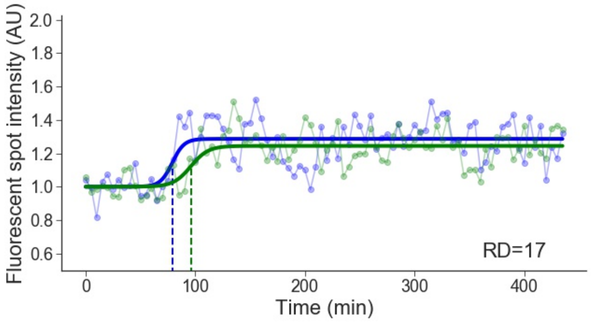


Cell line: Early 1, cell: S12 cell 1

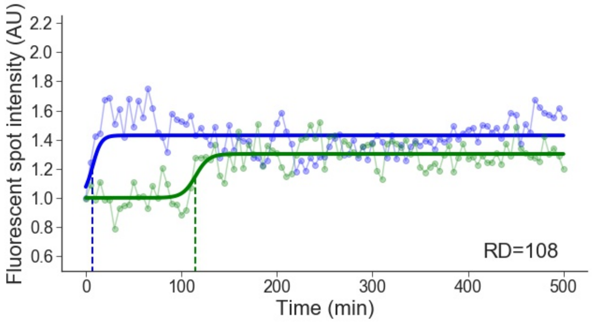


Cell line: Early 1, cell: S12 cell 2

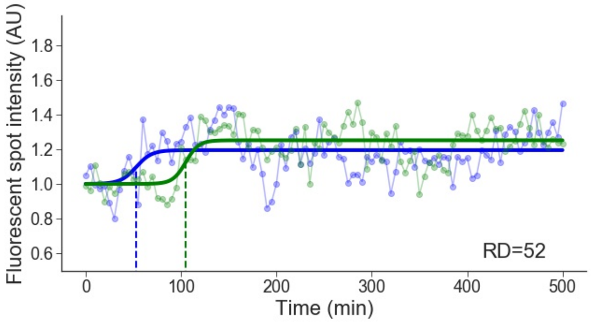


Cell line: Early 1, cell: S13 cell 1

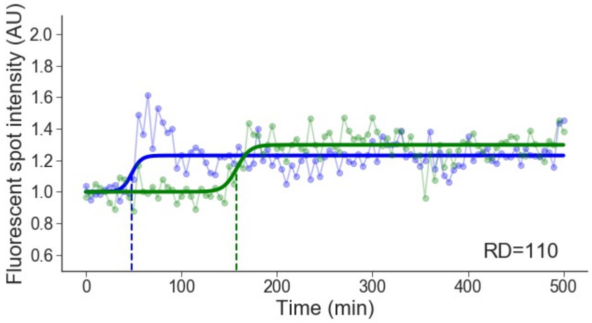


Cell line: Early 1, cell: S13 cell 2

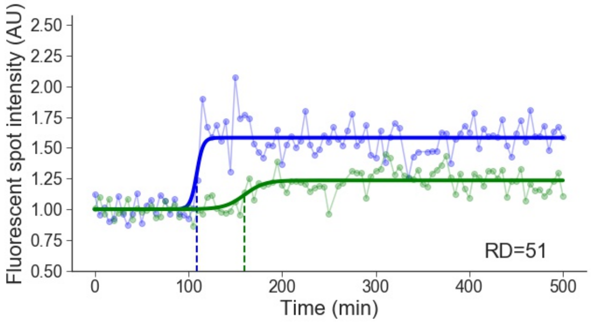


Cell line: Early 1, cell: S13 cell 3

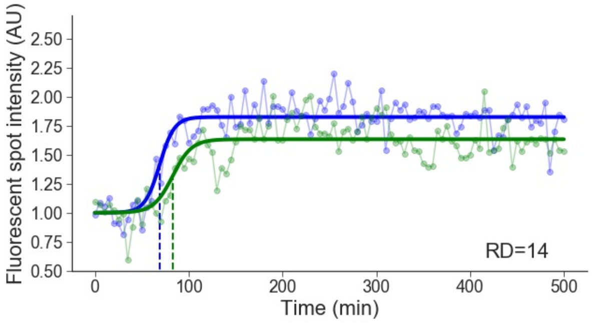


Cell line: Early 1, cell: S13 cell 4

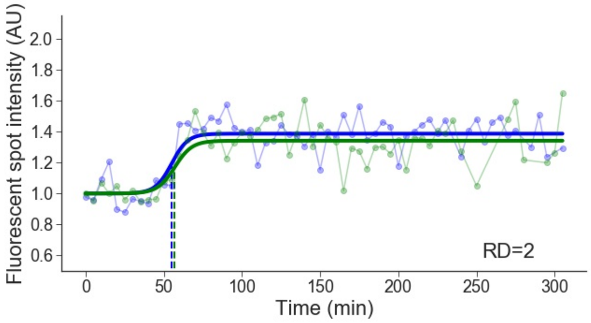

Cell line: Early 1, cell: S14 cell 2

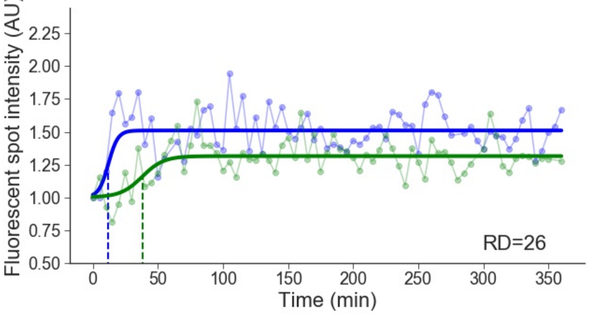


Cell line: Early 1, cell: S15 cell 2

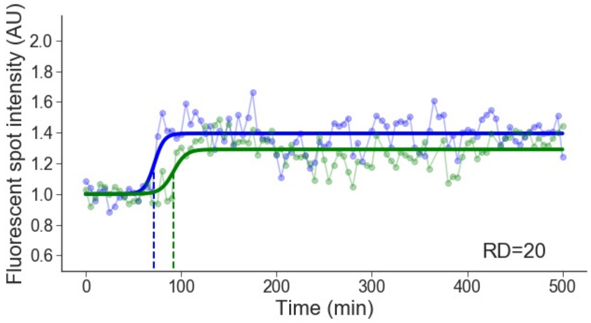


Cell line: Early 1, cell: S15 cell 3

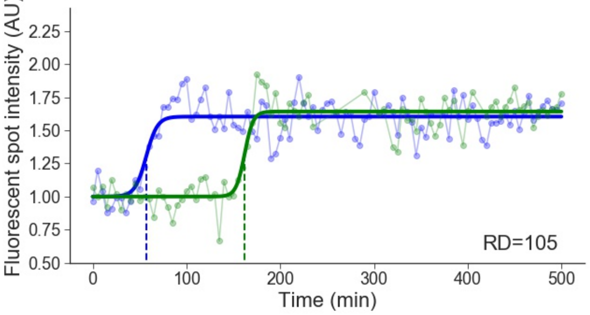


Cell line: Early 1, cell: S16 cell 1

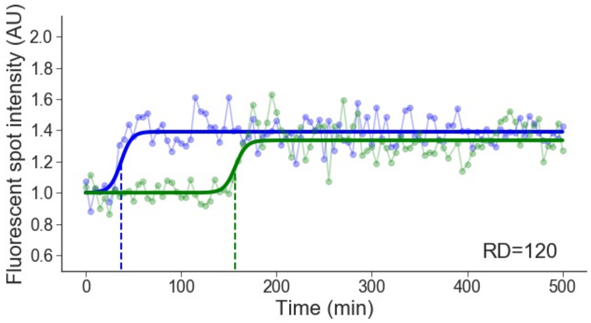


Cell line: Early 1, cell: S18 cell 2

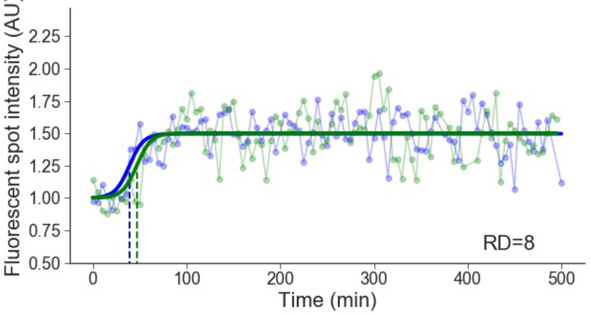


Cell line: Early 1, cell: S19 cell 2

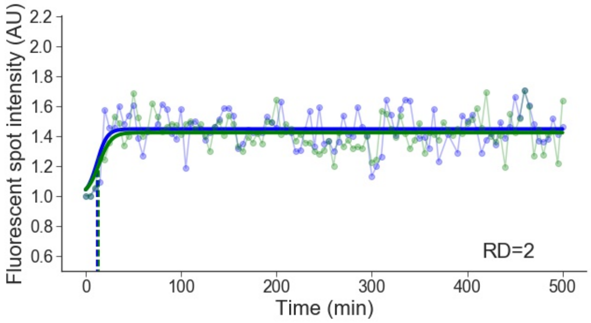


Cell line: Early 1, cell: S19 cell 3

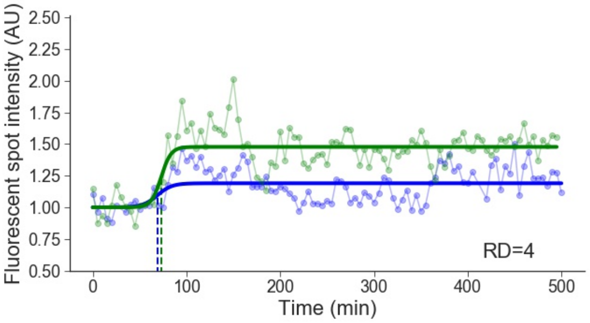


Cell line: Early 1, cell: S20 cell 2

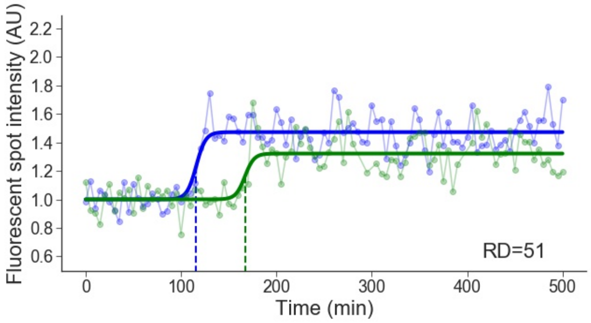

Cell line: Early 1, cell: S20 cell 3

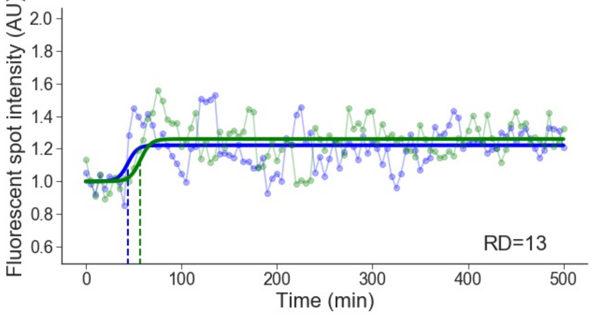


Cell line: Early 1, cell: S20 cell 4

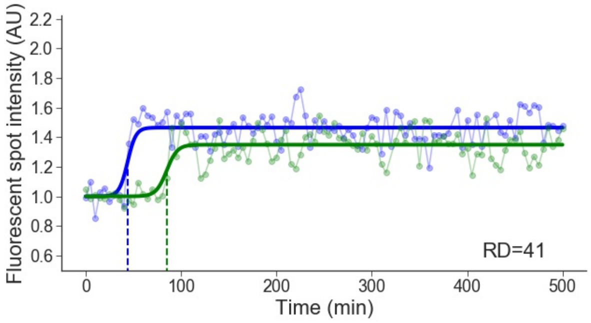


Cell line: Early 1, cell: S20 cell 5

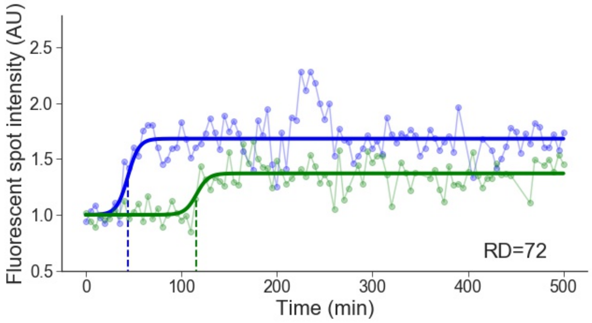

### 2 Early 2

Cell line: Early 2, cell: S3 cell 1

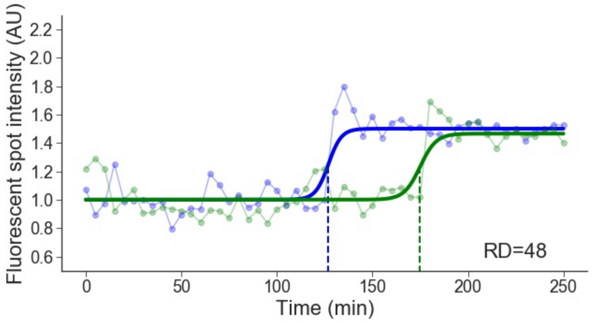


Cell line: Early 2, cell: S3 cell 3

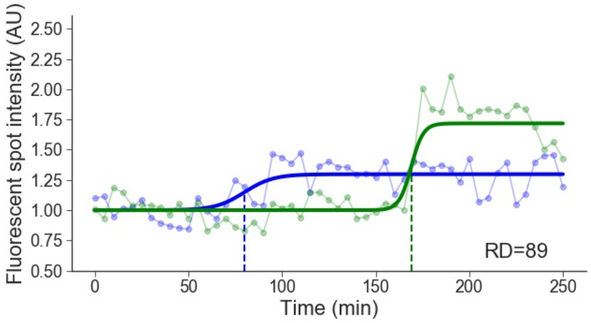


Cell line: Early 2, cell: S3 cell 4

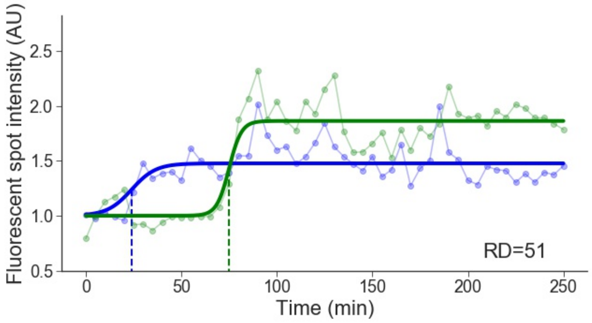


Cell line: Early 2, cell: S4 cell 3

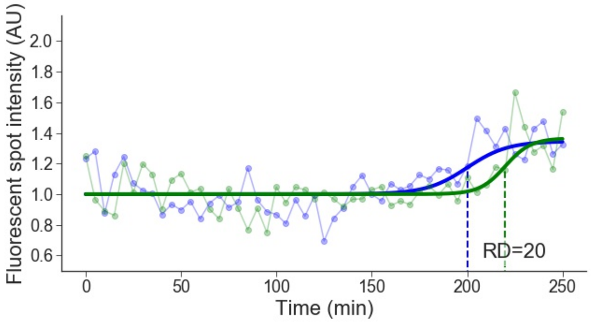


Cell line: Early 2, cell: S4 cell 6

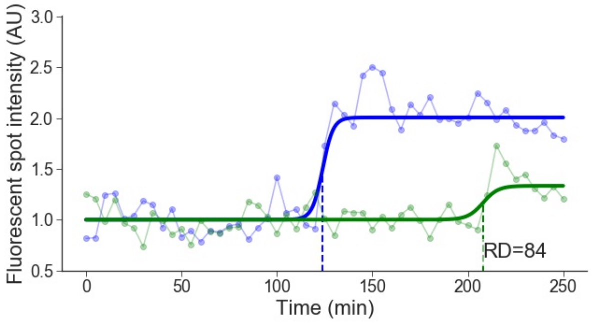


Cell line: Early 2, cell: S5 cell 1

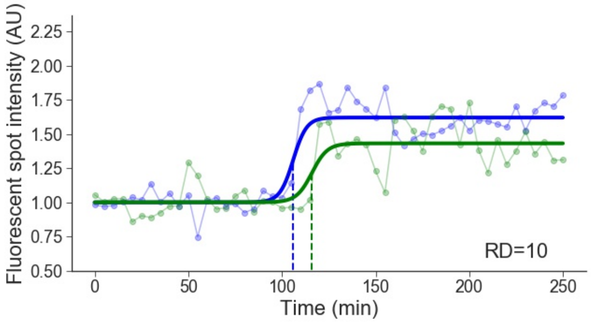


Cell line: Early 2, cell: S6 cell 1

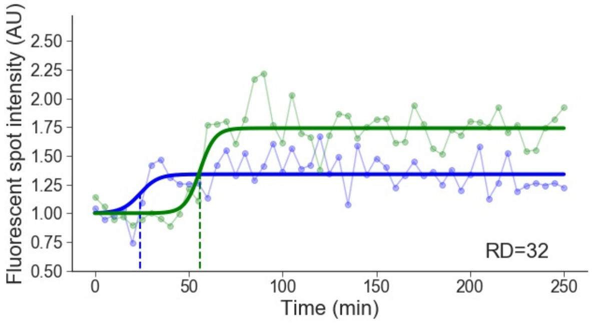


Cell line: Early 2, cell: S7 cell 1

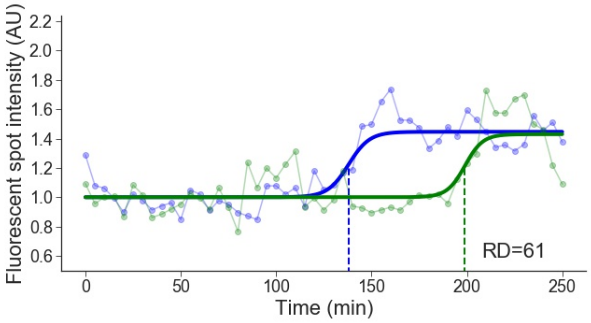

Cell line: Early 2, cell: S7 cell 2

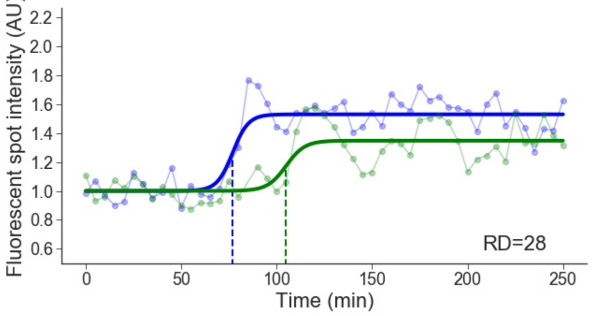


Cell line: Early 2, cell: S8 cell 3

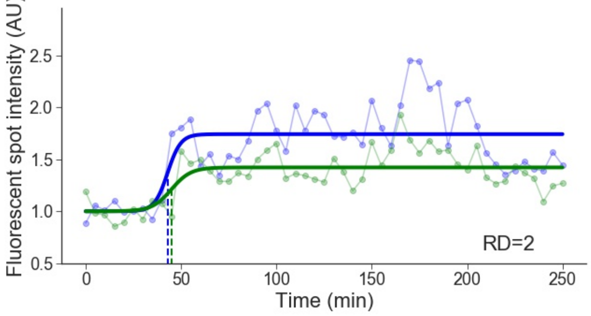


Cell line: Early 2, cell: S8 cell 4

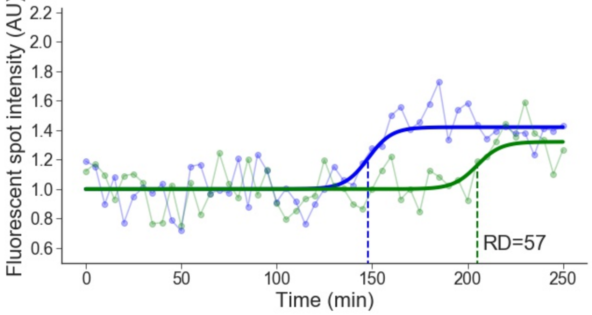


Cell line: Early 2, cell: S8 cell 5

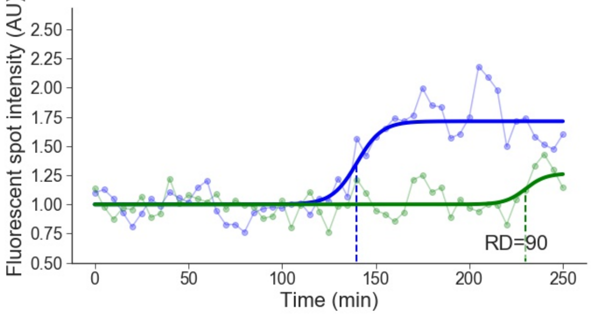


Cell line: Early 2, cell: S9 cell 1

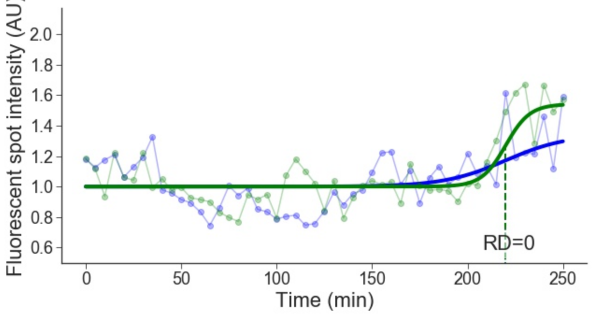


Cell line: Early 2, cell: S9 cell 2

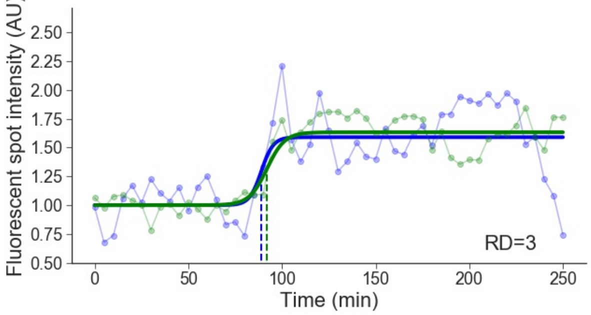


Cell line: Early 2, cell: S9 cell 3

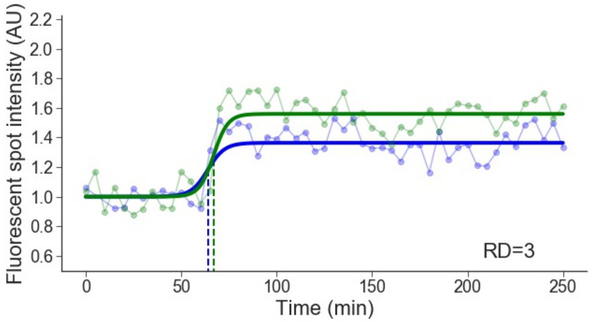


Cell line: Early 2, cell: S10 cell 5

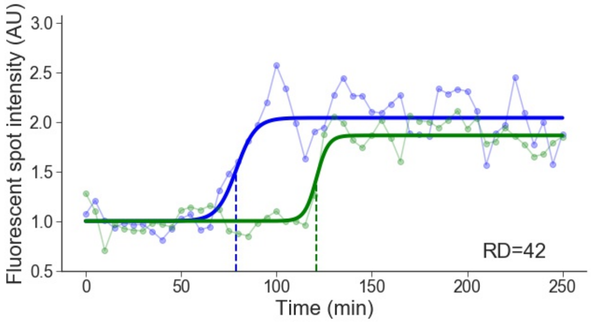

Cell line: Early 2, cell: S10 cell 6

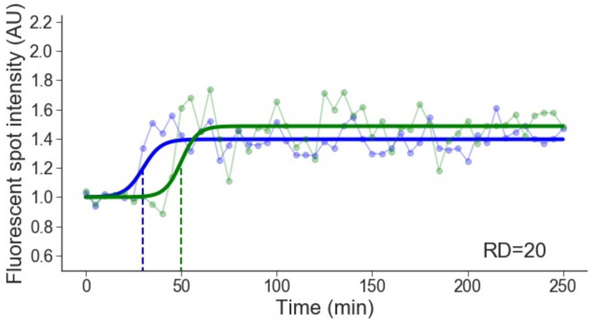


Cell line: Early 2, cell: S11 cell 1

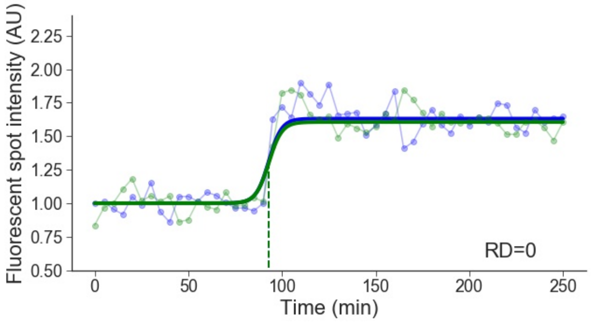


Cell line: Early 2, cell: S11 cell 4

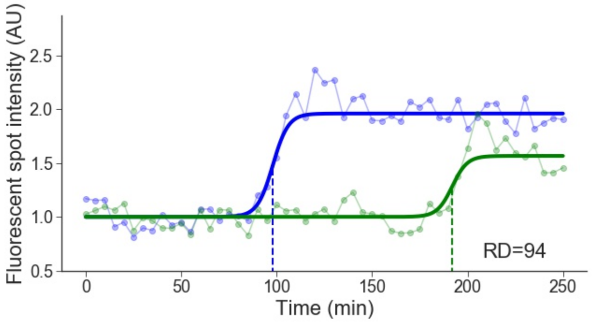


Cell line: Early 2, cell: S11 cell 5

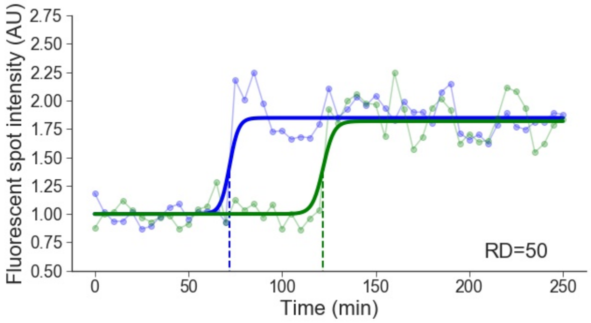


Cell line: Early 2, cell: S11 cell 6

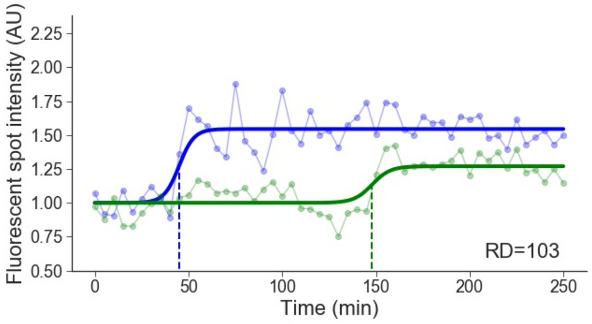


Cell line: Early 2, cell: S12 cell 1

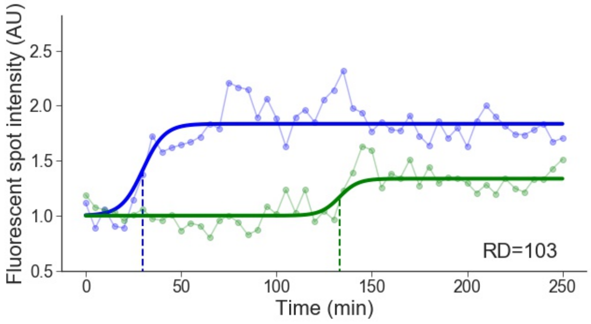


Cell line: Early 2, cell: S12 cell 2

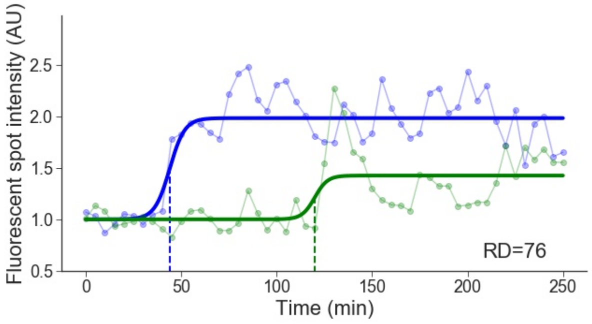


Cell line: Early 2, cell: S12 cell 4

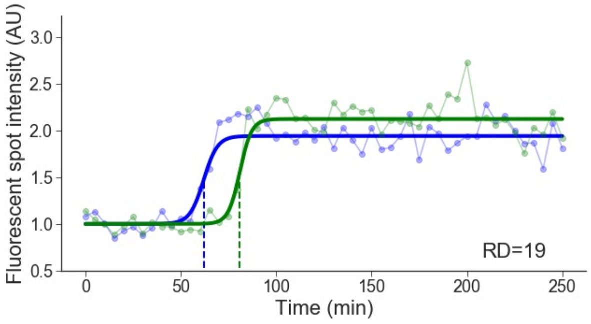

Cell line: Early 2, cell: S13 cell 2

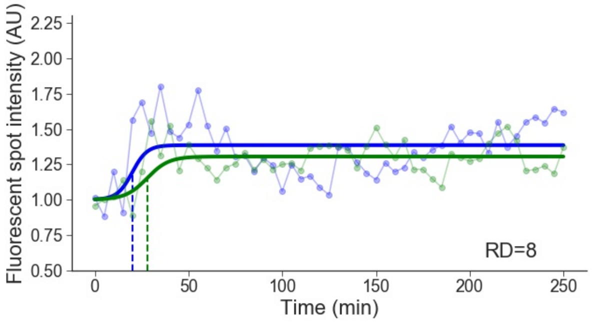


Cell line: Early 2, cell: S13 cell 5

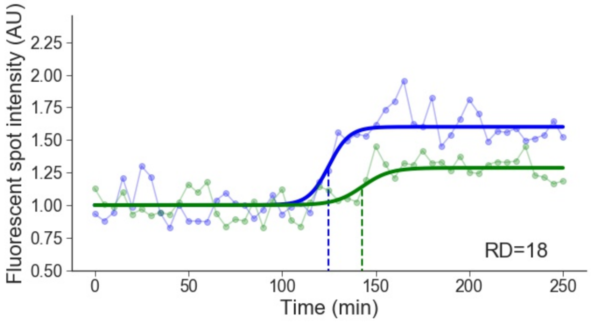


Cell line: Early 2, cell: S14 cell 1

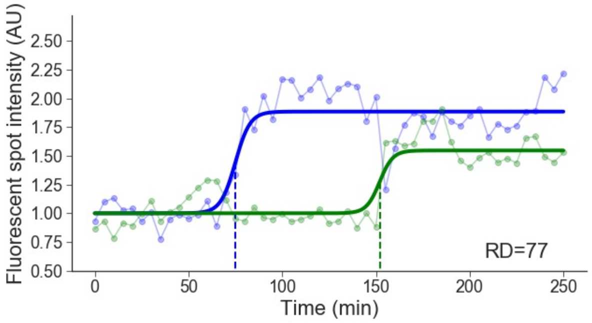


Cell line: Early 2, cell: S14 cell 3

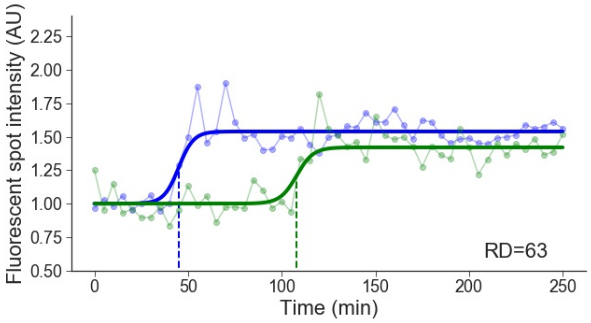


Cell line: Early 2, cell: S14 cell 4

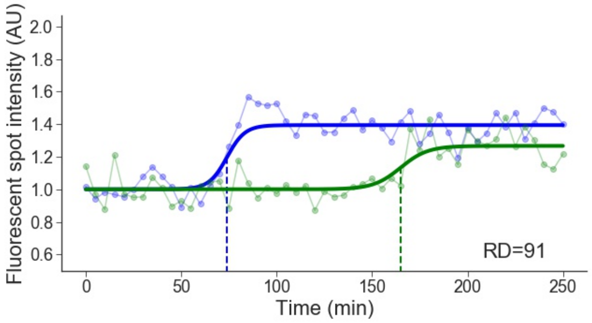


Cell line: Early 2, cell: S14 cell 6

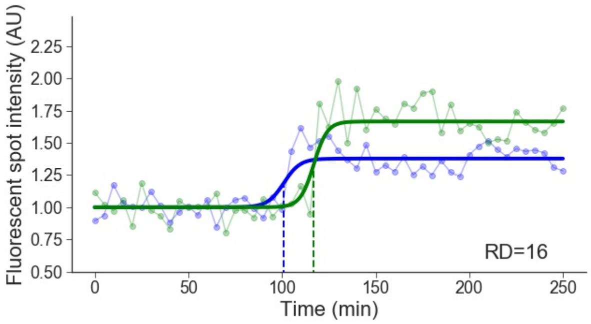


Cell line: Early 2, cell: S14 cell 7

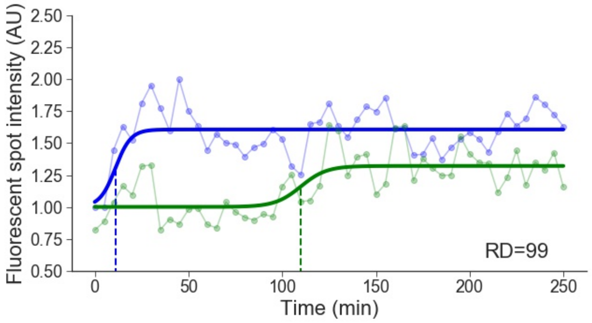


Cell line: Early 2, cell: S15 cell 2

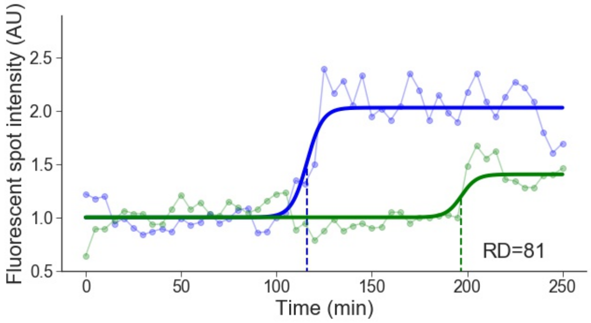

Cell line: Early 2, cell: S15 cell 3

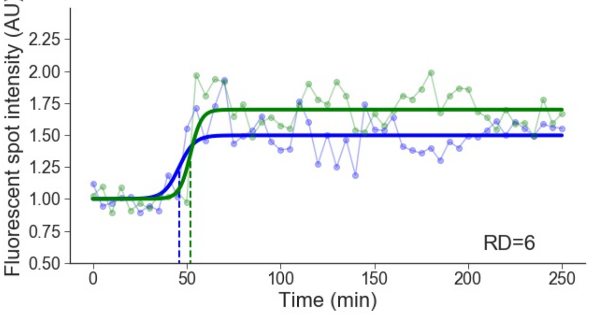


Cell line: Early 2, cell: S15 cell 4

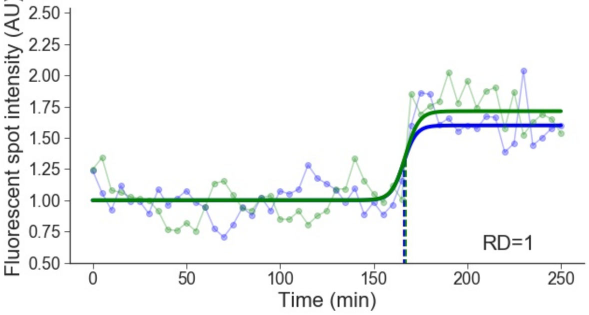


Cell line: Early 2, cell: S16 cell 1

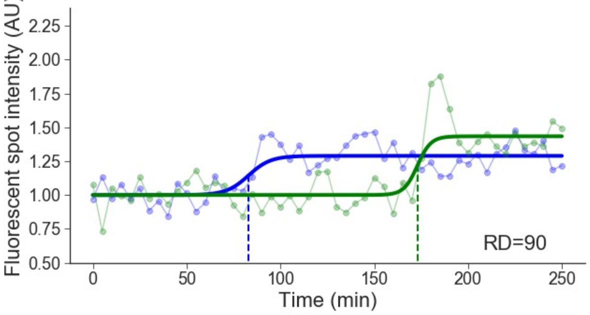


Cell line: Early 2, cell: S16 cell 2

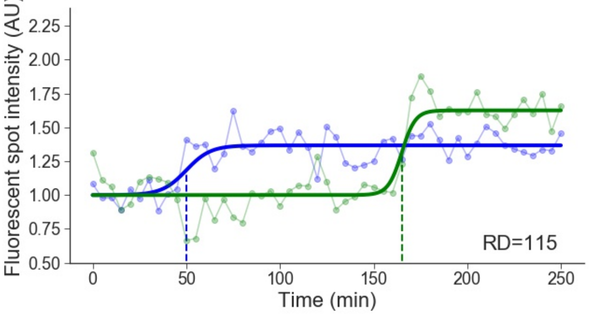


Cell line: Early 2, cell: S16 cell 4

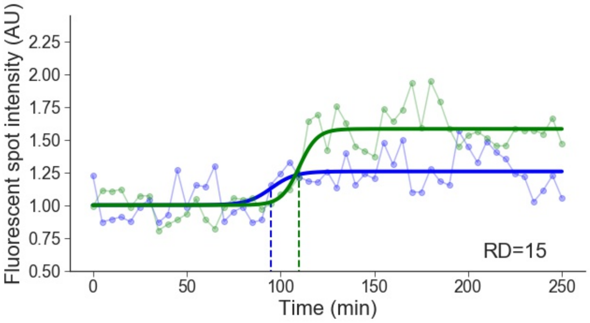


Cell line: Early 2, cell: S16 cell 5

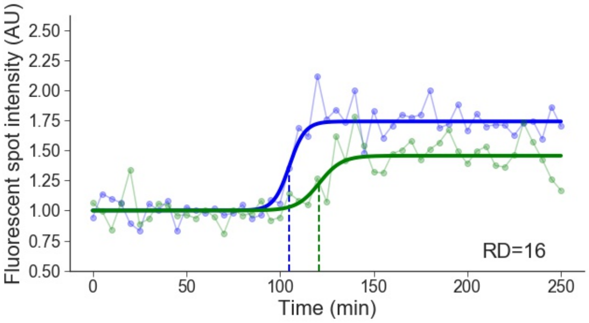


Cell line: Early 2, cell: S16 cell 7

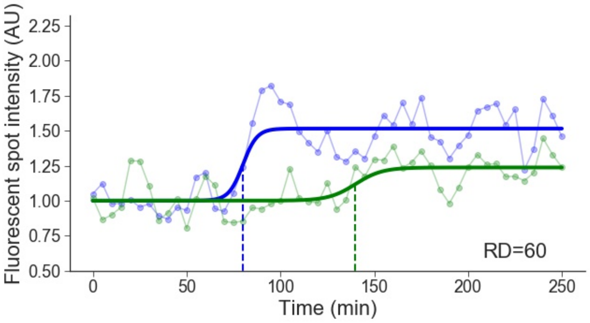


Cell line: Early 2, cell: S17 cell 1

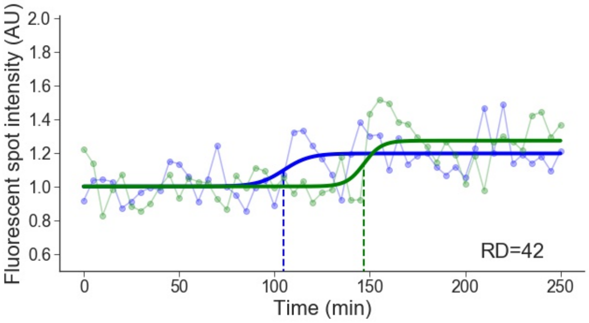

Cell line: Early 2, cell: S17 cell 3

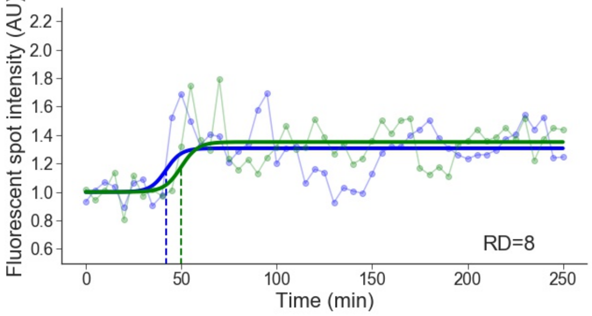


Cell line: Early 2, cell: S17 cell 4

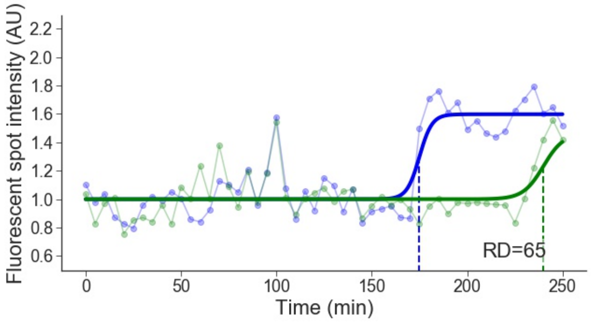


Cell line: Early 2, cell: S17 cell 5

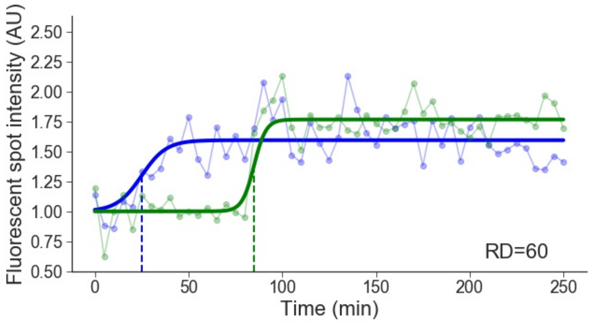


Cell line: Early 2, cell: S17 cell 6

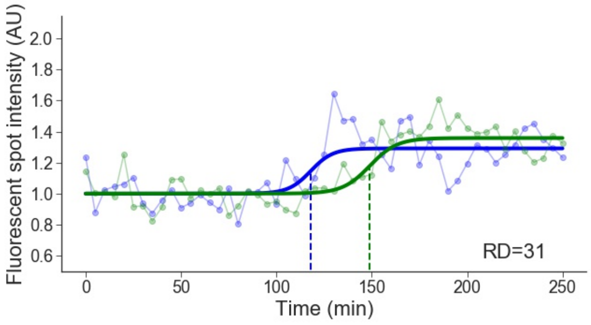


Cell line: Early 2, cell: S18 cell 3

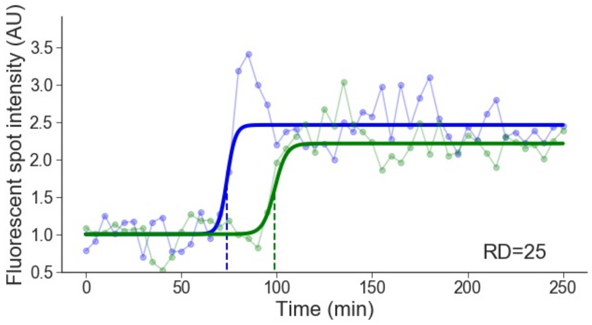


Cell line: Early 2, cell: S19 cell 2

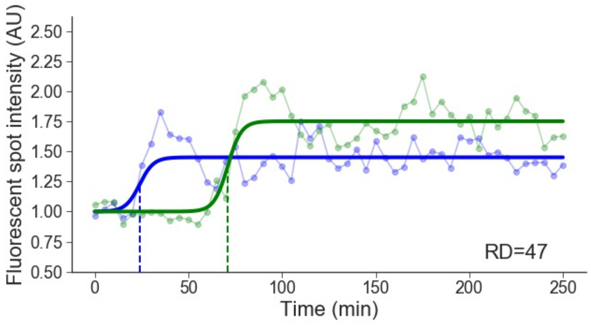


Cell line: Early 2, cell: S19 cell 3

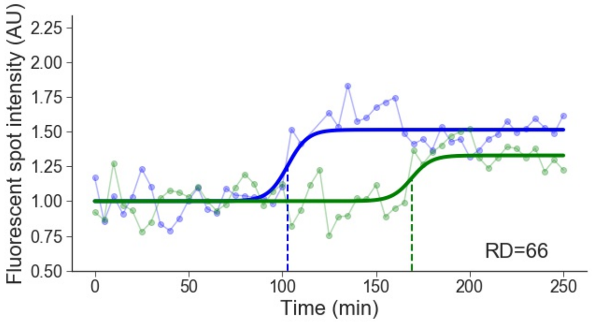


Cell line: Early 2, cell: S19 cell 4

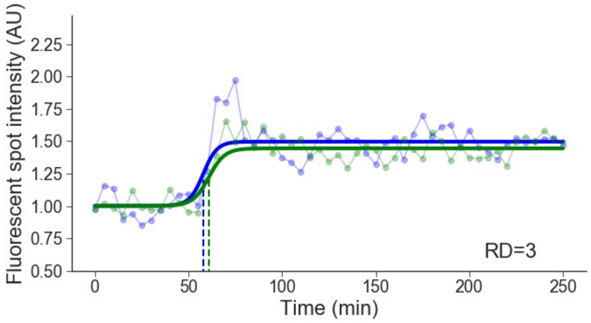

Cell line: Early 2, cell: S19 cell 5

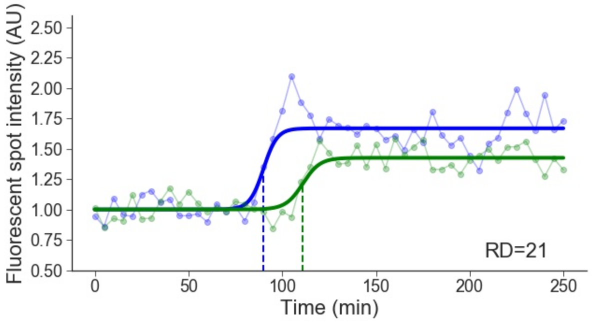


Cell line: Early 2, cell: S19 cell 7

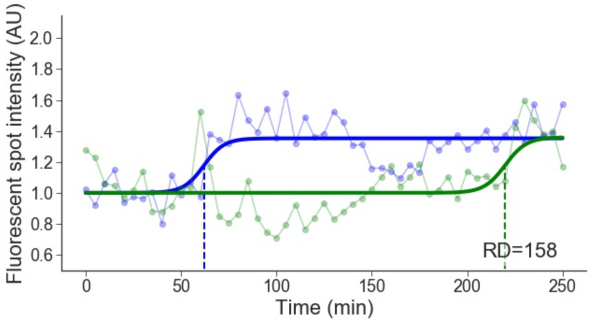


Cell line: Early 2, cell: S19 cell 9

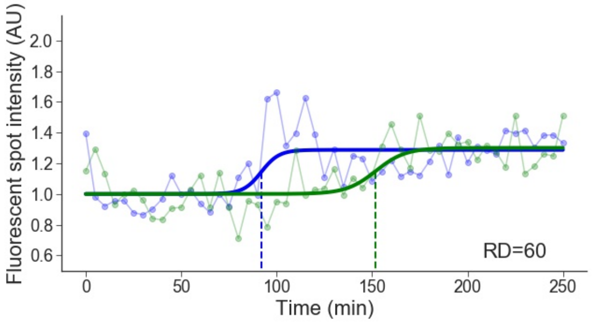

### 3 Early 2 + Mid-late 2

Cell line: Early 2 + Mid-late 2, cell: S5 cell 1

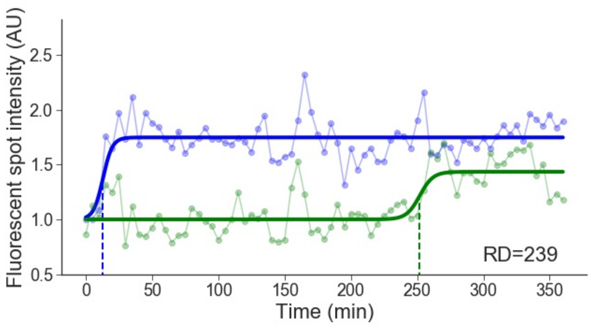


Cell line: Early 2 + Mid-late 2, cell: S9 cell 5

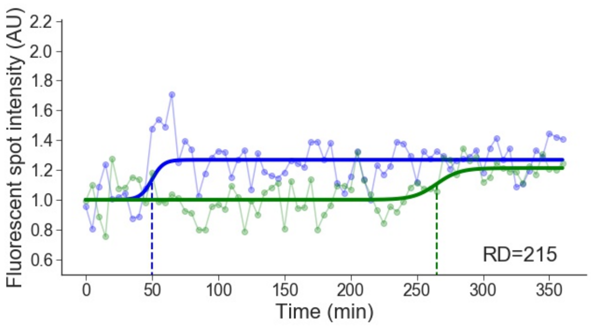


Cell line: Early 2 + Mid-late 2, cell: S10 cell 7

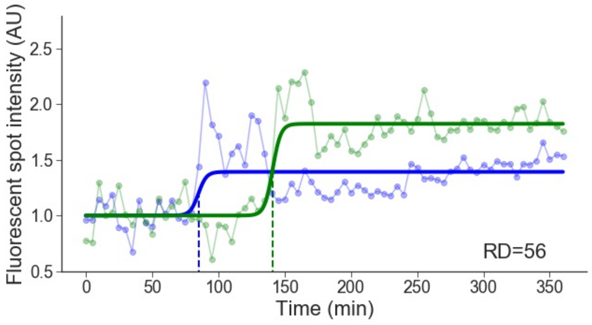


Cell line: Early 2 + Mid-late 2, cell: S12 cell 2

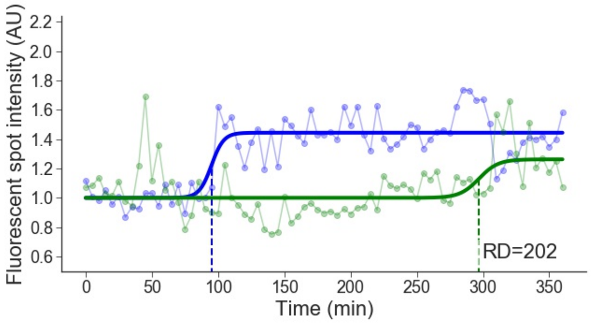


Cell line: Early 2 + Mid-late 2, cell: S13 cell 3

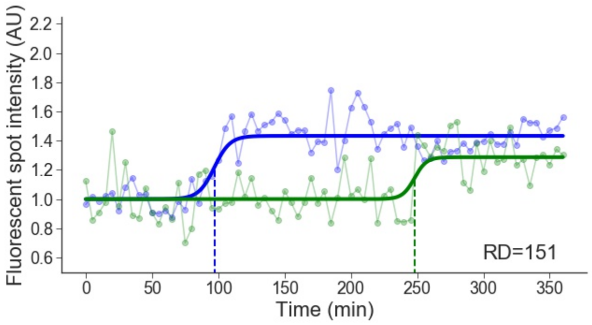


Cell line: Early 2 + Mid-late 2, cell: S13 cell 4

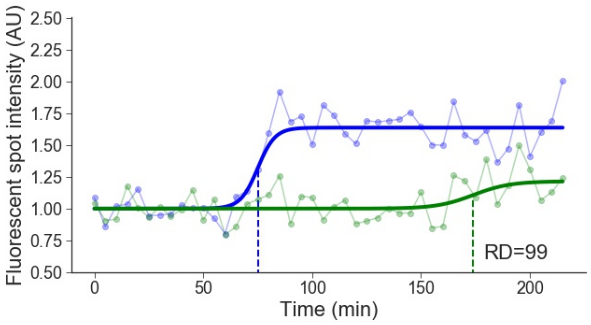


Cell line: Early 2 + Mid-late 2, cell: S14 cell 4

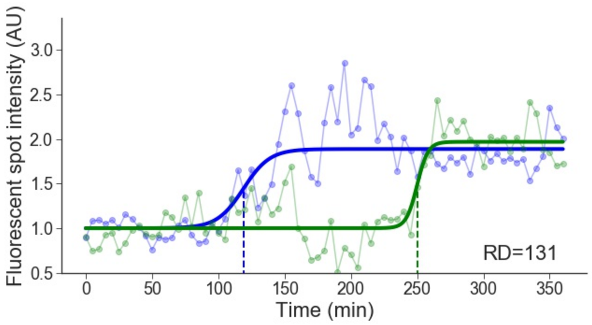


Cell line: Early 2 + Mid-late 2, cell: S14 cell 5

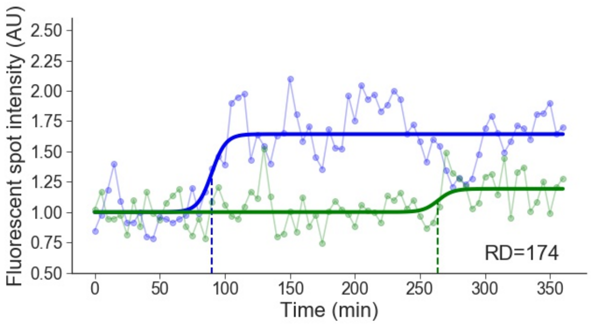

Cell line: Early 2 + Mid-late 2, cell: S15 cell 4

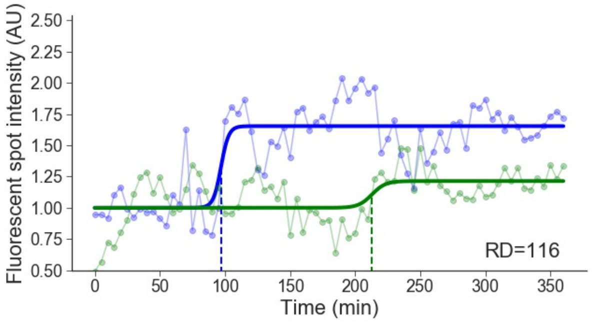


Cell line: Early 2 + Mid-late 2, cell: S17 cell 2

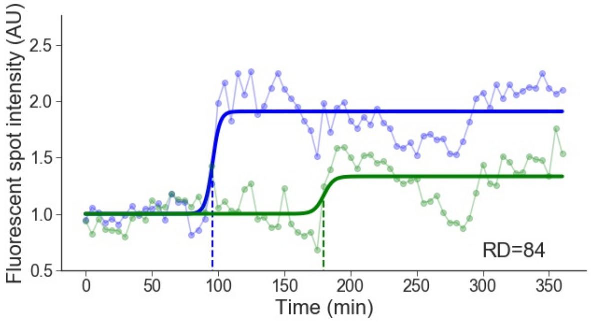


Cell line: Early 2 + Mid-late 2, cell: S17 cell 4

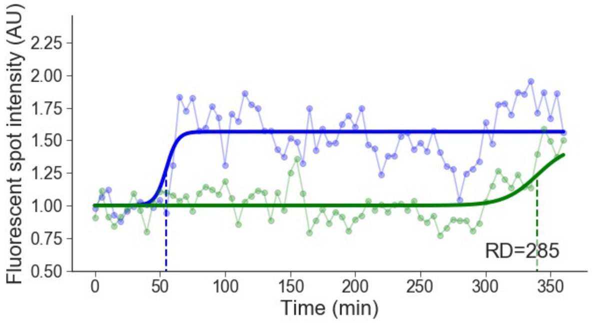


Cell line: Early 2 + Mid-late 2, cell: S18 cell 1

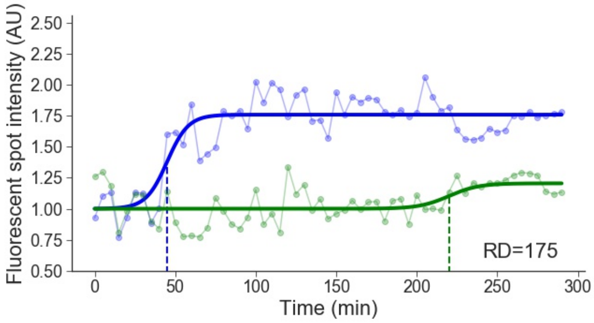


Cell line: Early 2 + Mid-late 2, cell: S18 cell 2

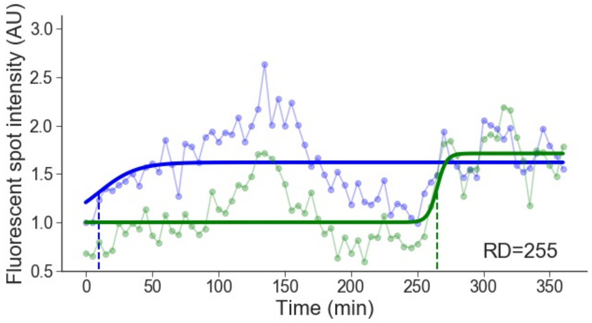


Cell line: Early 2 + Mid-late 2, cell: S18 cell 3

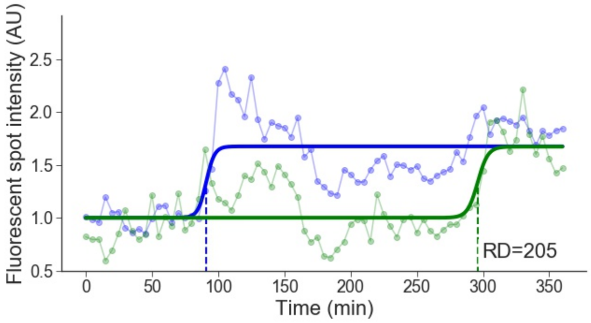


Cell line: Early 2 + Mid-late 2, cell: S19 cell 4

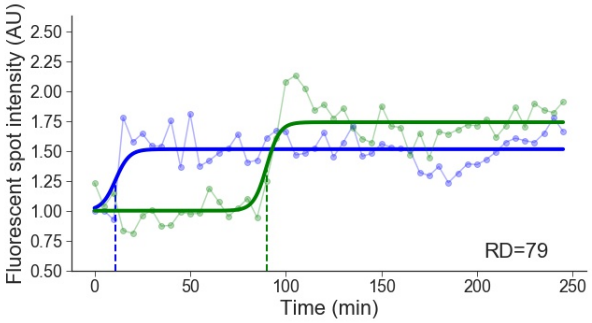


Cell line: Early 2 + Mid-late 2, cell: S20 cell 1

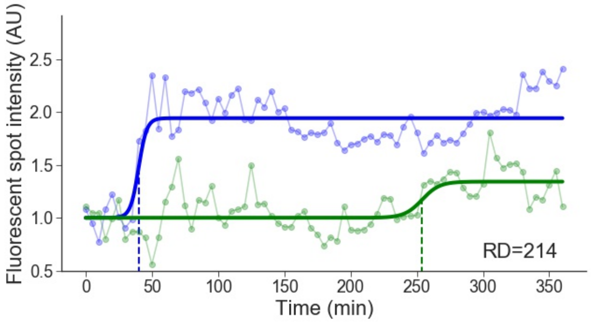

Cell line: Early 2 + Mid-late 2, cell: S23 cell 2

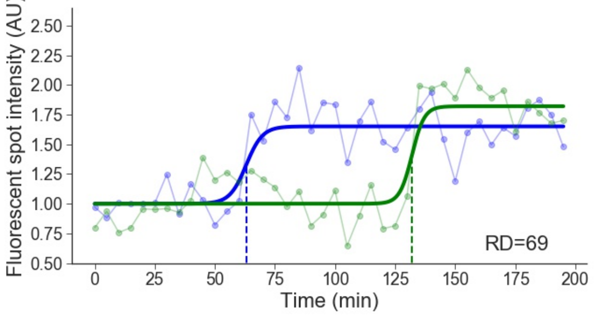


Cell line: Early 2 + Mid-late 2, cell: S24 cell 2

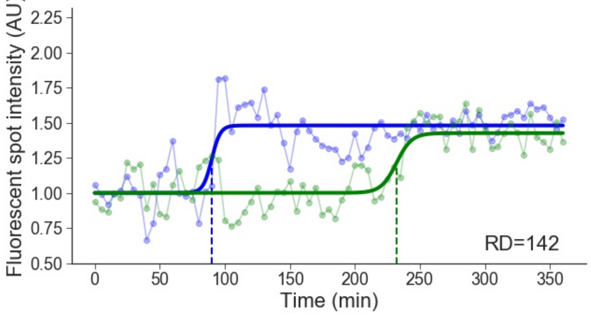


Cell line: Early 2 + Mid-late 2, cell: S24 cell 3

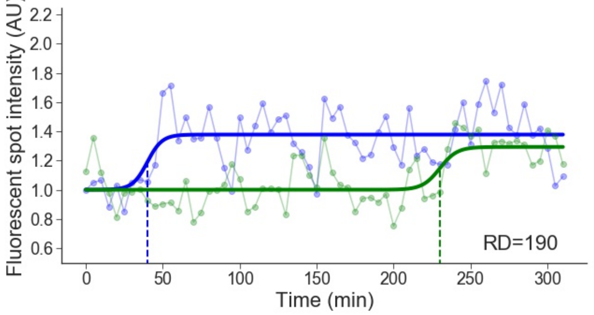


Cell line: Early 2 + Mid-late 2, cell: S28 cell 1

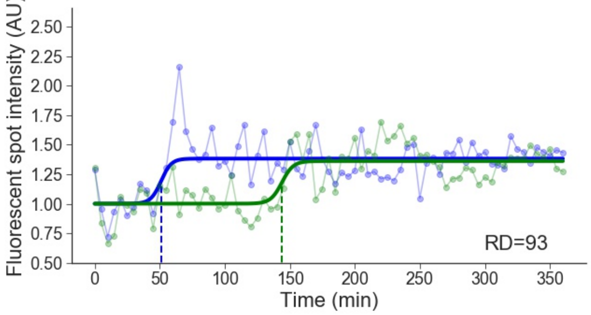


Cell line: Early 2 + Mid-late 2, cell: S28 cell 5

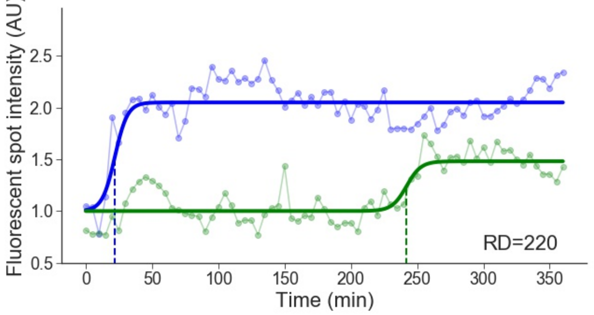


Cell line: Early 2 + Mid-late 2, cell: S30 cell 4

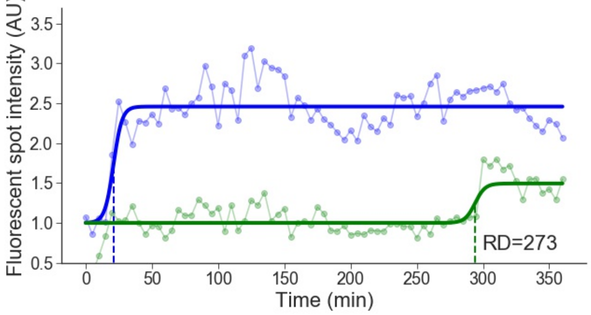


Cell line: Early 2 + Mid-late 2, cell: S34 cell 3

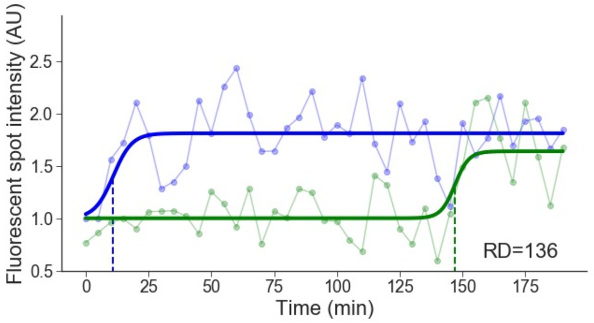


Cell line: Early 2 + Mid-late 2, cell: S38 cell 2

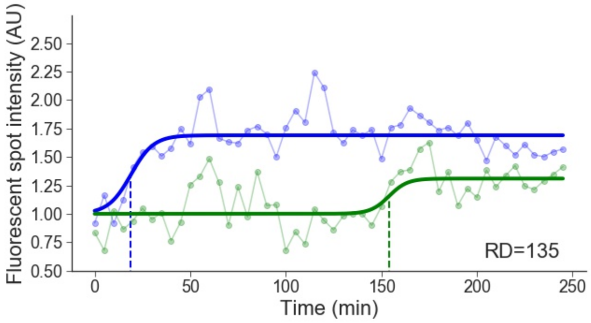

### 4 Mid-late 1

Cell line: Mid-late 1, cell: S2 cell 2

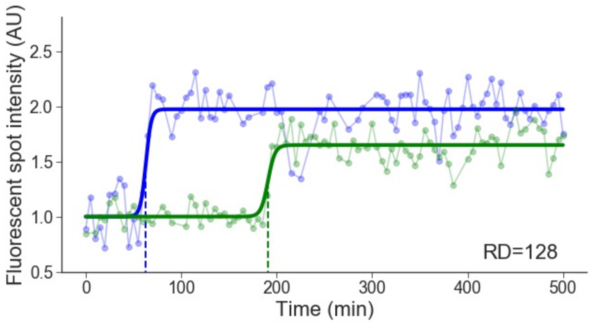


Cell line: Mid-late 1, cell: S2 cell 3

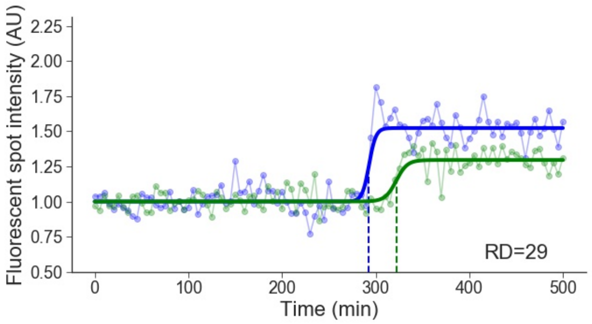


Cell line: Mid-late 1, cell: S4 cell 1

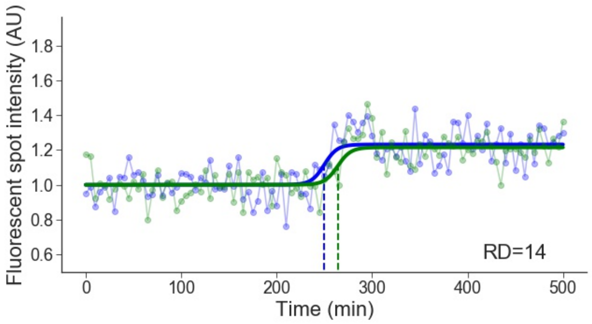


Cell line: Mid-late 1, cell: S5 cell 3

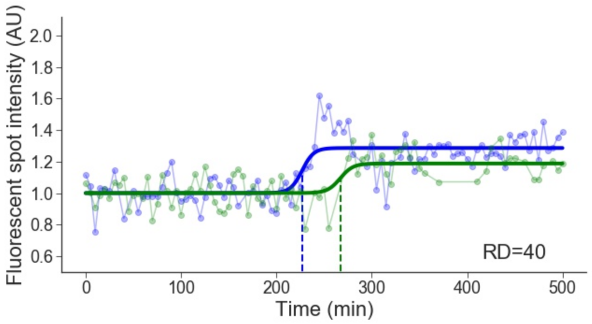


Cell line: Mid-late 1, cell: S7 cell 5

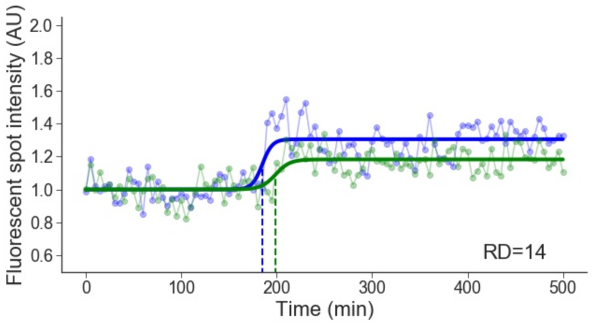


Cell line: Mid-late 1, cell: S7 cell 6

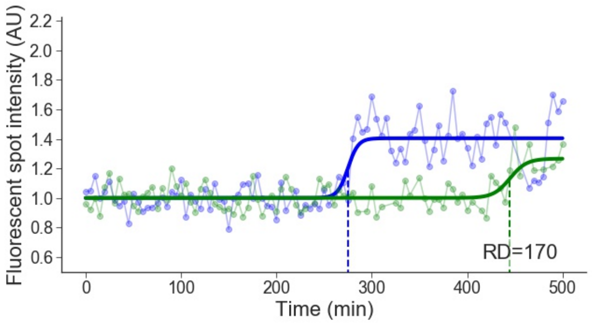


Cell line: Mid-late 1, cell: S8 cell 1

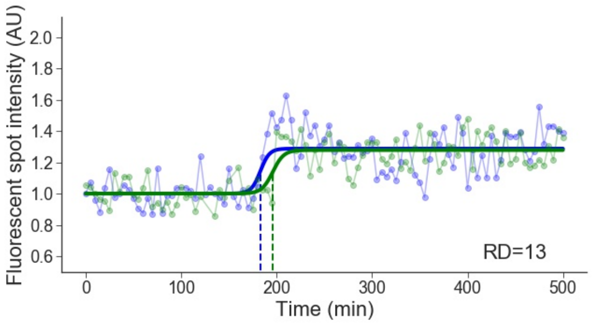


Cell line: Mid-late 1, cell: S8 cell 2

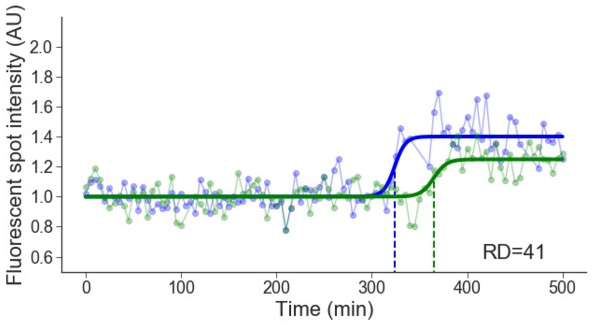

Cell line: Mid-late 1, cell: S8 cell 3

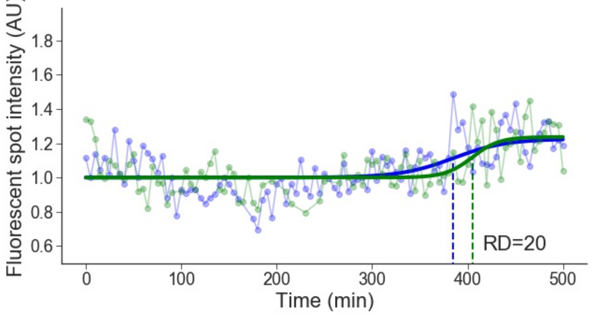


Cell line: Mid-late 1, cell: S8 cell 6

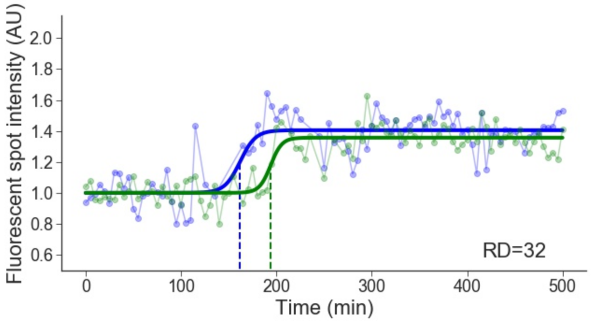


Cell line: Mid-late 1, cell: S11 cell 2

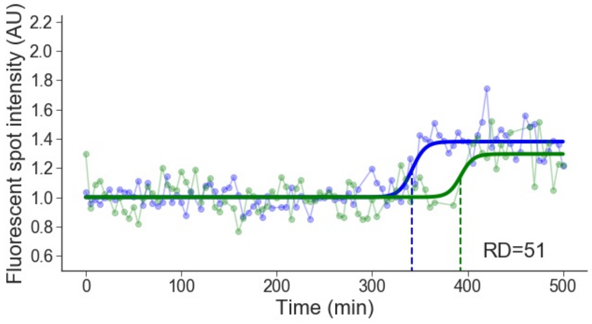


Cell line: Mid-late 1, cell: S12 cell 1

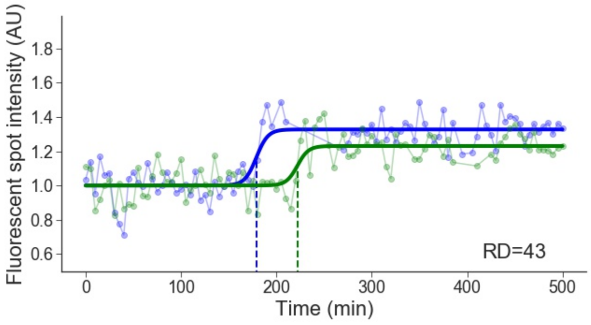


Cell line: Mid-late 1, cell: S12 cell 2

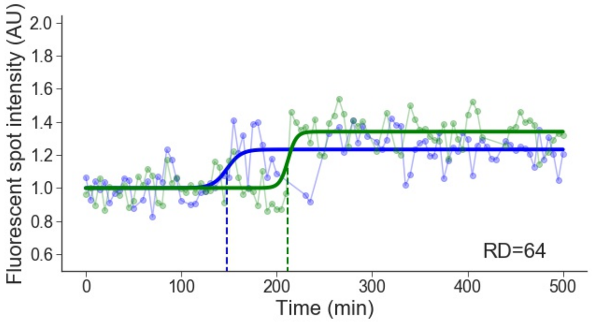


Cell line: Mid-late 1, cell: S12 cell 4

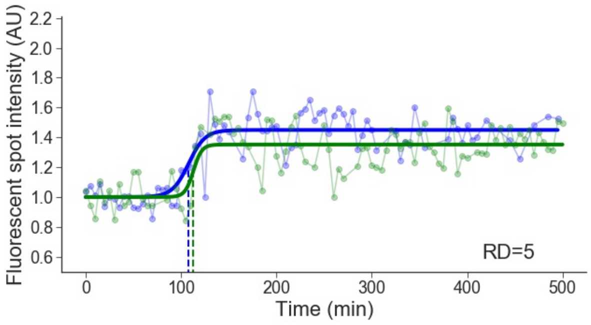


Cell line: Mid-late 1, cell: S15 cell 2

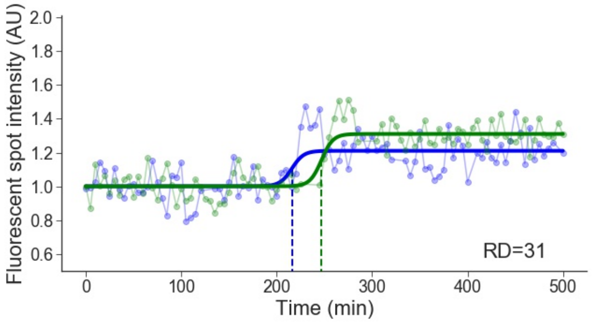


Cell line: Mid-late 1, cell: S27 cell 1

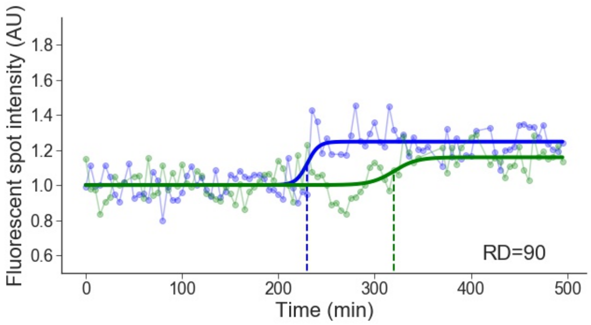

Cell line: Mid-late 1, cell: S27 cell 2

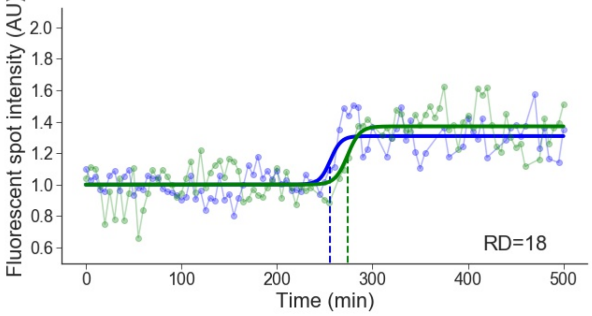


Cell line: Mid-late 1, cell: S27 cell 3

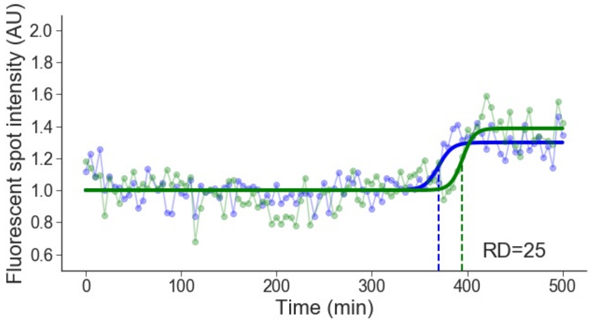


Cell line: Mid-late 1, cell: S27 cell 4

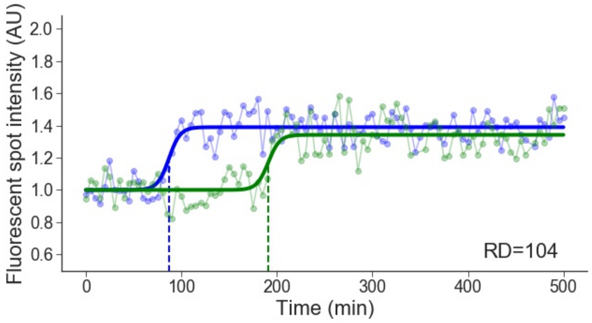


Cell line: Mid-late 1, cell: S27 cell 5

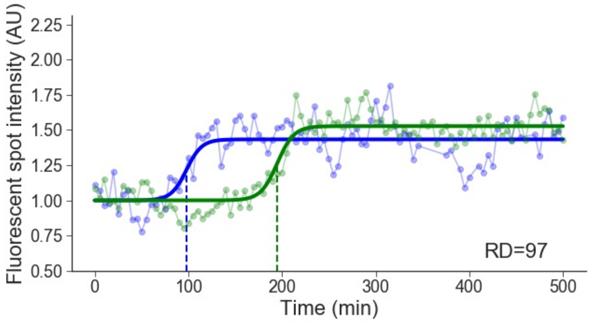


Cell line: Mid-late 1, cell: S27 cell 6

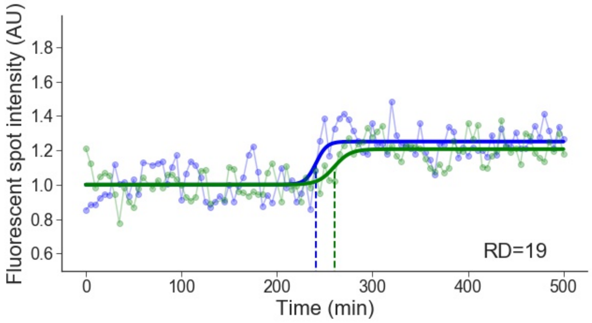


Cell line: Mid-late 1, cell: S28 cell 2

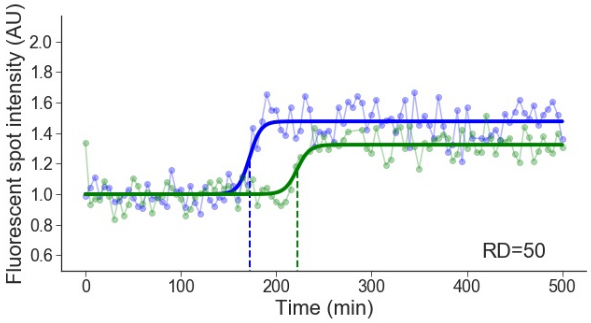


Cell line: Mid-late 1, cell: S28 cell 3

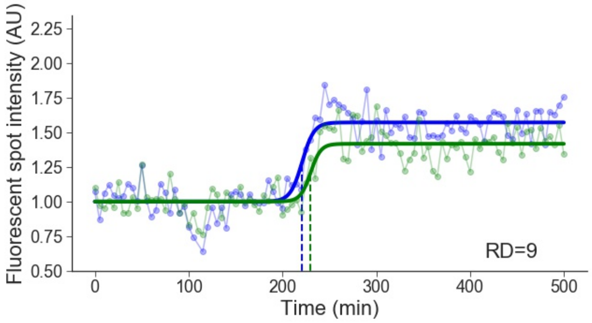


Cell line: Mid-late 1, cell: S30 cell 1

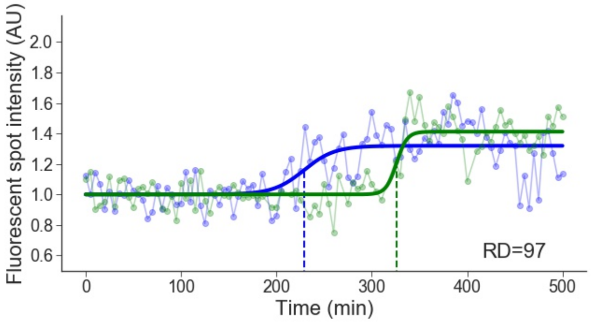

Cell line: Mid-late 1, cell: S30 cell 2

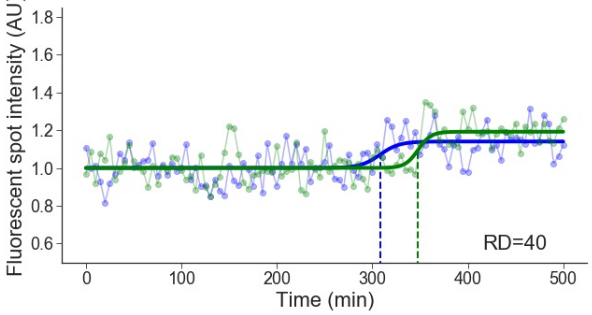


Cell line: Mid-late 1, cell: S31 cell 1

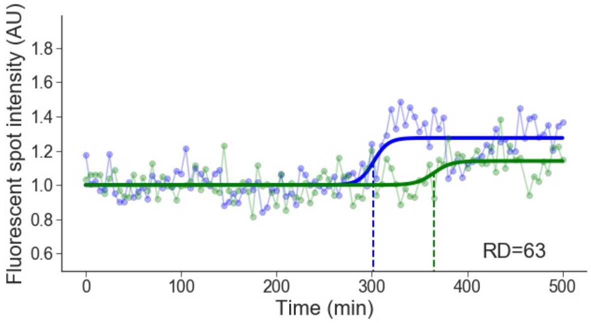


Cell line: Mid-late 1, cell: S31 cell 2

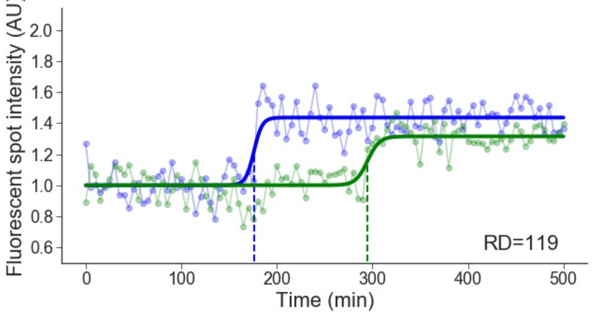


Cell line: Mid-late 1, cell: S34 cell 1

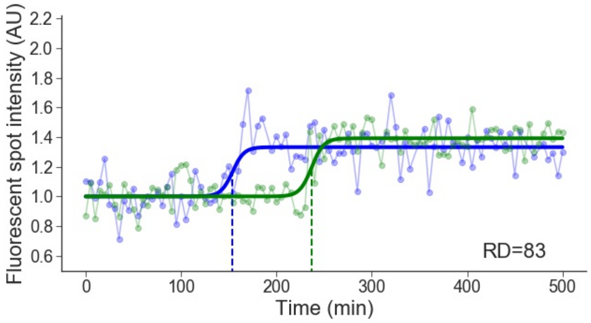


Cell line: Mid-late 1, cell: S34 cell 2

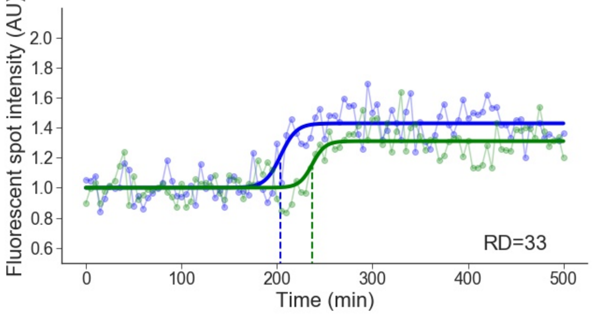


Cell line: Mid-late 1, cell: S34 cell 4

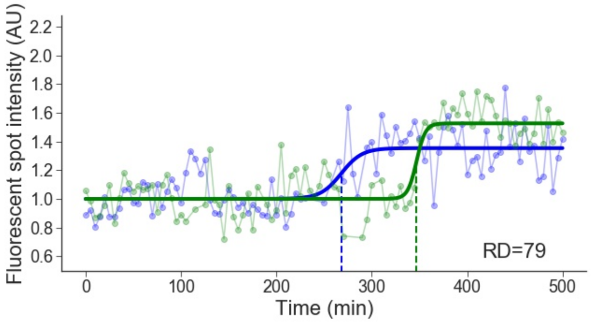


Cell line: Mid-late 1, cell: S34 cell 5

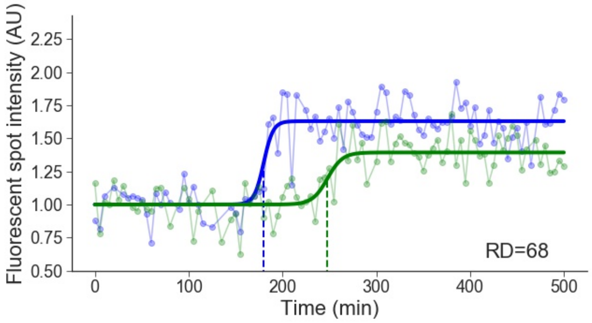


Cell line: Mid-late 1, cell: S34 cell 6

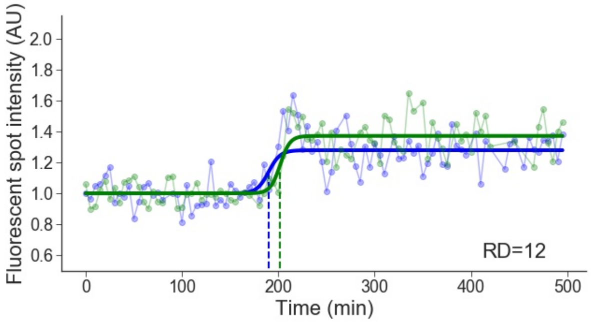

Cell line: Mid-late 1, cell: S34 cell 7

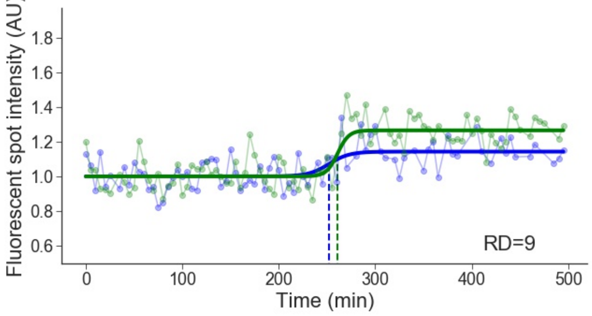


Cell line: Mid-late 1, cell: S35 cell 1

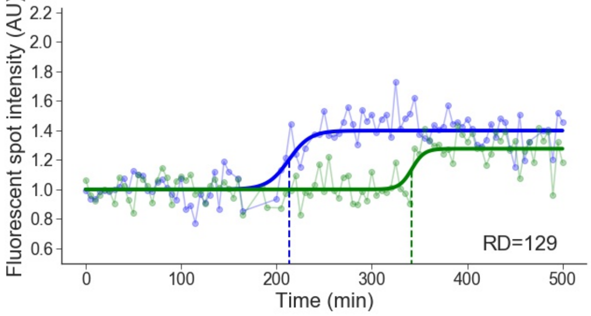


Cell line: Mid-late 1, cell: S35 cell 2

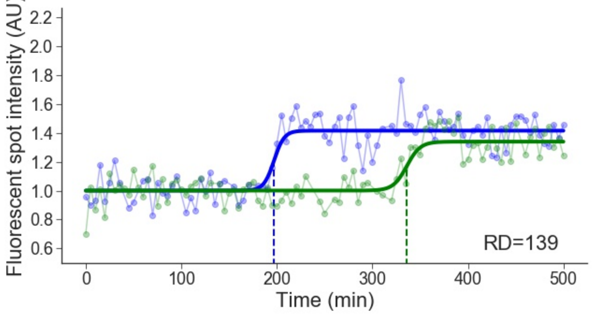


Cell line: Mid-late 1, cell: S35 cell 4

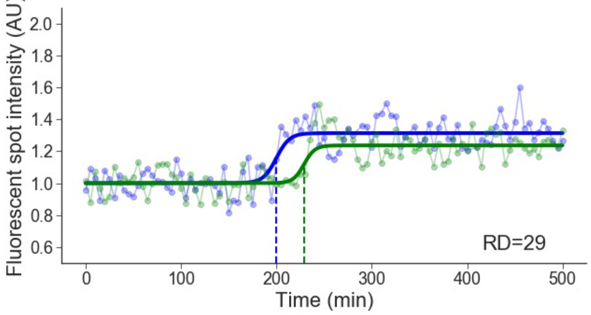


Cell line: Mid-late 1, cell: S35 cell 5

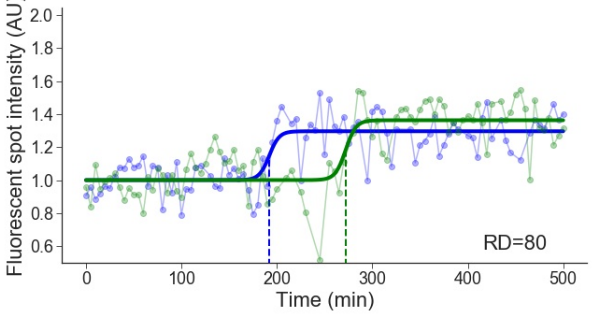


Cell line: Mid-late 1, cell: S38 cell 1

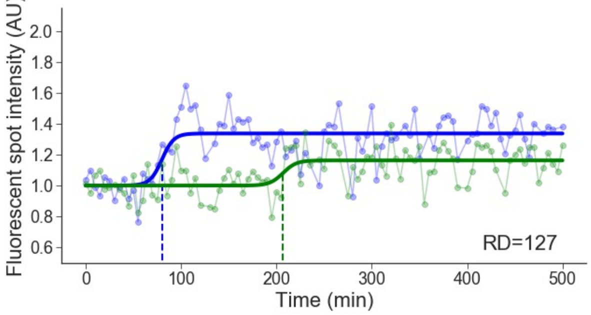


Cell line: Mid-late 1, cell: S38 cell 2

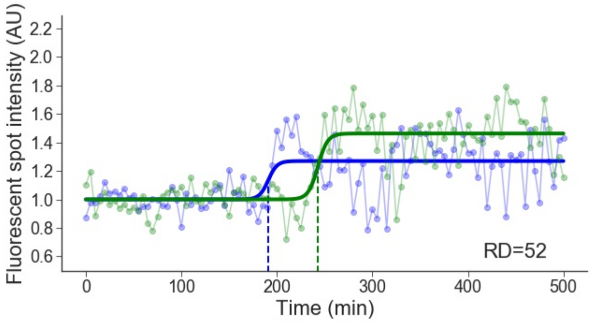


Cell line: Mid-late 1, cell: S39 cell 1

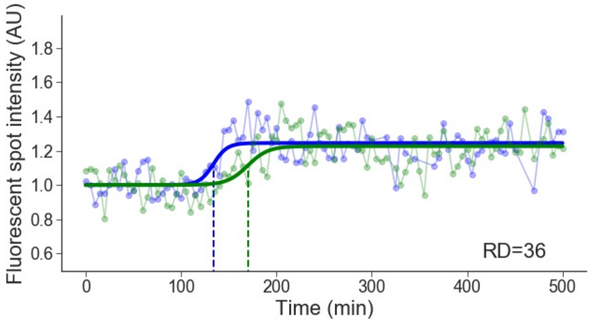

Cell line: Mid-late 1, cell: S40 cell 1

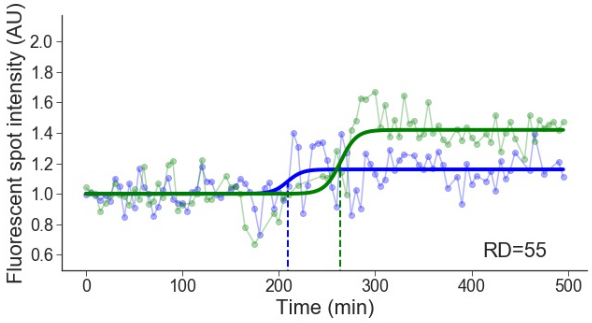


Cell line: Mid-late 1, cell: S40 cell 2

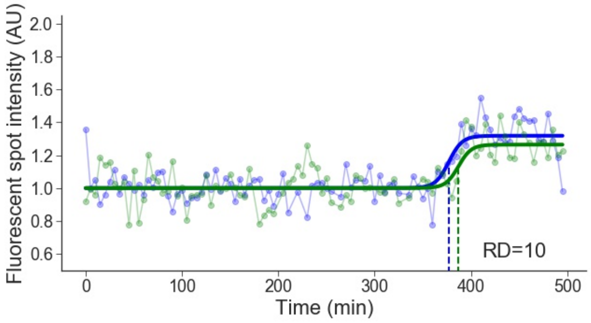


Cell line: Mid-late 1, cell: S40 cell 3

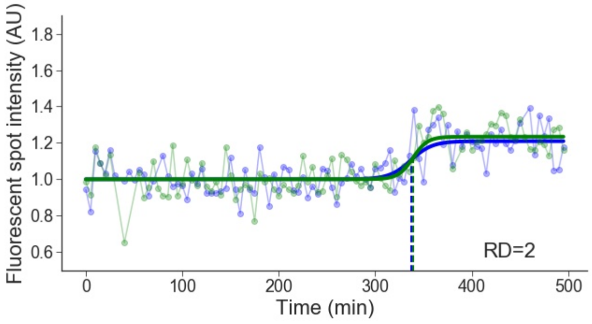


Cell line: Mid-late 1, cell: S40 cell 4

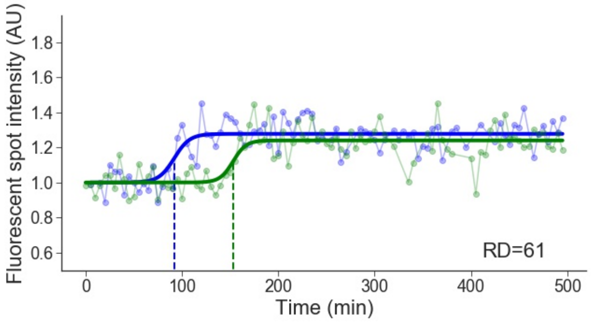


Cell line: Mid-late 1, cell: S40 cell 5

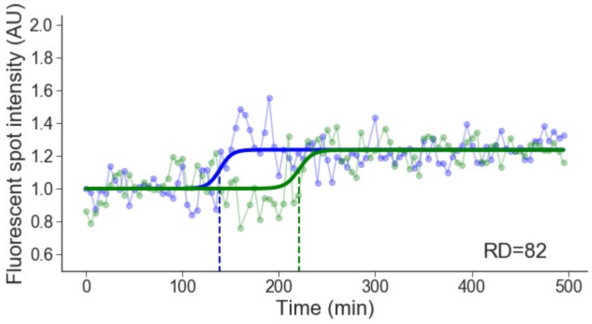


Cell line: Mid-late 1, cell: S40 cell 6

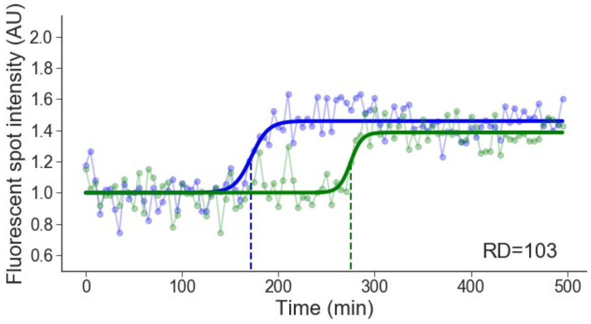


Cell line: Mid-late 1, cell: S42 cell 4

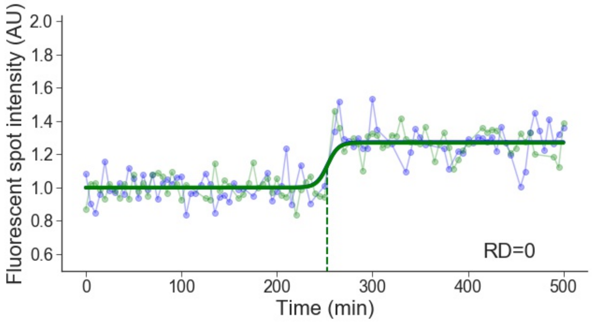


Cell line: Mid-late 1, cell: S42 cell 5

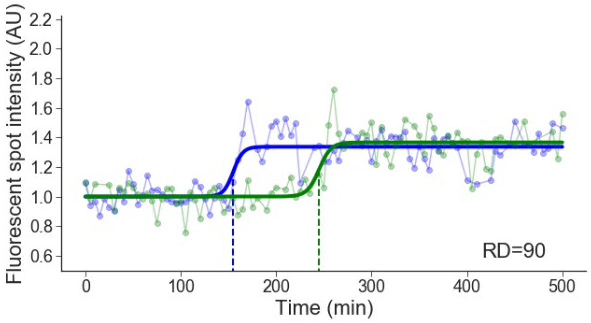

Cell line: Mid-late 1, cell: S43 cell 2

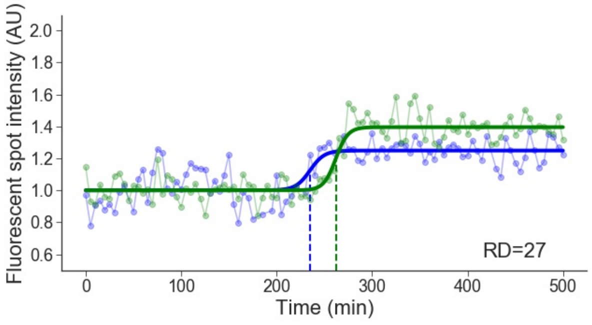


Cell line: Mid-late 1, cell: S46 cell 1

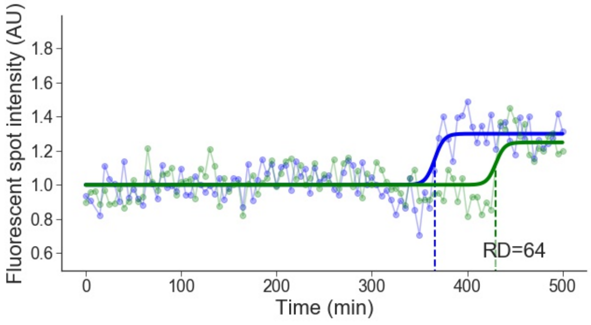


Cell line: Mid-late 1, cell: S46 cell 2

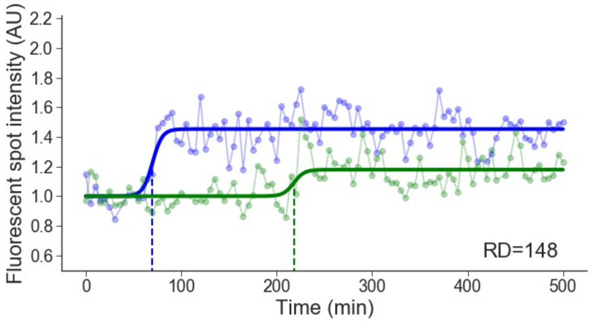


Cell line: Mid-late 1, cell: S47 cell 1

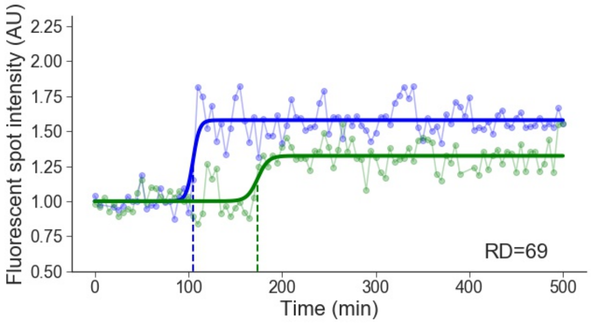


Cell line: Mid-late 1, cell: S48 cell 1

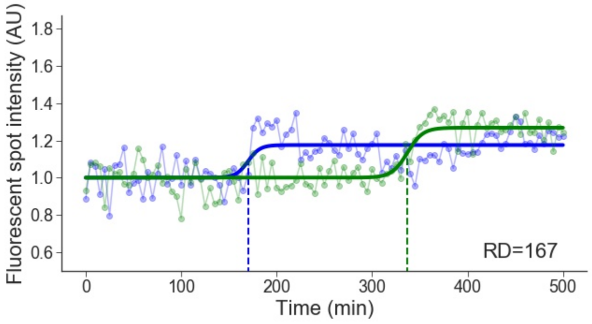


Cell line: Mid-late 1, cell: S48 cell 4

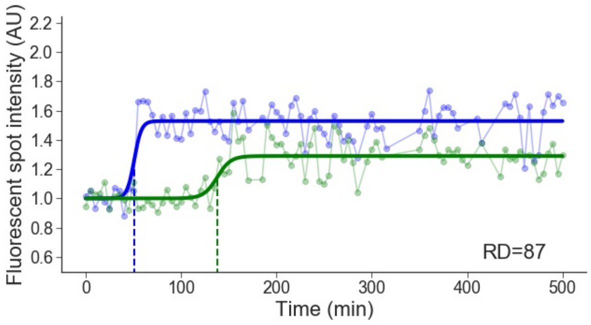


Cell line: Mid-late 1, cell: S51 cell 4

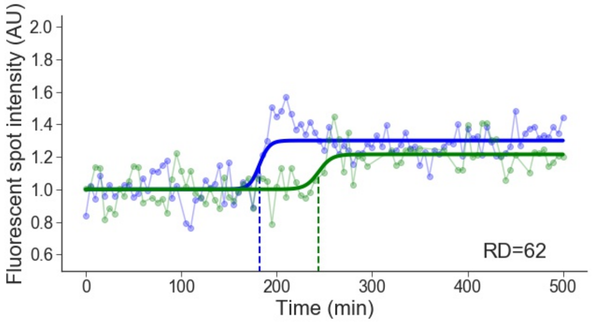


Cell line: Mid-late 1, cell: S53 cell 1

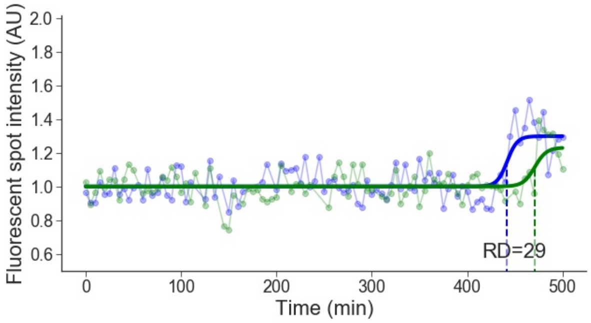

Cell line: Mid-late 1, cell: S53 cell 2

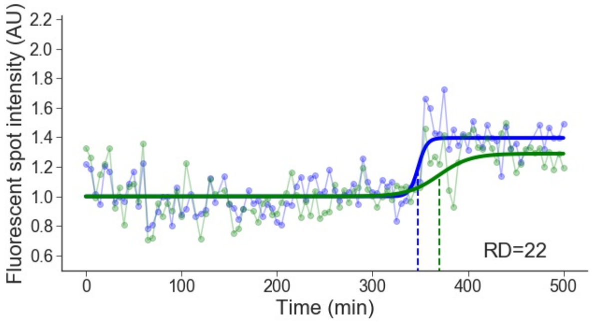


Cell line: Mid-late 1, cell: S57 cell 1

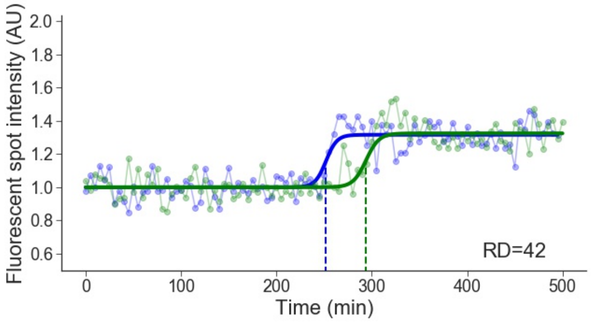


Cell line: Mid-late 1, cell: S57 cell 2

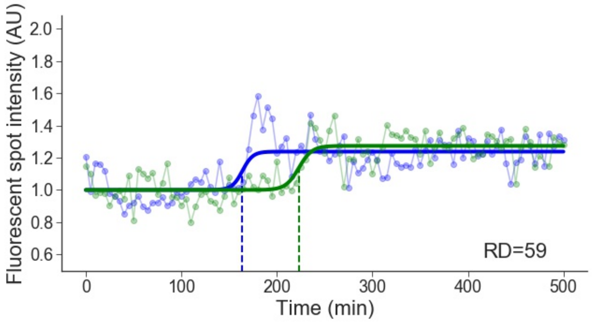

### 5 Mid-late 2

Cell line: Mid-late 2, cell: S1 cell 2

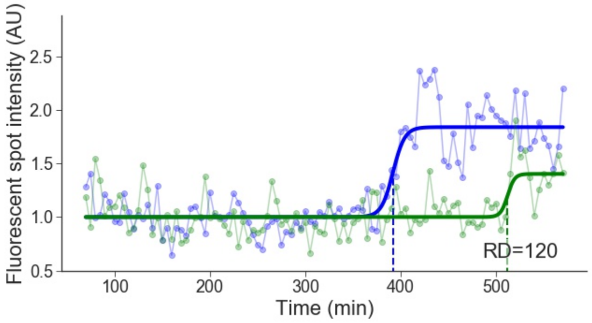


Cell line: Mid-late 2, cell: S1 cell 3

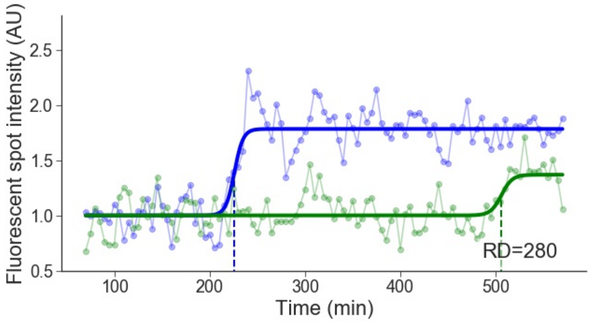


Cell line: Mid-late 2, cell: S3 cell 1

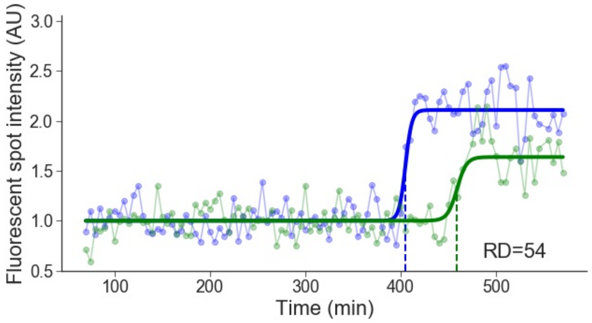


Cell line: Mid-late 2, cell: S3 cell 2

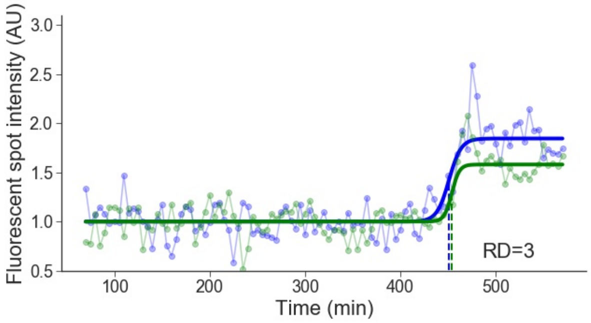


Cell line: Mid-late 2, cell: S3 cell 3

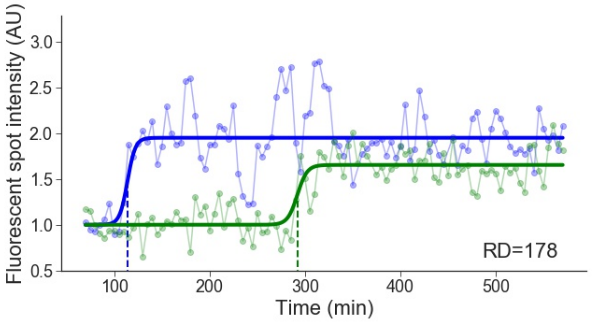


Cell line: Mid-late 2, cell: S5 cell 1

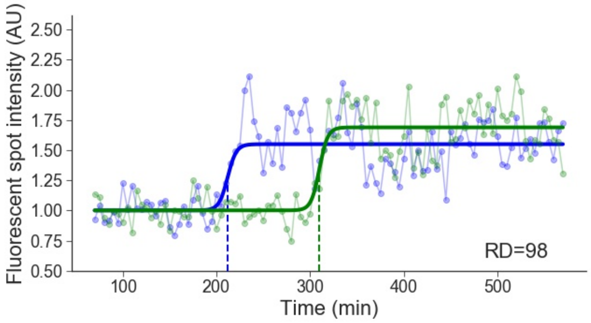


Cell line: Mid-late 2, cell: S6 cell 2

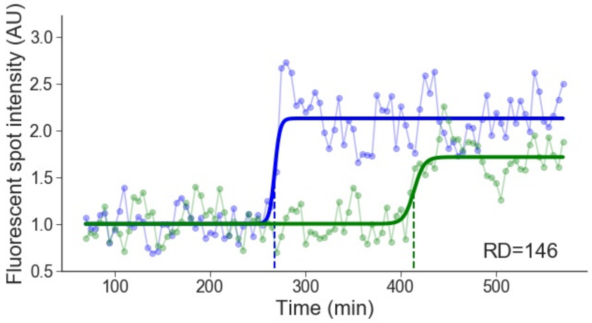


Cell line: Mid-late 2, cell: S7 cell 2

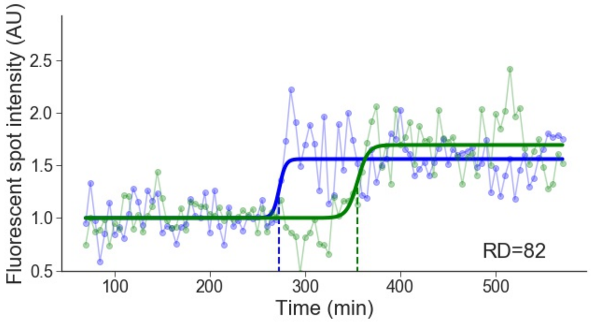

Cell line: Mid-late 2, cell: S7 cell 3

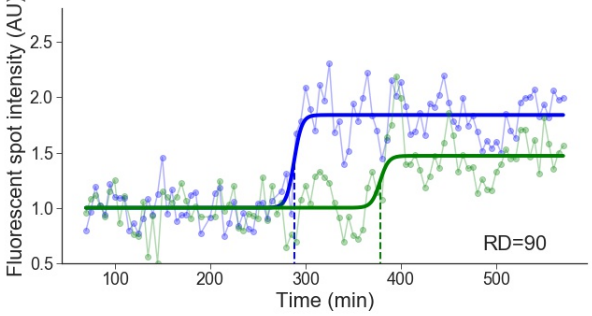


Cell line: Mid-late 2, cell: S8 cell 3

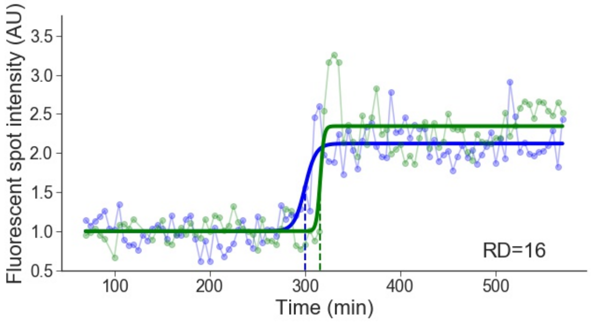


Cell line: Mid-late 2, cell: S8 cell 4

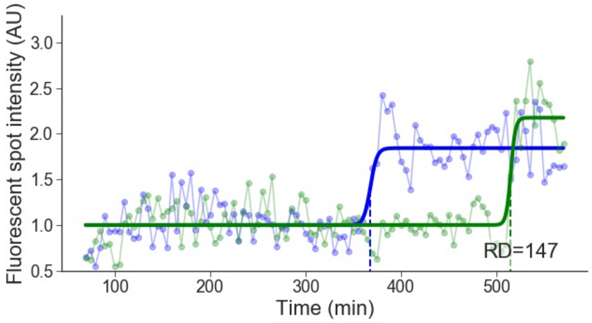


Cell line: Mid-late 2, cell: S8 cell 7

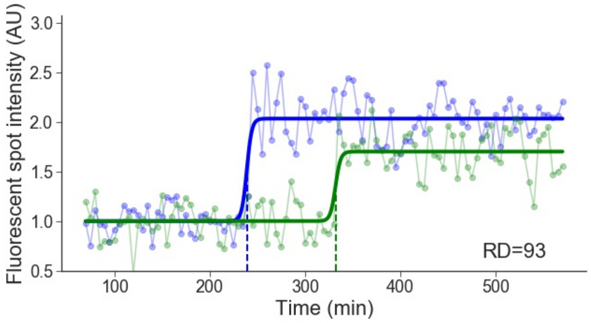


Cell line: Mid-late 2, cell: S11 cell 1

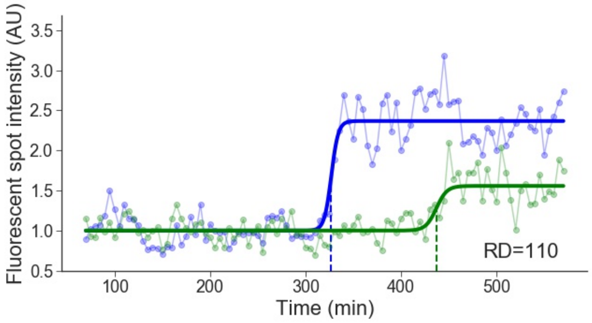


Cell line: Mid-late 2, cell: S12 cell 3

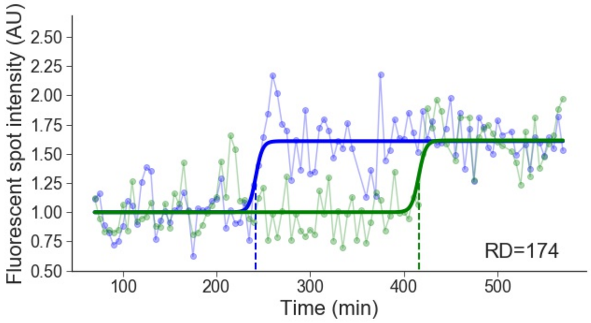


Cell line: Mid-late 2, cell: S13 cell 1

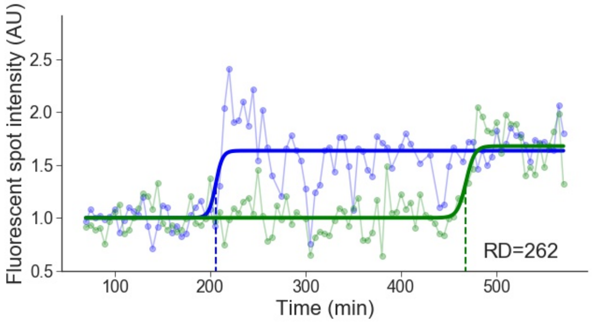


Cell line: Mid-late 2, cell: S14 cell 2

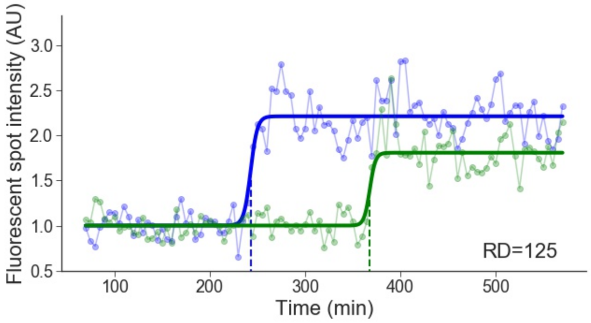

Cell line: Mid-late 2, cell: S15 cell 3

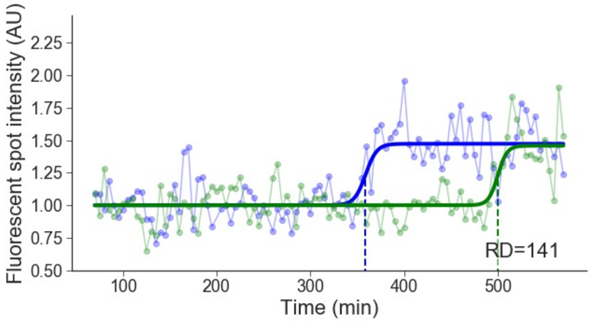


Cell line: Mid-late 2, cell: S15 cell 4

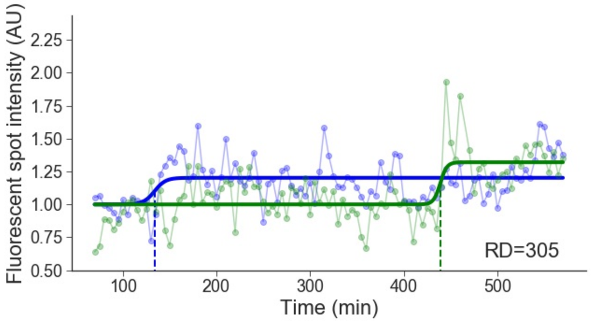


Cell line: Mid-late 2, cell: S17 cell 4

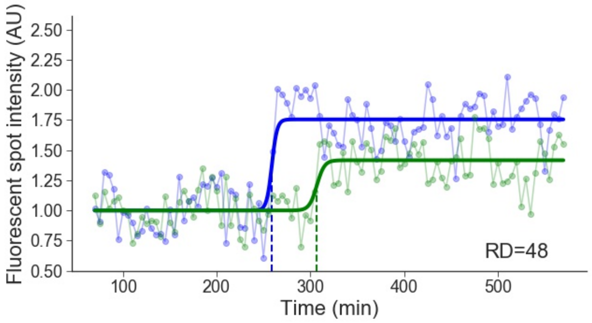


Cell line: Mid-late 2, cell: S18 cell 2

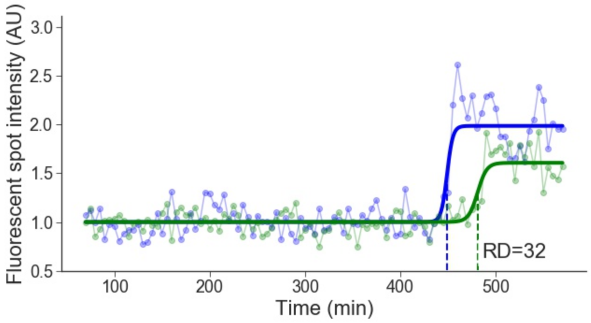


Cell line: Mid-late 2, cell: S19 cell 1

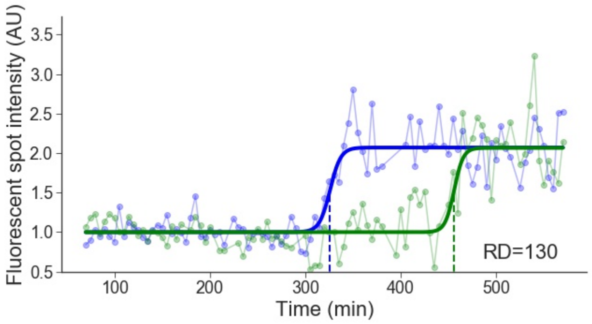


Cell line: Mid-late 2, cell: S20 cell 1

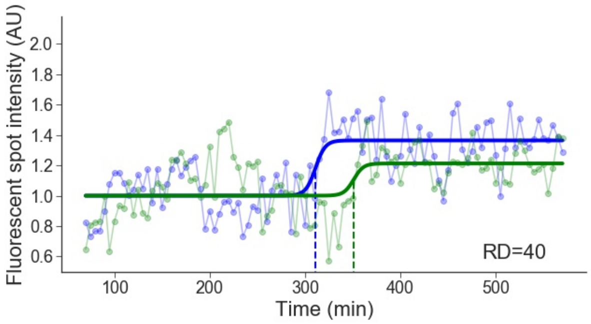


Cell line: Mid-late 2, cell: S20 cell 2

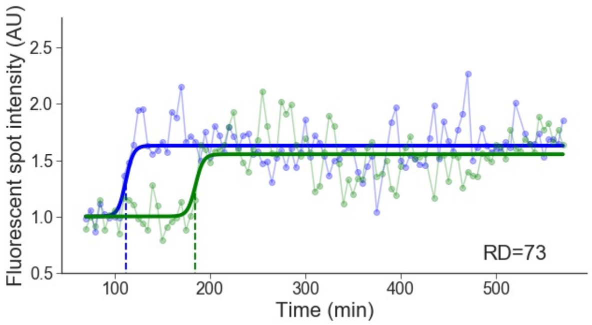


Cell line: Mid-late 2, cell: S21 cell 1

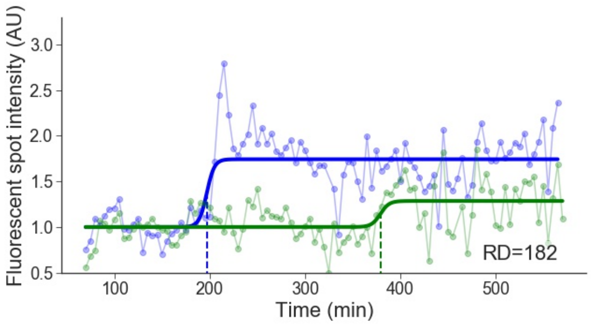

Cell line: Mid-late 2, cell: S22 cell 1

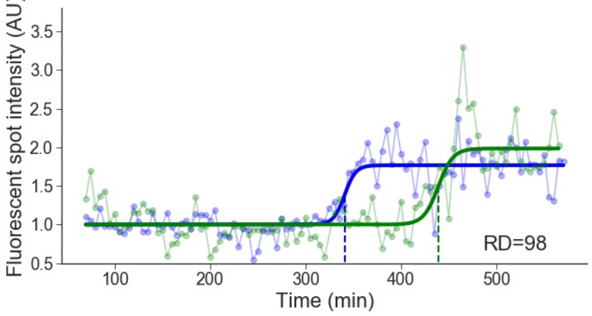


Cell line: Mid-late 2, cell: S23 cell 1

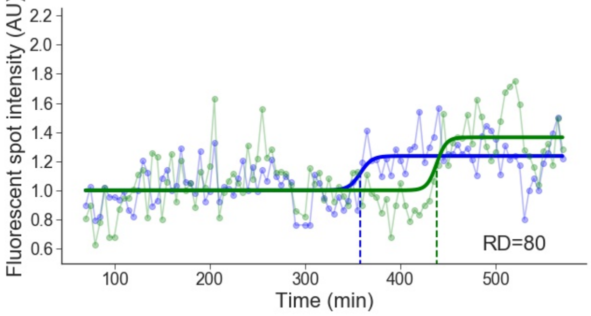


Cell line: Mid-late 2, cell: S23 cell 3

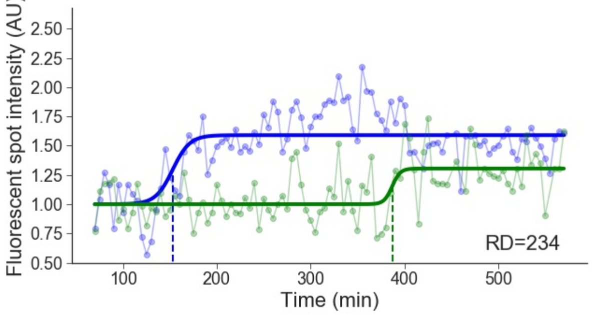


Cell line: Mid-late 2, cell: S24 cell 1

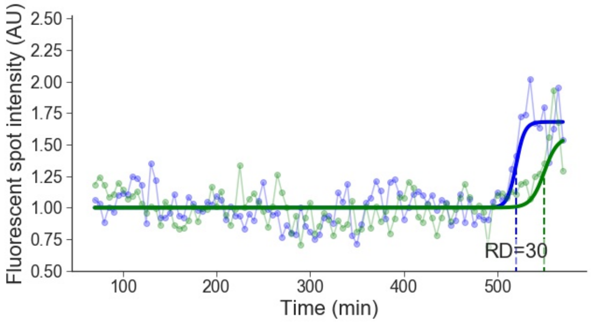


Cell line: Mid-late 2, cell: S25 cell 1

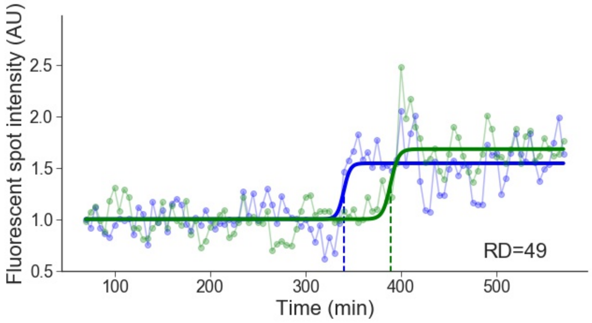


Cell line: Mid-late 2, cell: S25 cell 2

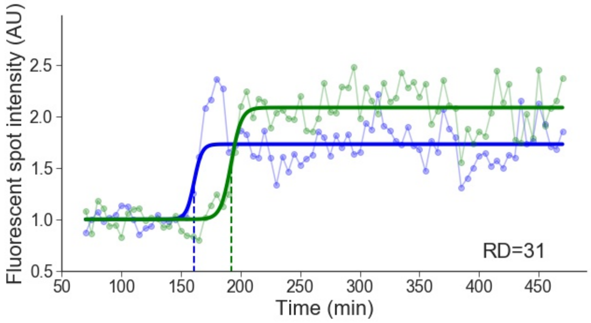


Cell line: Mid-late 2, cell: S26 cell 1

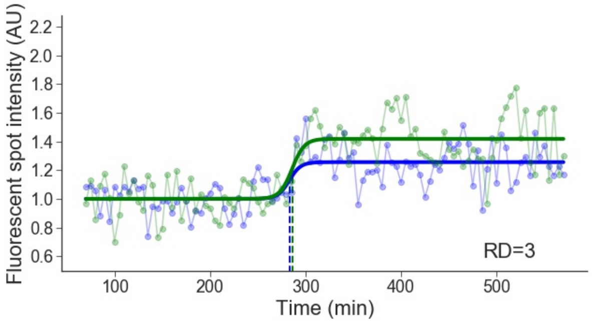


Cell line: Mid-late 2, cell: S26 cell 2

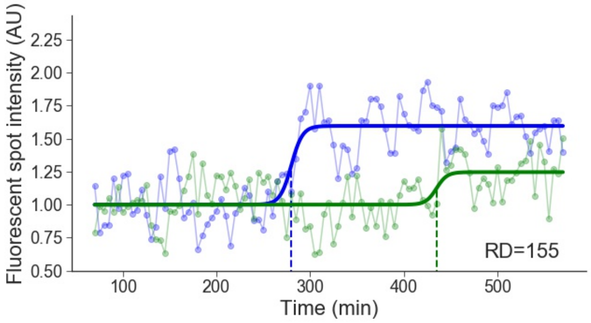

Cell line: Mid-late 2, cell: S28 cell 2

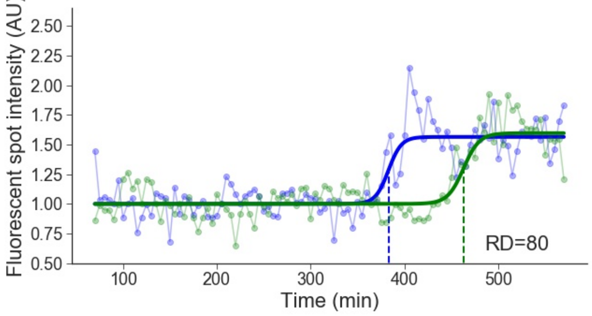


Cell line: Mid-late 2, cell: S28 cell 3

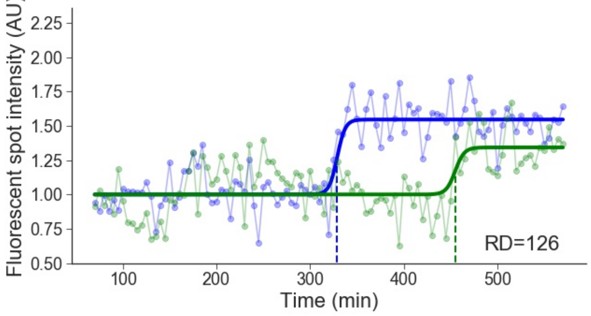


Cell line: Mid-late 2, cell: S30 cell 1

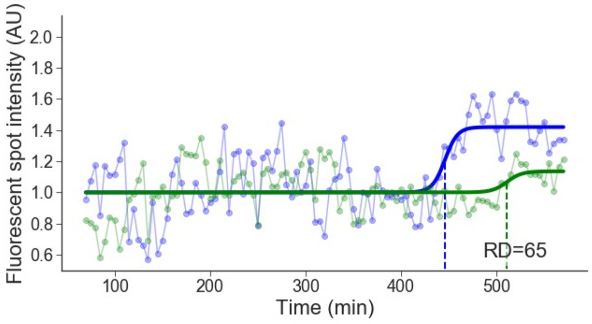


Cell line: Mid-late 2, cell: S30 cell 2

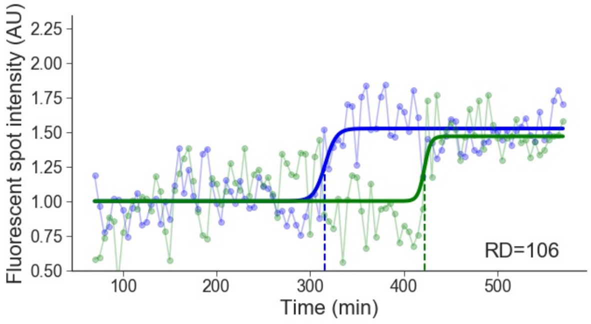


Cell line: Mid-late 2, cell: S30 cell 3

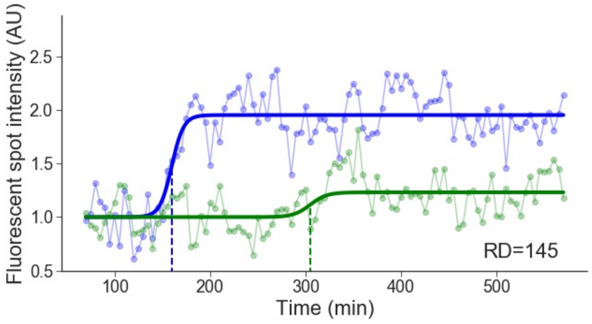


Cell line: Mid-late 2, cell: S30 cell 5

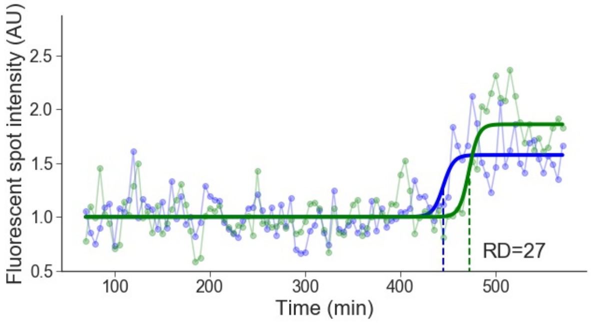


Cell line: Mid-late 2, cell: S32 cell 2

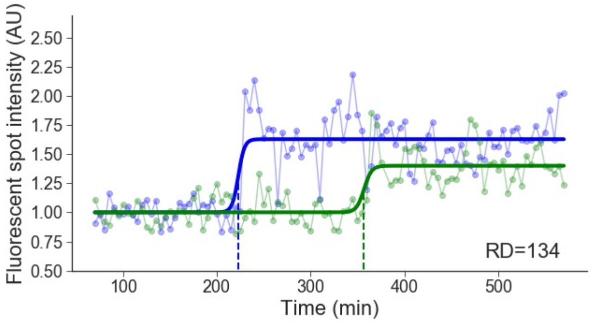


Cell line: Mid-late 2, cell: S33 cell 3

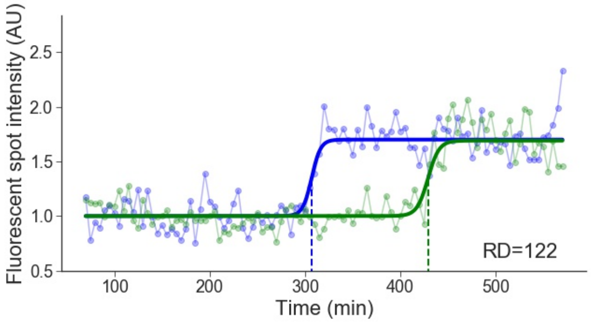

Cell line: Mid-late 2, cell: S34 cell 4

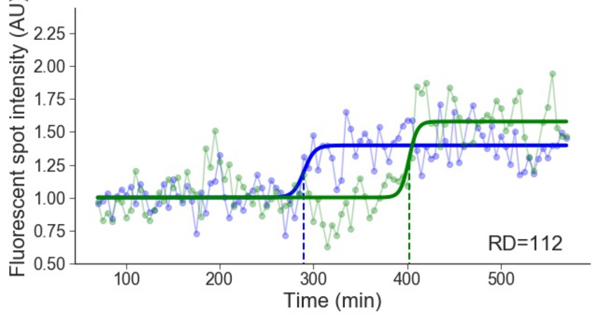


Cell line: Mid-late 2, cell: S35 cell 3

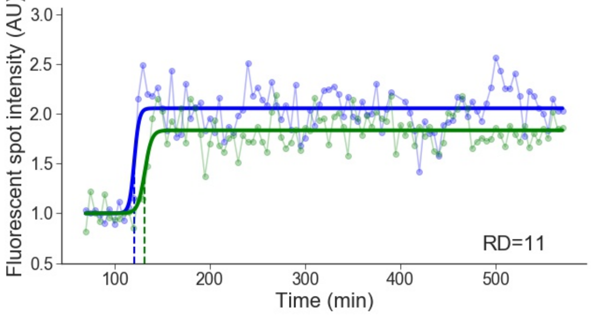


Cell line: Mid-late 2, cell: S35 cell 4

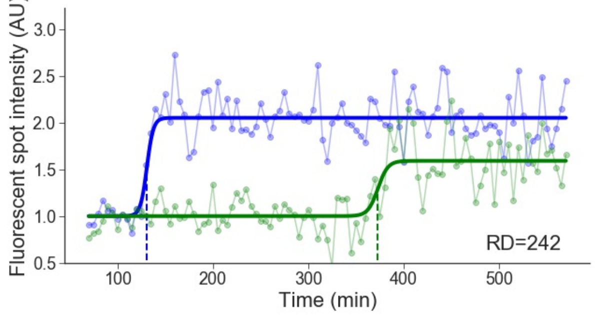


Cell line: Mid-late 2, cell: S36 cell 2

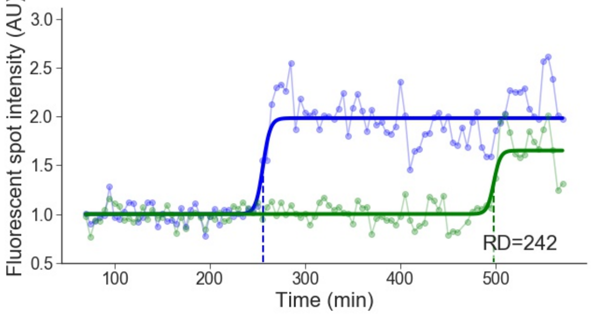


Cell line: Mid-late 2, cell: S36 cell 3

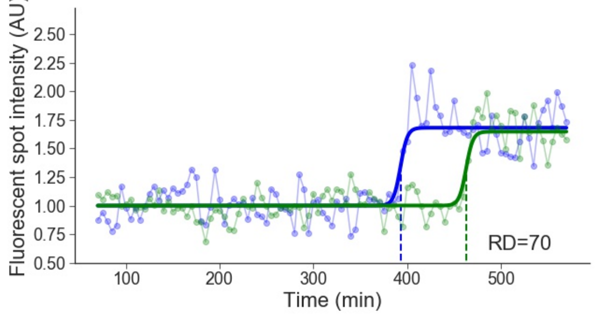


Cell line: Mid-late 2, cell: S38 cell 3

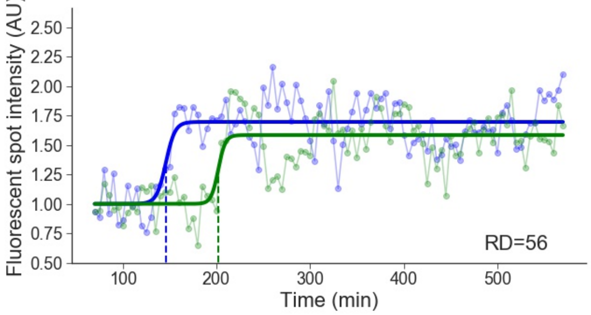


Cell line: Mid-late 2, cell: S39 cell 2

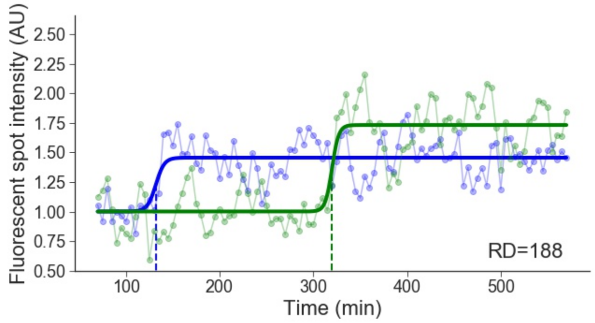


Cell line: Mid-late 2, cell: S39 cell 5

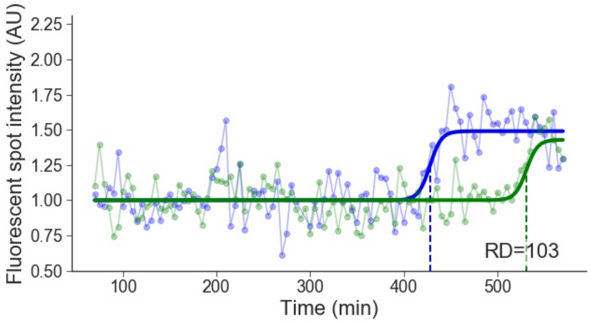

Cell line: Mid-late 2, cell: S39 cell 6

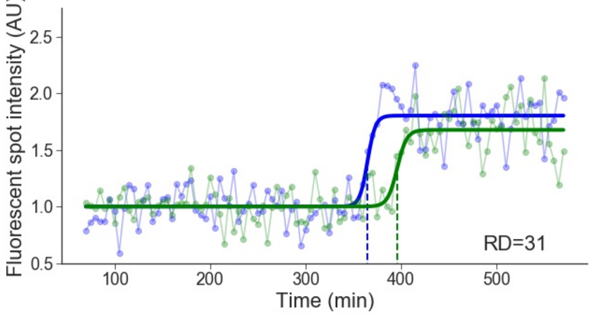


Cell line: Mid-late 2, cell: S40 cell 1

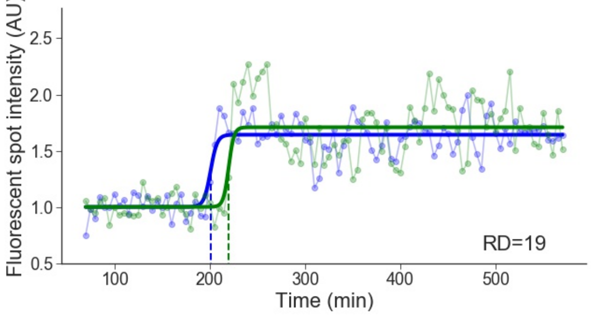


Cell line: Mid-late 2, cell: S40 cell 2

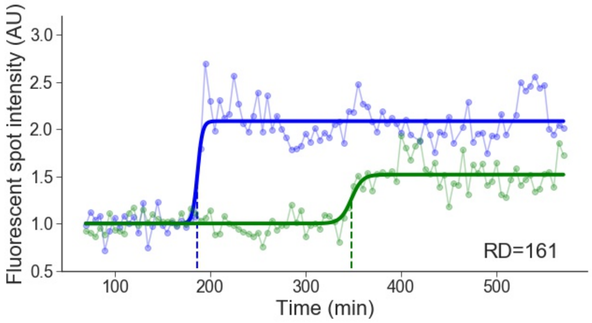


Cell line: Mid-late 2, cell: S40 cell 5

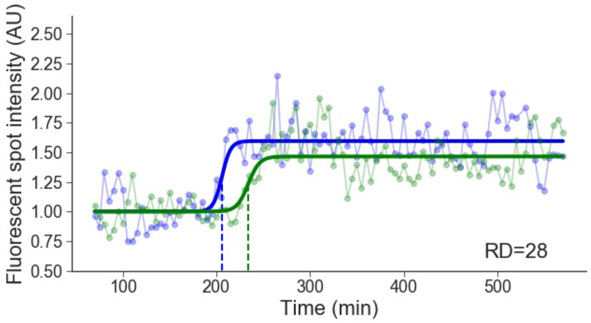


Cell line: Mid-late 2, cell: S41 cell 1

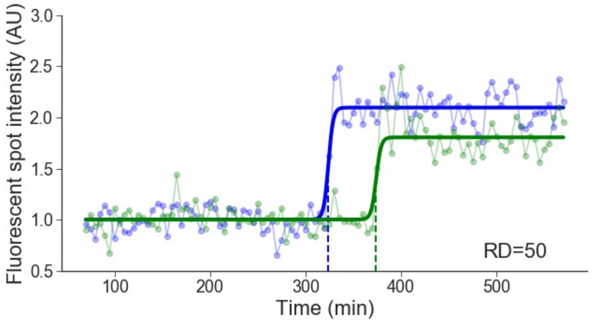


Cell line: Mid-late 2, cell: S41 cell 3

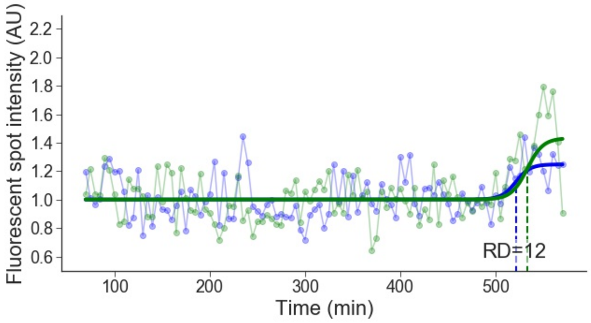


Cell line: Mid-late 2, cell: S42 cell 1

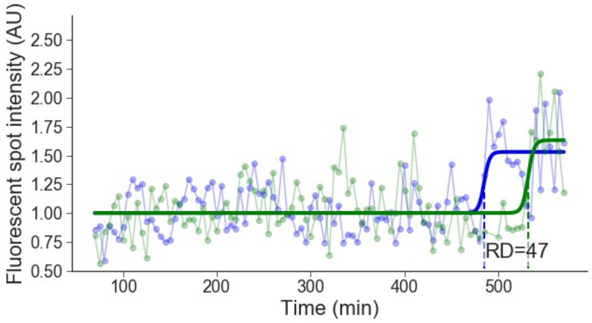

### 6 Late 1

Cell line: Late 1, cell: S1 cell 1

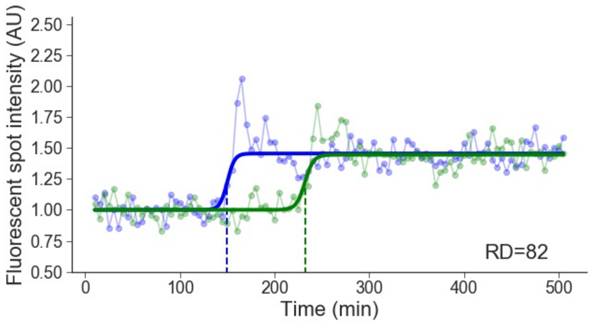


Cell line: Late 1, cell: S1 cell 2

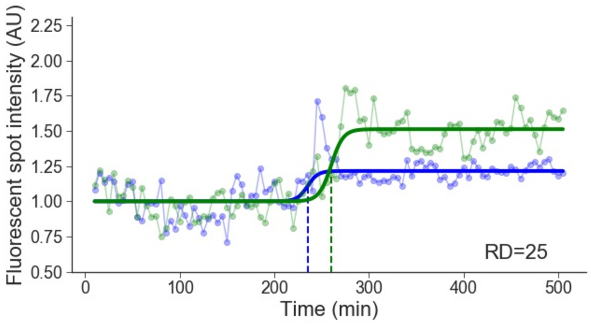


Cell line: Late 1, cell: S3 cell 1

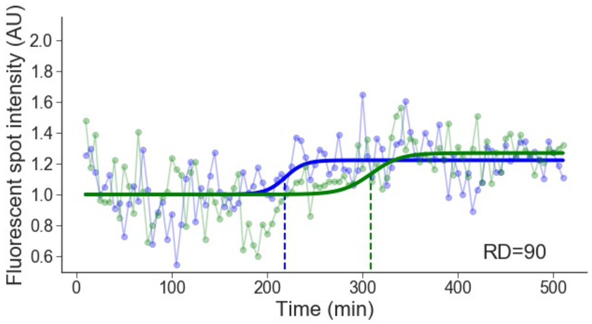


Cell line: Late 1, cell: S3 cell 2

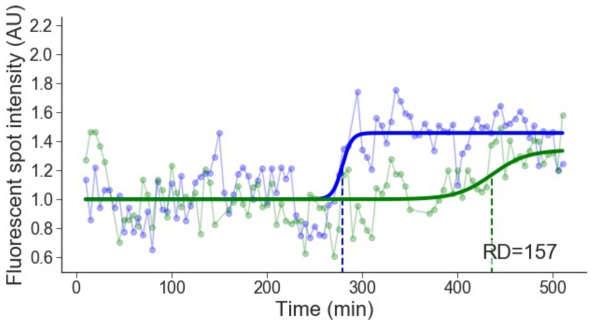


Cell line: Late 1, cell: S4 cell 1

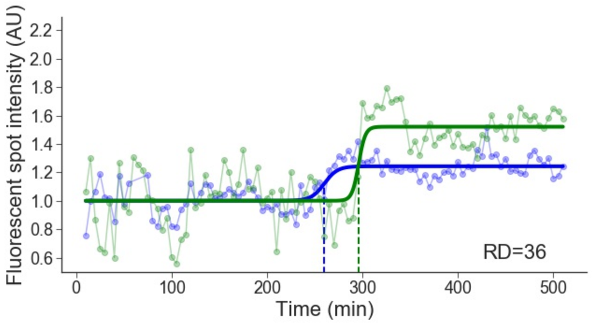


Cell line: Late 1, cell: S4 cell 2

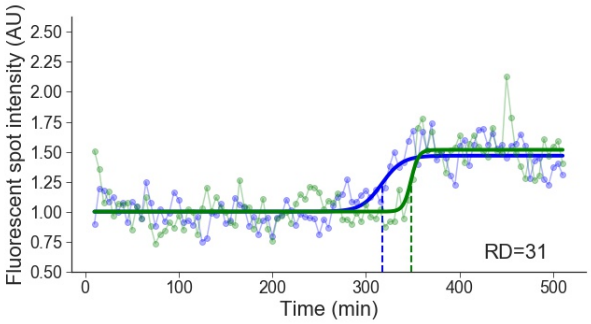


Cell line: Late 1, cell: S4 cell 4

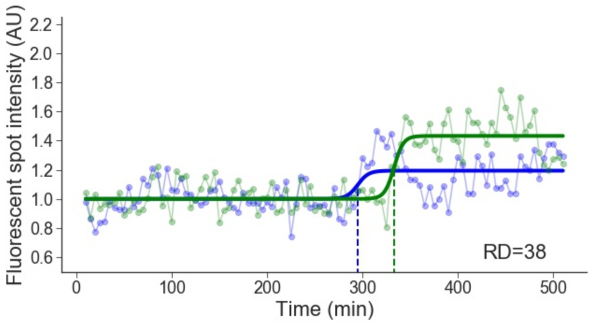


Cell line: Late 1, cell: S5 cell 1

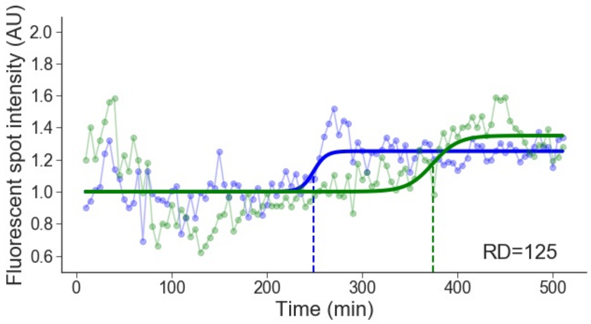

Cell line: Late 1, cell: S5 cell 2

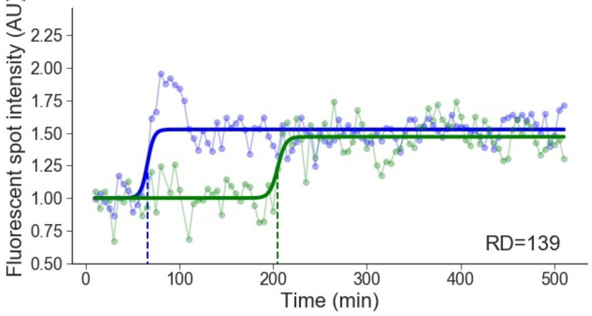


Cell line: Late 1, cell: S5 cell 4

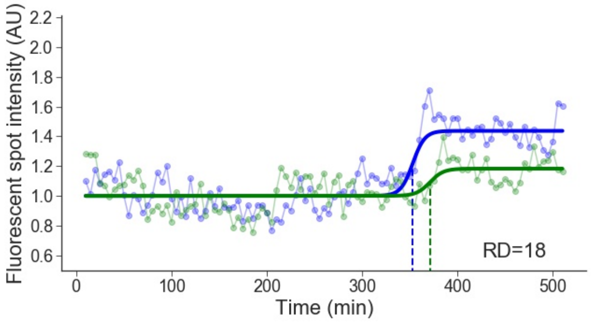


Cell line: Late 1, cell: S6 cell 1

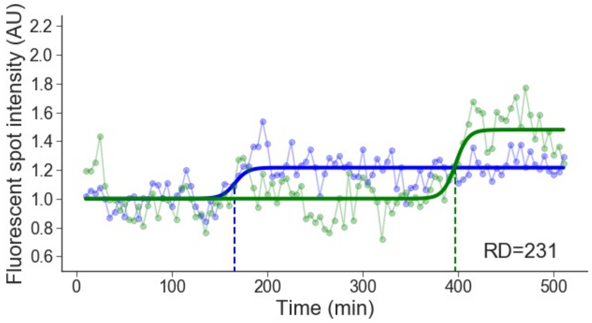


Cell line: Late 1, cell: S6 cell 2

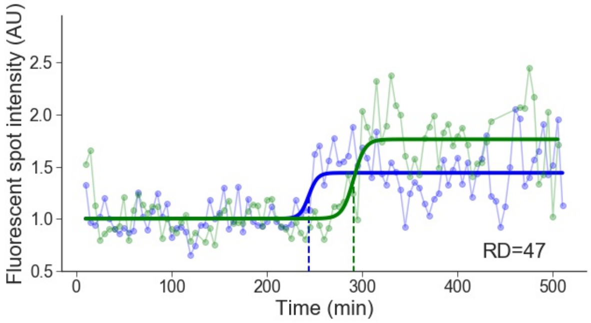


Cell line: Late 1, cell: S6 cell 3

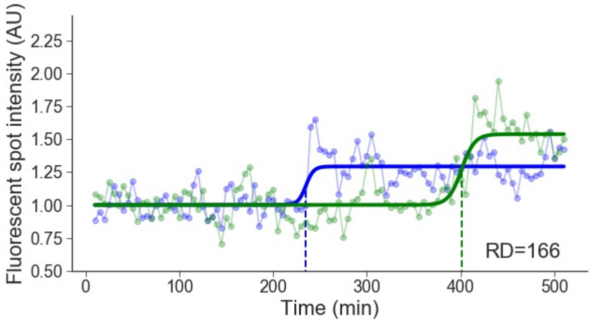


Cell line: Late 1, cell: S6 cell 5

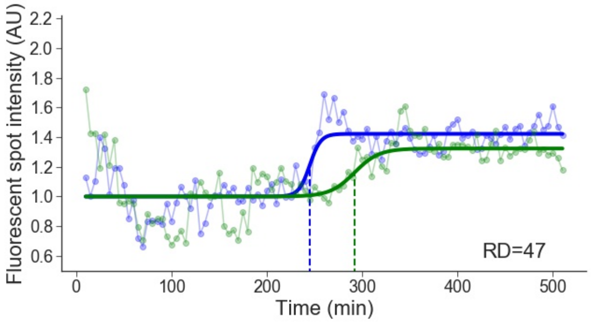


Cell line: Late 1, cell: S8 cell 1

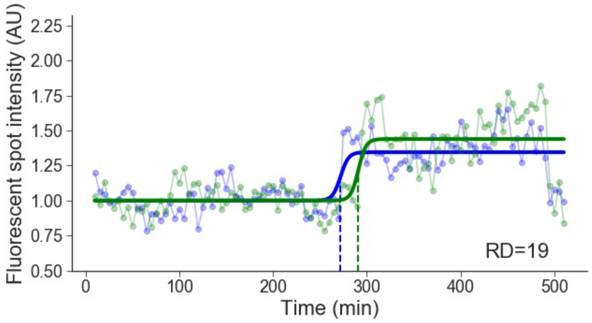


Cell line: Late 1, cell: S8 cell 2

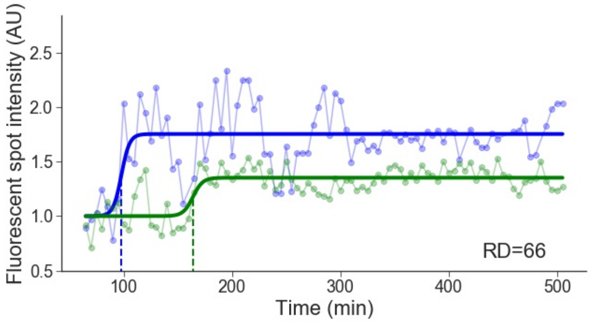

Cell line: Late 1, cell: S8 cell 3

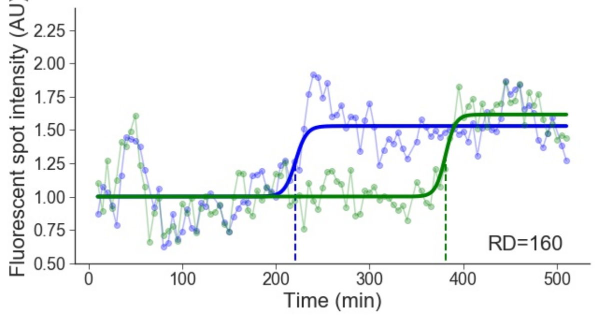


Cell line: Late 1, cell: S9 cell 1

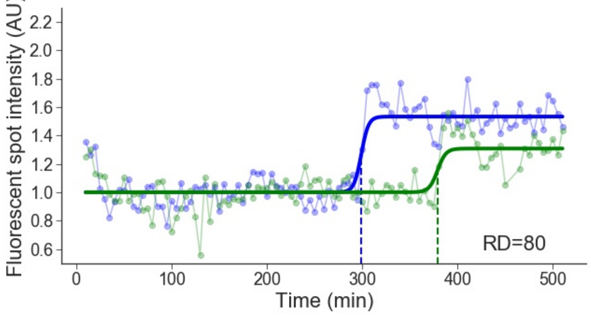


Cell line: Late 1, cell: S9 cell 2

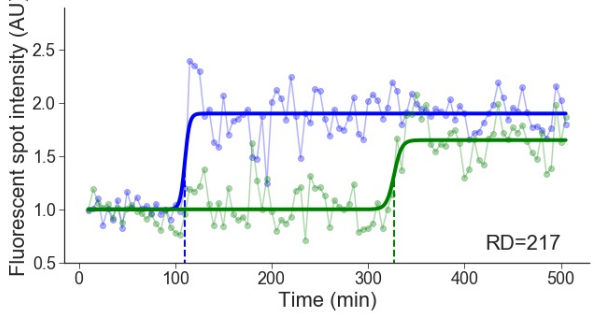


Cell line: Late 1, cell: S10 cell 1

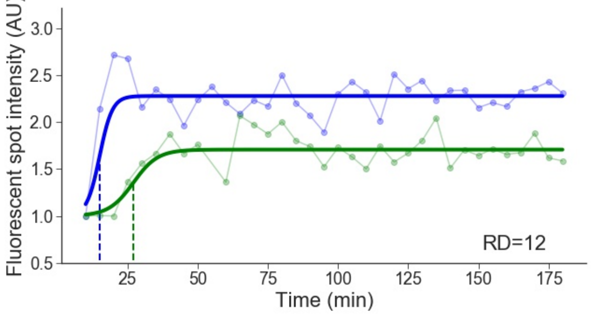


Cell line: Late 1, cell: S11 cell 2

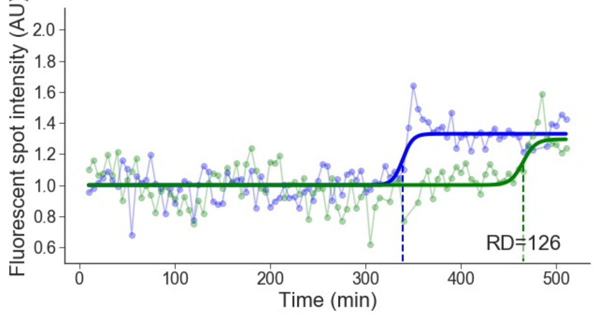


Cell line: Late 1, cell: S11 cell 3

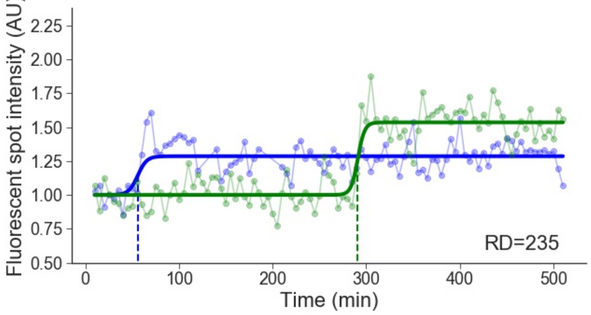


Cell line: Late 1, cell: S12 cell 1

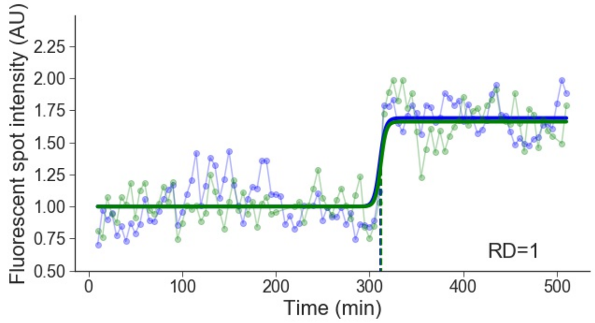


Cell line: Late 1, cell: S13 cell 1

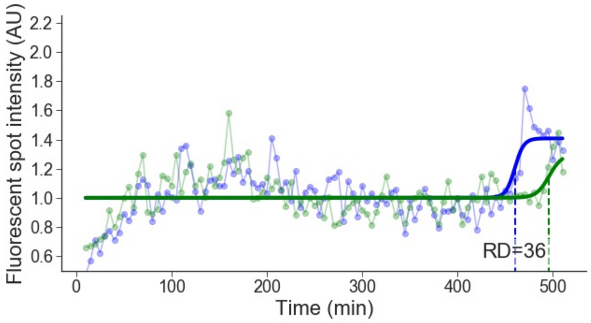

Cell line: Late 1, cell: S14 cell 2

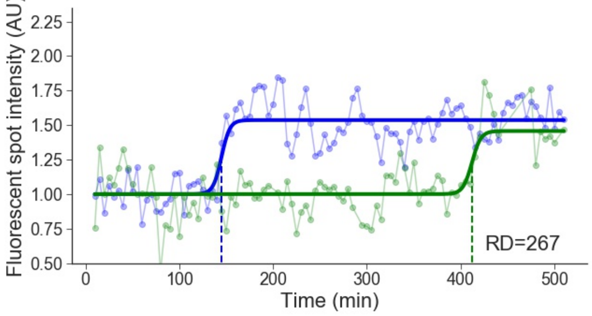


Cell line: Late 1, cell: S15 cell 1

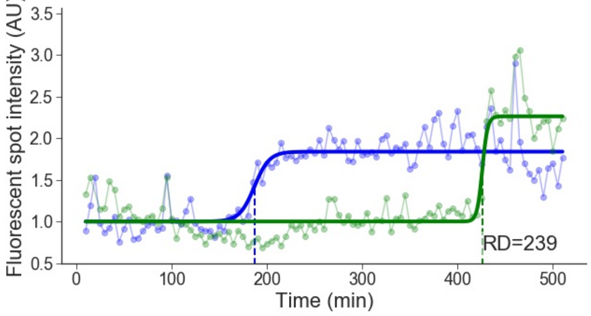


Cell line: Late 1, cell: S15 cell 2

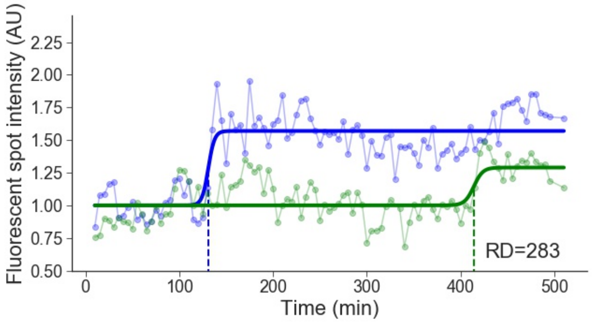


Cell line: Late 1, cell: S15 cell 3

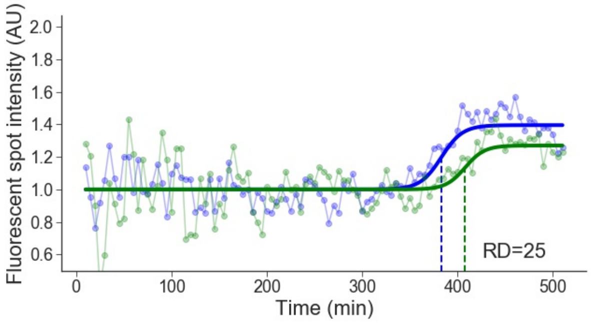


Cell line: Late 1, cell: S16 cell 1

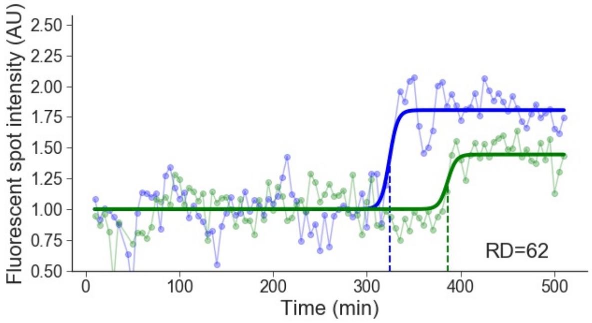


Cell line: Late 1, cell: S16 cell 2

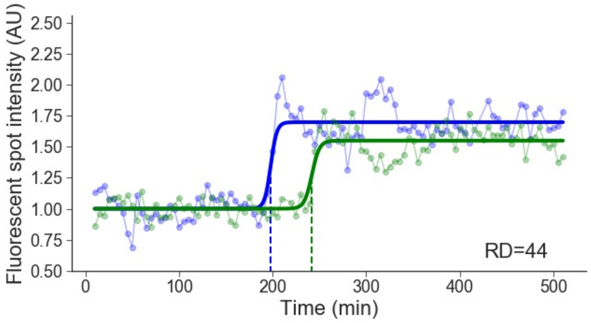


Cell line: Late 1, cell: S16 cell 3

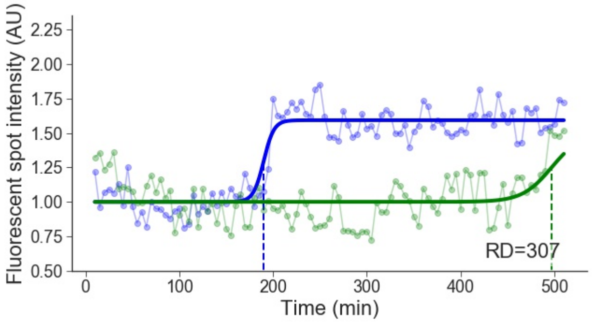


Cell line: Late 1, cell: S16 cell 4

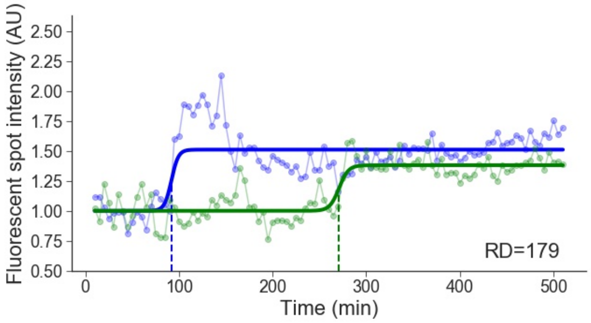

Cell line: Late 1, cell: S17 cell 1

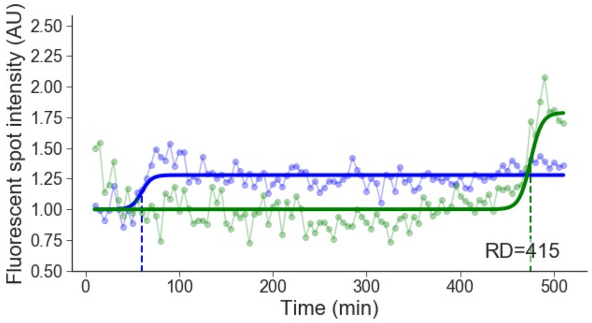


Cell line: Late 1, cell: S17 cell 2

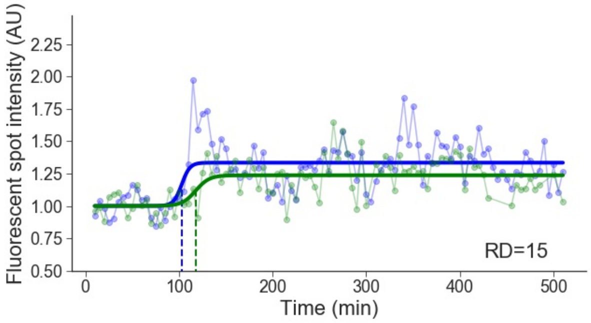


Cell line: Late 1, cell: S17 cell 3

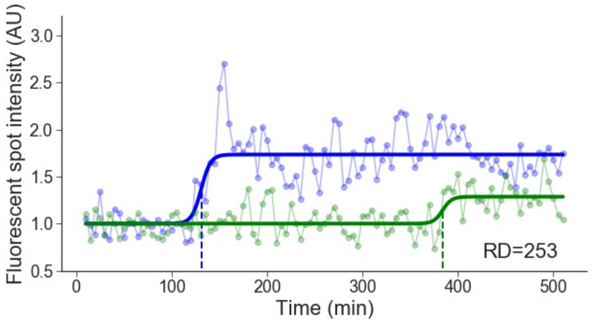


Cell line: Late 1, cell: S17 cell 4

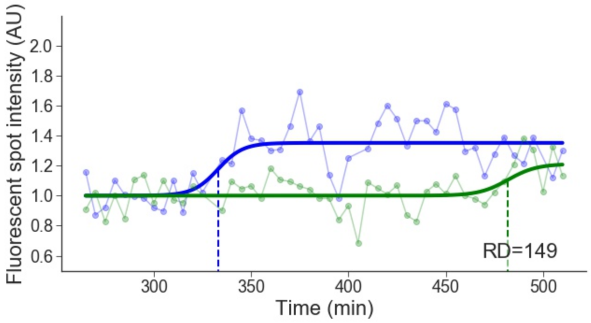


Cell line: Late 1, cell: S17 cell 6

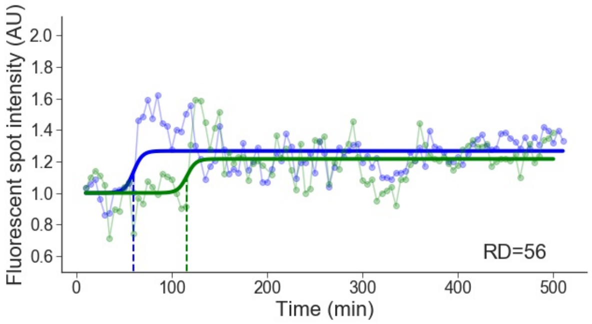


Cell line: Late 1, cell: S18 cell 1

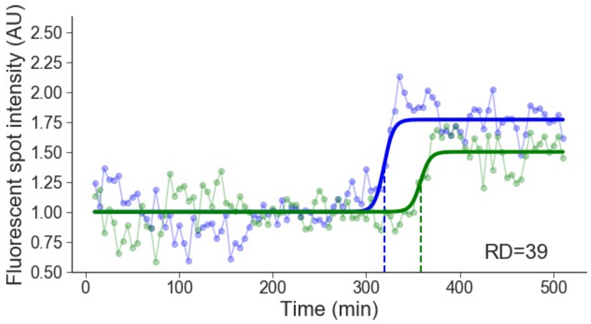


Cell line: Late 1, cell: S19 cell 1

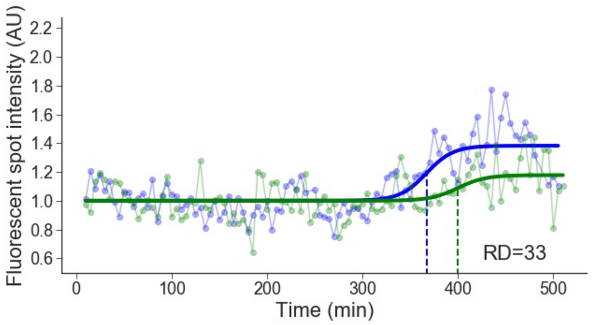


Cell line: Late 1, cell: S19 cell 2

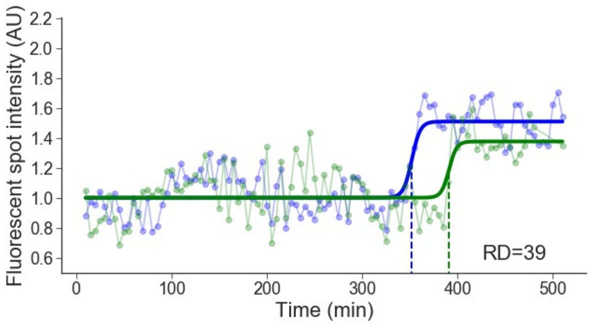

Cell line: Late 1, cell: S19 cell 3

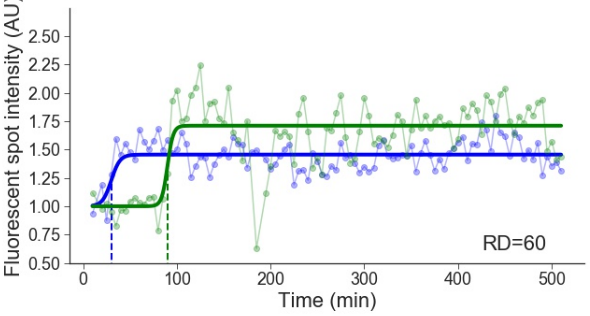


Cell line: Late 1, cell: S19 cell 4

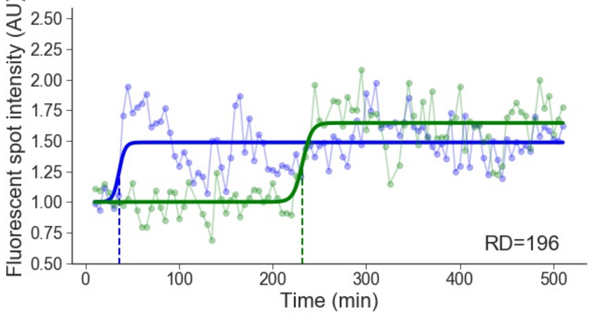


Cell line: Late 1, cell: S20 cell 2

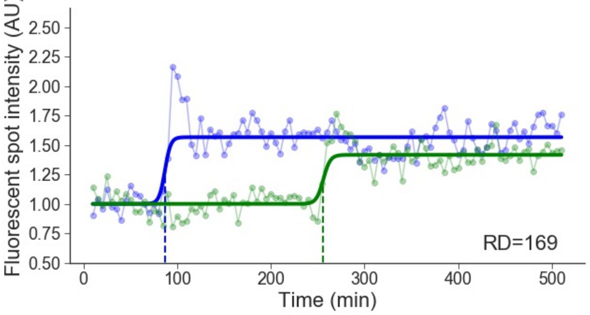


Cell line: Late 1, cell: S21 cell 1

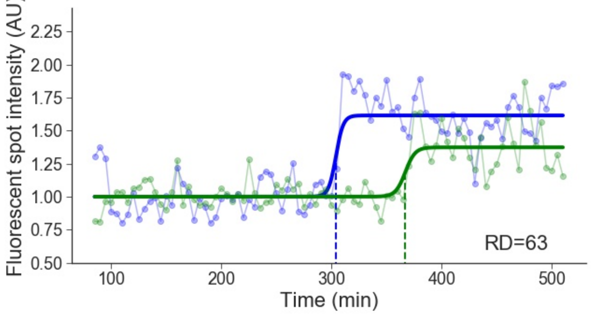


Cell line: Late 1, cell: S21 cell 2

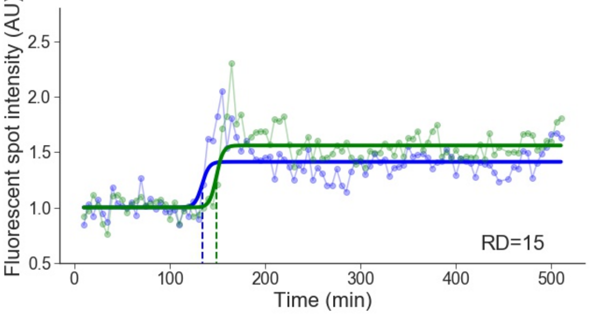


Cell line: Late 1, cell: S23 cell 1

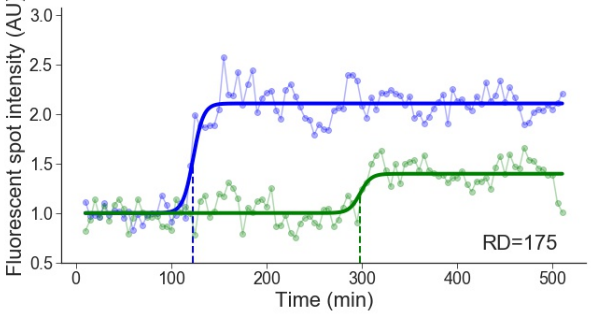


Cell line: Late 1, cell: S24 cell 1

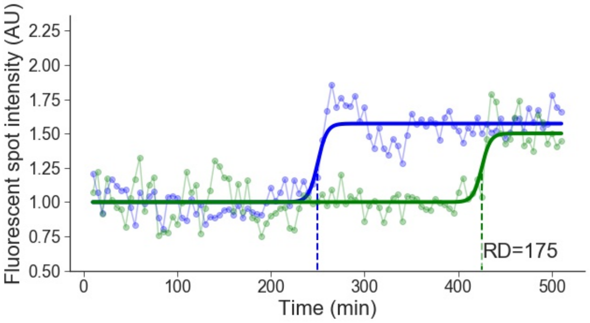


Cell line: Late 1, cell: S24 cell 2

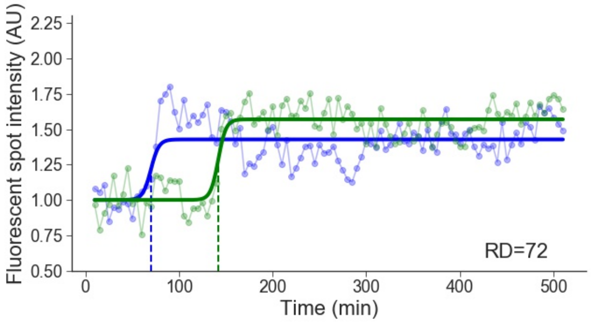

Cell line: Late 1, cell: S24 cell 3

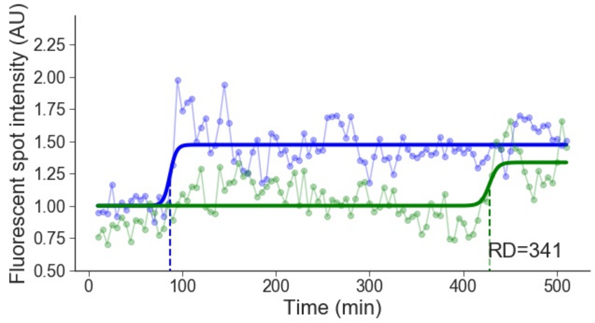


Cell line: Late 1, cell: S24 cell 4

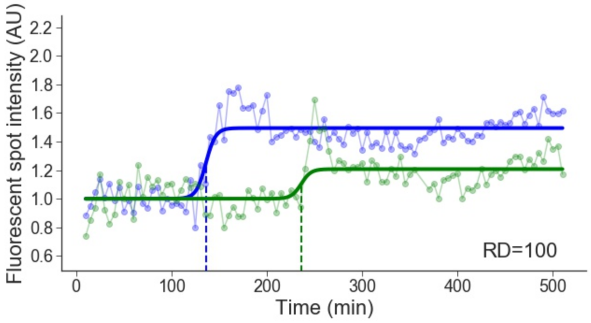

### 7 Late 2

Cell line: Late 2, cell: S1 cell 1

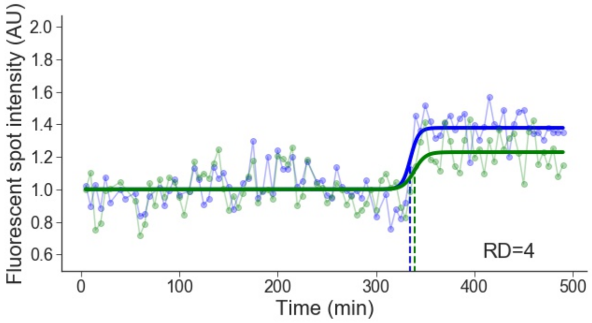


Cell line: Late 2, cell: S6 cell 1

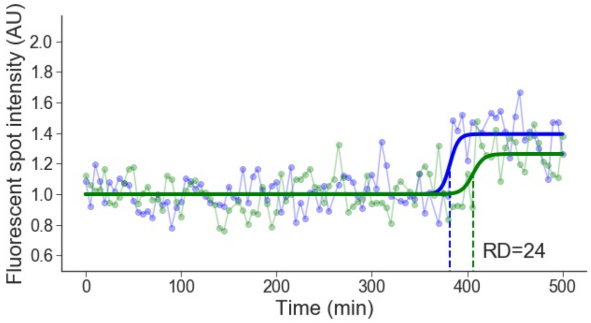


Cell line: Late 2, cell: S8 cell 1

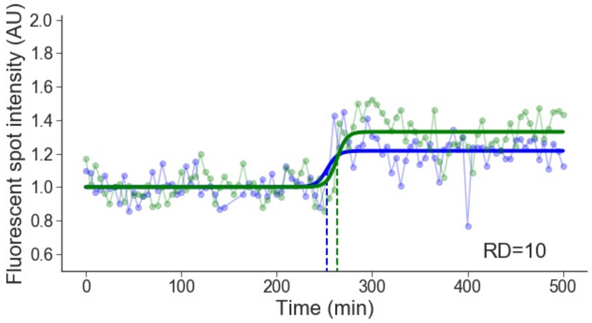


Cell line: Late 2, cell: S9 cell 2

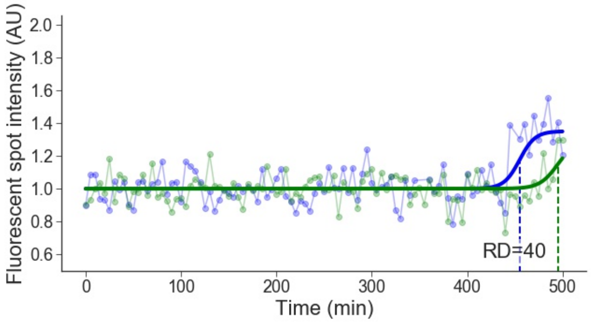


Cell line: Late 2, cell: S10 cell 1

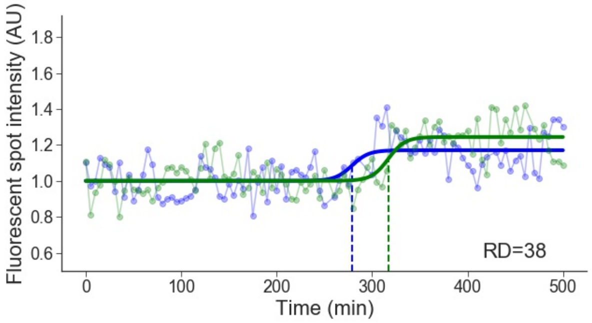


Cell line: Late 2, cell: S10 cell 2

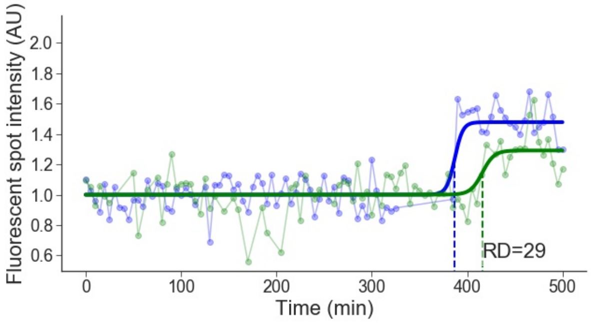


Cell line: Late 2, cell: S10 cell 3

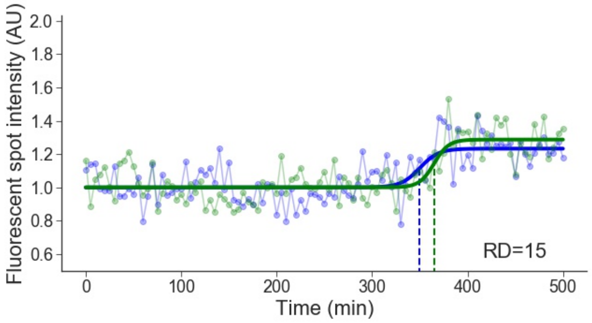


Cell line: Late 2, cell: S10 cell 4

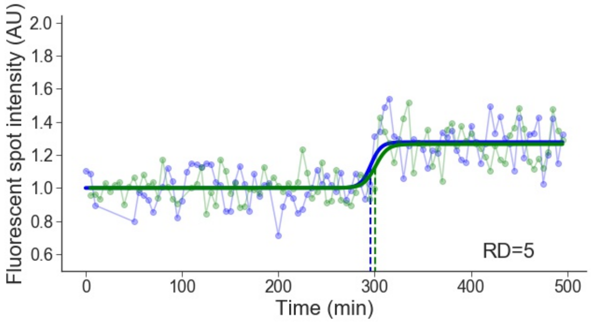

Cell line: Late 2, cell: S11 cell 1

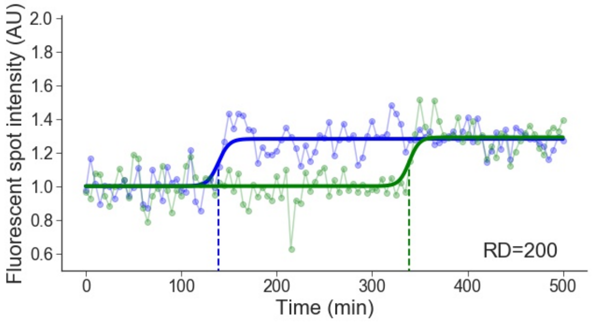


Cell line: Late 2, cell: S11 cell 2

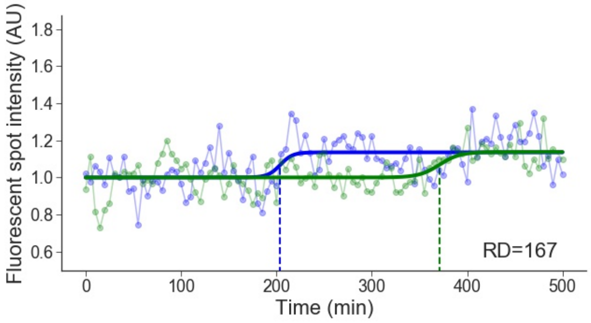


Cell line: Late 2, cell: S12 cell 1

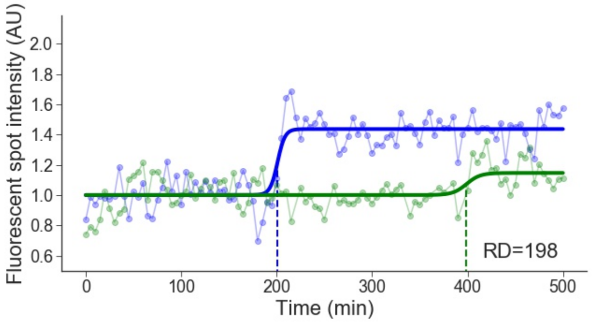


Cell line: Late 2, cell: S12 cell 2

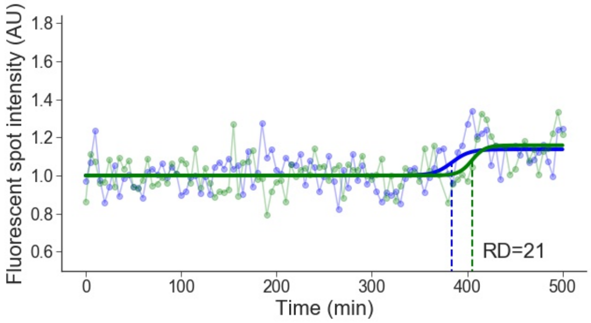


Cell line: Late 2, cell: S13 cell 1

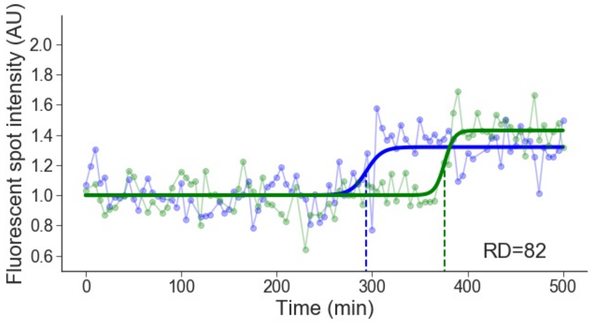


Cell line: Late 2, cell: S13 cell 2

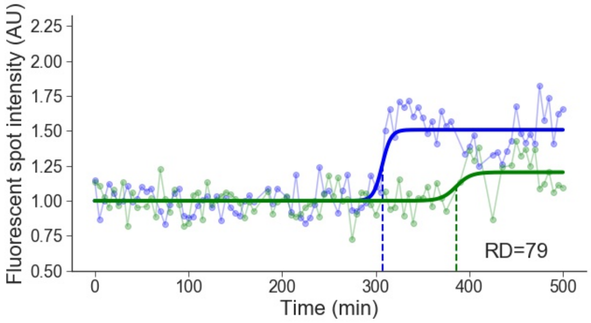


Cell line: Late 2, cell: S14 cell 1

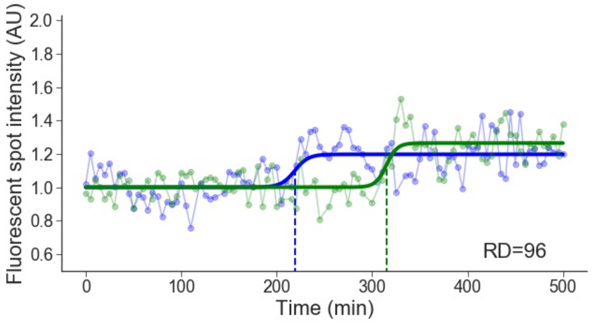


Cell line: Late 2, cell: S16 cell 1

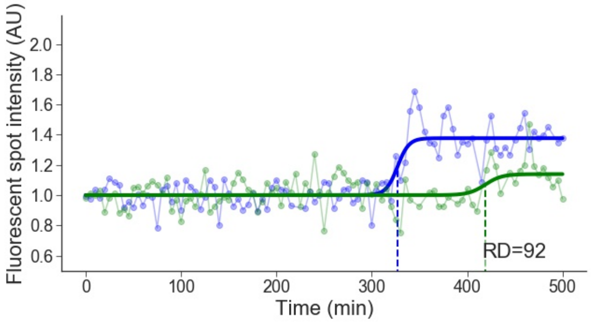

Cell line: Late 2, cell: S17 cell 1

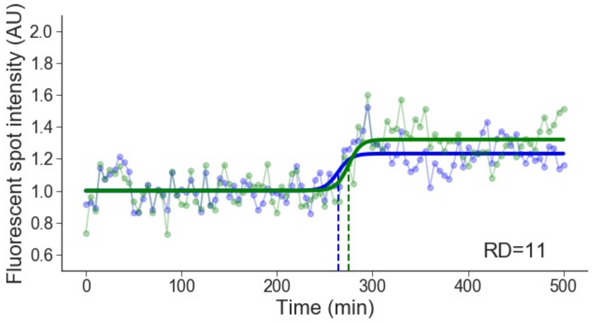


Cell line: Late 2, cell: S17 cell 2

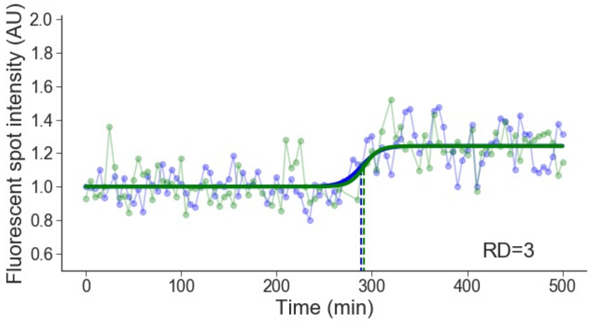


Cell line: Late 2, cell: S17 cell 3

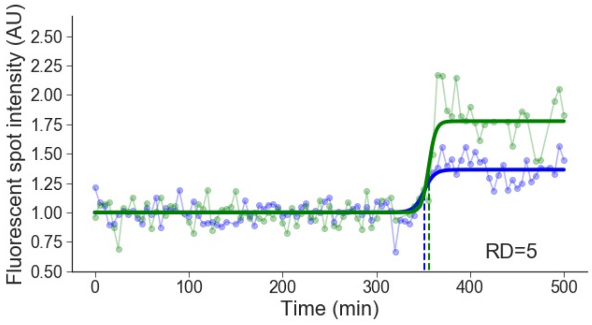


Cell line: Late 2, cell: S19 cell 1

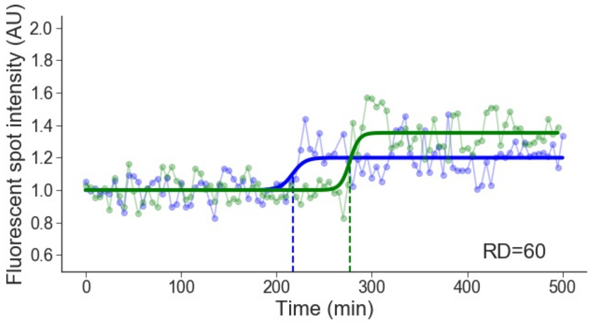


Cell line: Late 2, cell: S19 cell 2

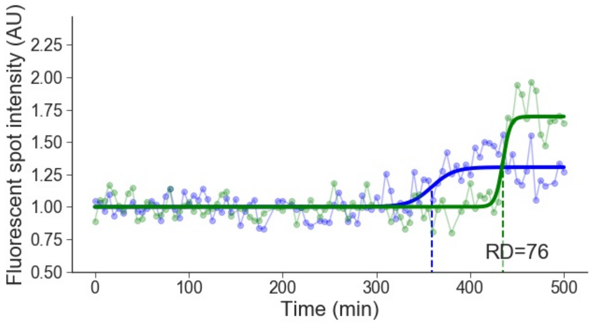


Cell line: Late 2, cell: S21 cell 1

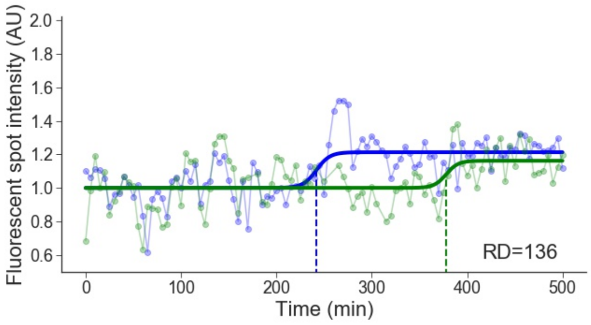


Cell line: Late 2, cell: S22 cell 4

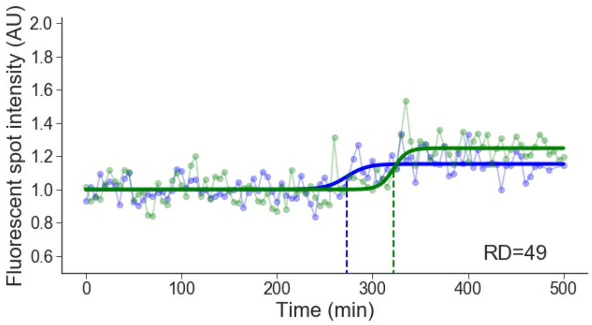


Cell line: Late 2, cell: S23 cell 1

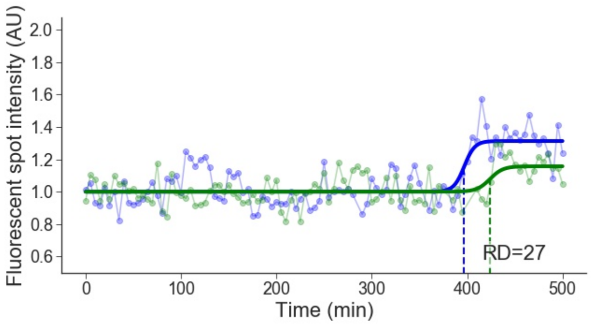

Cell line: Late 2, cell: S23 cell 2

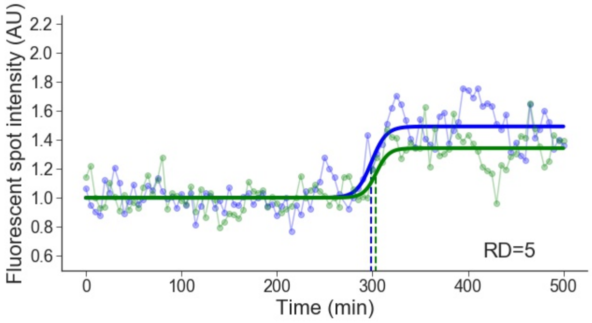


Cell line: Late 2, cell: S23 cell 3

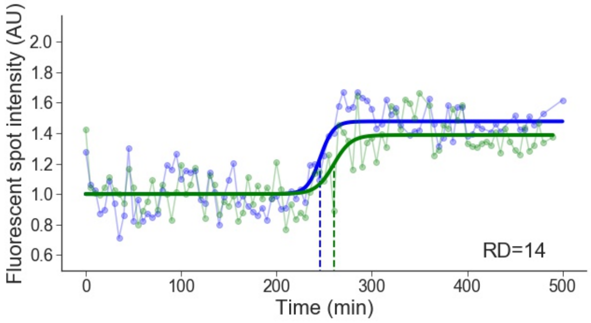


Cell line: Late 2, cell: S23 cell 4

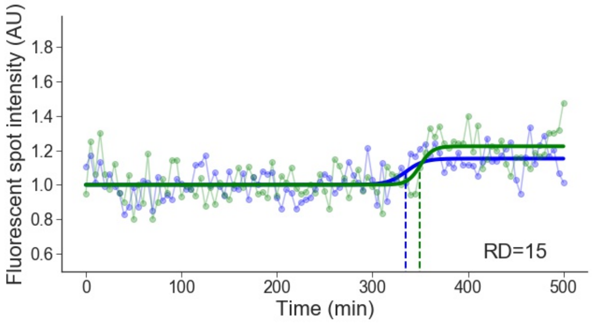


Cell line: Late 2, cell: S24 cell 1

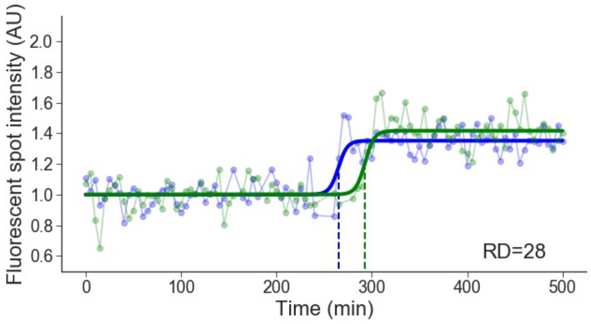


Cell line: Late 2, cell: S27 cell 1

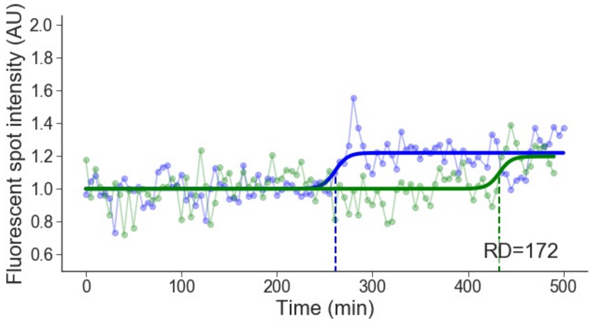


Cell line: Late 2, cell: S27 cell 2

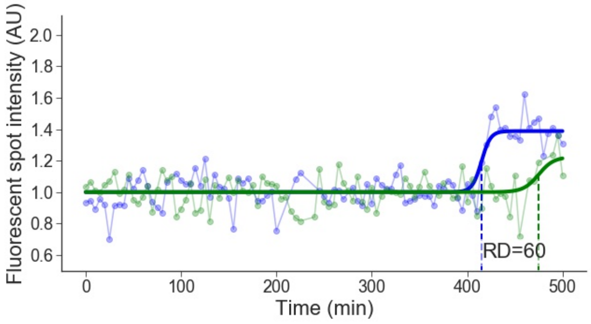


Cell line: Late 2, cell: S28 cell 1

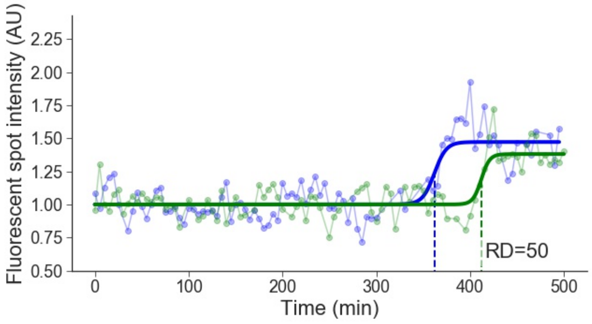


Cell line: Late 2, cell: S29 cell 1

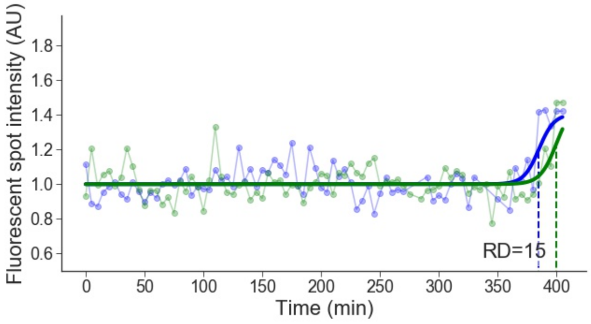

Cell line: Late 2, cell: S29 cell 2

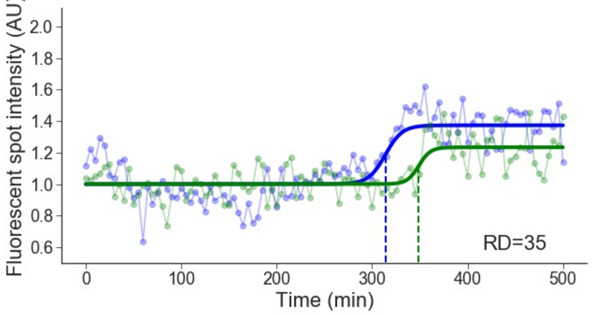


Cell line: Late 2, cell: S30 cell 1

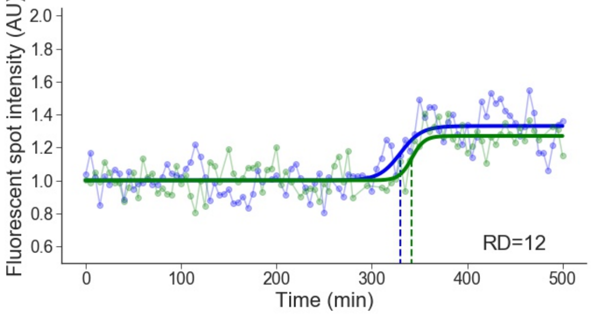


Cell line: Late 2, cell: S30 cell 2

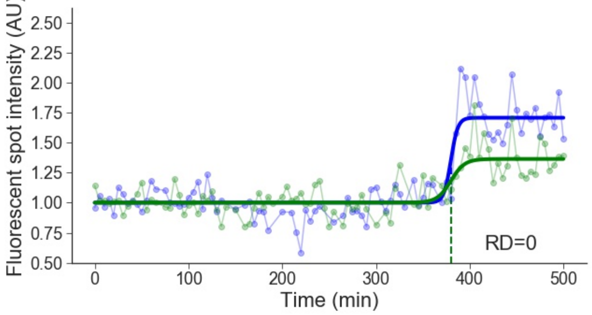


Cell line: Late 2, cell: S31 cell 2

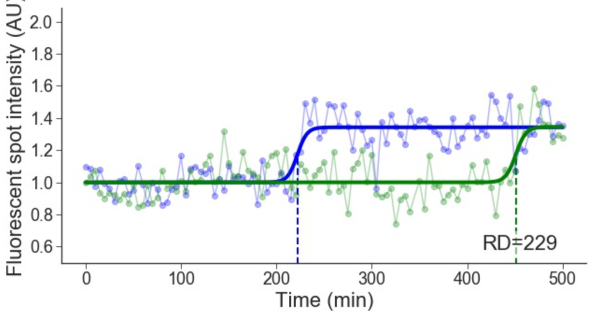


Cell line: Late 2, cell: S31 cell 4

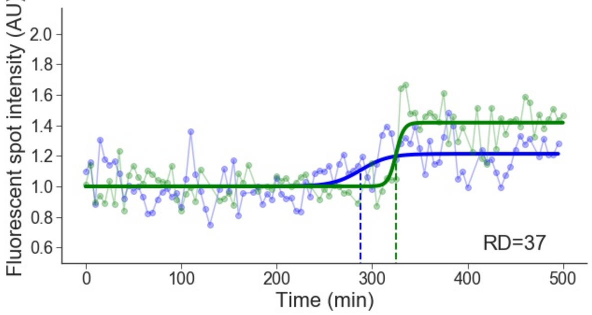


Cell line: Late 2, cell: S34 cell 1

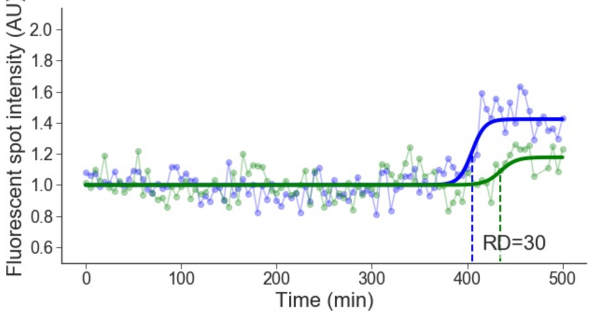


Cell line: Late 2, cell: S34 cell 3

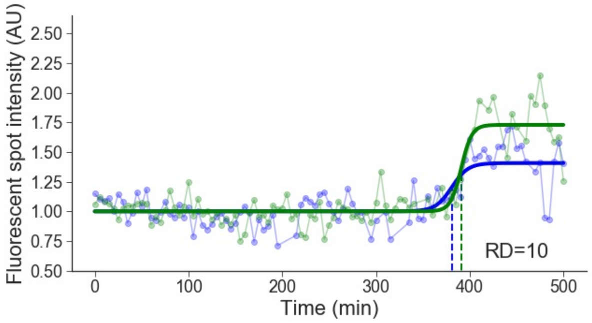


Cell line: Late 2, cell: S37 cell 1

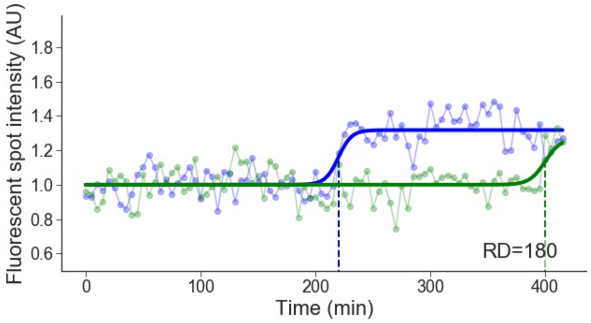

Cell line: Late 2, cell: S40 cell 1

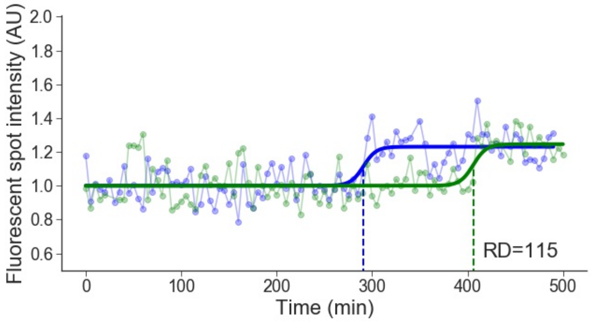


Cell line: Late 2, cell: S40 cell 2

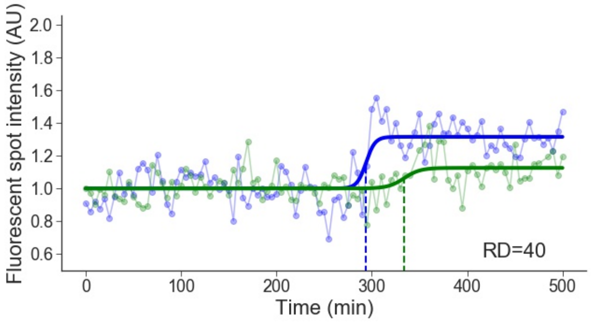


Cell line: Late 2, cell: S41 cell 3

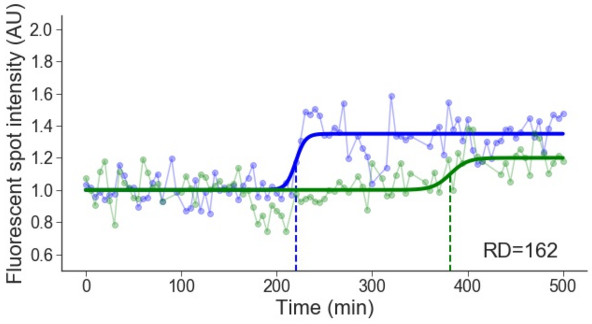


Cell line: Late 2, cell: S42 cell 1

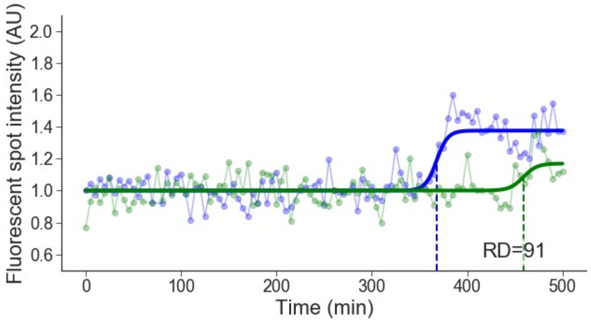


Cell line: Late 2, cell: S43 cell 1

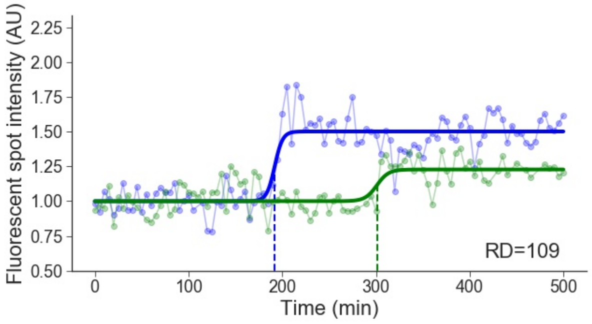


Cell line: Late 2, cell: S44 cell 1

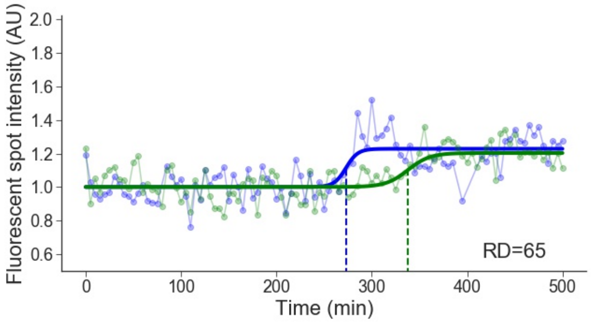


Cell line: Late 2, cell: S46 cell 1

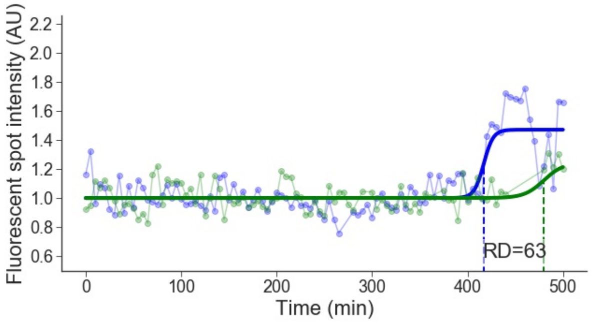


Cell line: Late 2, cell: S46 cell 2

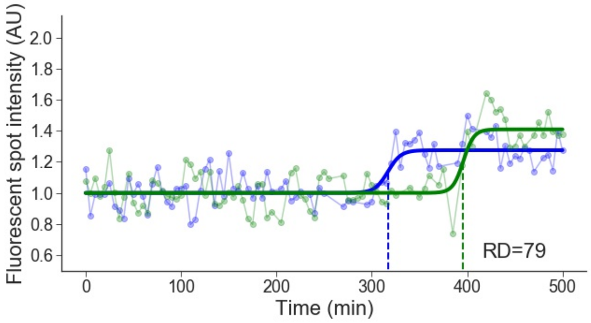

Cell line: Late 2, cell: S46 cell 3

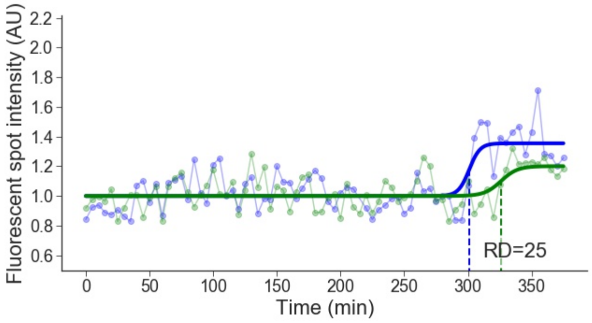


Cell line: Late 2, cell: S48 cell 1

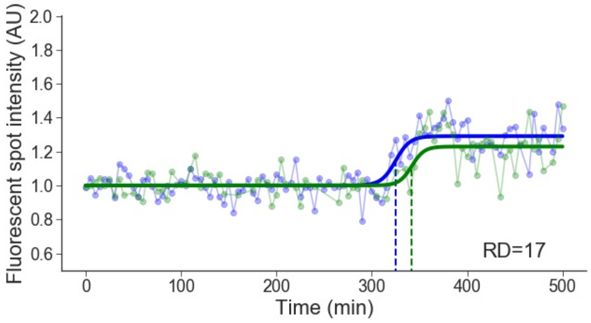


Cell line: Late 2, cell: S49 cell 1

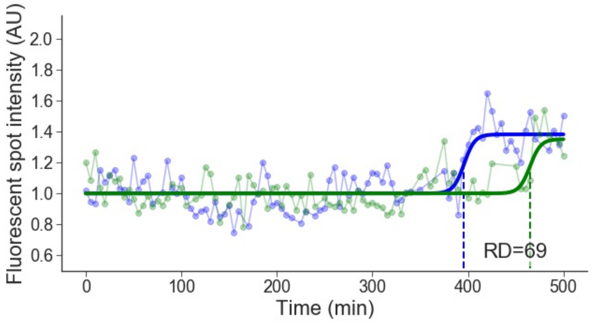


Cell line: Late 2, cell: S49 cell 2

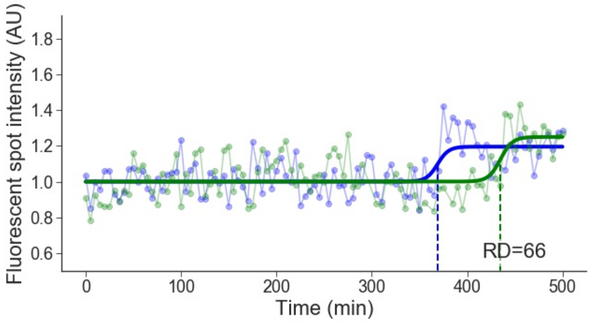


Cell line: Late 2, cell: S50 cell 1

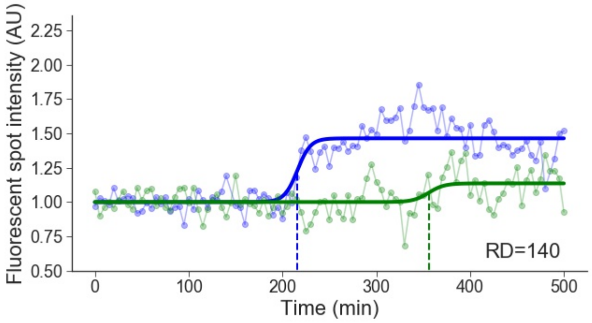


Cell line: Late 2, cell: S50 cell 2

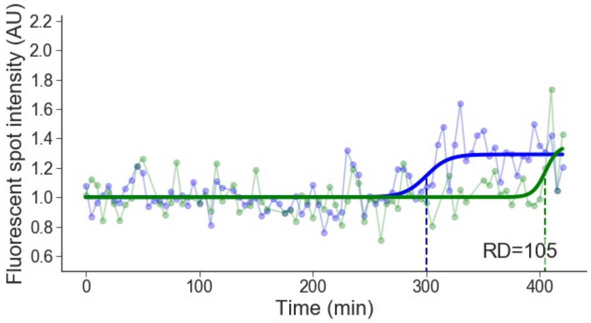

**Supplementary Table 1 : Primer sequences**

| Use | Name | Sequence (5'>3') | Short description | Position in the DT40 genome (Build galGal5, Dec. 2015) |
|---|---|---|---|---|
| **Plasmid construction** | | | | |
| TetR cloning | TetR FOR | CTTCGAATTCACCATGGATCCAAAAAAGAAGAGAAAGGTAGATCCAAAAAAGAAGAGAAAGGTAATGGTGTCTAGATTAGATAAAAG | TetR insertion into the pEGFPN1 vector, flanked with 2 nuclear localization signal sequences | - |
| TetR cloning | TetR REV | TACCGGTACCGCAGACCCACTTTCACATTTAAGTTGTTTTTCTAATCCGCATATGATC | TetR insertion into the pEGFPN1 vector | - |
| TetO array cloning | TetO FOR | GGGGACAACTTTGTATACAAAAGTTG GGCTCGTATGTTGTGTGGAA | Primer flanked with attB5 sequence for TetO array cloning in pDONR 221 P5-P4 | - |
| TetO array cloning | TetO REV | GGGGACAACTTTGTATAGAAAAGTTGGGTG GTAACATCGTTGCTGCTCCA | Primer flanked with attB4 sequence for TetO array cloning in pDONR 221 P5-P4 | - |
| TetO array insertion in DT40 cells in Early 1 locus | Early1 5'arm FOR | GGGGACAAGTTTGTACAAAAAAGCAGGCT TGCTTCTCACCCACACACTT | Primer flanked with attB1 sequence for 5'arm cloning in pDONR 221 P1-P5r, for the insertion of TetO array in Early 1 region | Early 1 - 5' arm: chr1:91,742,981-91,744,494 |
| TetO array insertion in DT40 cells in Early 1 locus | Early1 5'arm REV | GGGGACAACTTTTGTATACAAAGTTGT TAGGCTGCCGTGTTCTTTCT | Primer flanked with attB5r sequence for 5'arm cloning in pDONR 221 P1-P5r, for the insertion of TetO array in Early 1 region | |
| TetO array insertion in DT40 cells in Early 1 locus | Early1 3'arm FOR | GGGGACAACTTTGTATAATAAAGTTG TGGAGAAAAGGGGAGACAGA | Primer flanked with attB3 sequence for 3'arm cloning in pDONR 221 P3-P2, for the insertion of TetO array in Early 1 region | Early 1 - 3' arm: chr1:91,744,624-91,746,104 |
| TetO array insertion in DT40 cells in Early 1 locus | Early1 3'arm REV | GGGGACCACTTTGTACAAGAAAGCTGGGTA AGCAGTGGCAGTGTGAGAGA | Primer flanked with attB2 sequence for 3'arm cloning in pDONR 221 P3-P2, for the insertion of TetO array in Early 1 region | |
| TetO array insertion in DT40 cells in Early 2 locus | Early2 5'arm FOR | GGGGACAAGTTTGTACAAAAAAGCAGGCT CAGGCTGGGACATATCACAA | Primer flanked with attB1 sequence for 5'arm cloning in pDONR 221 P1-P5r, for the insertion of TetO array in Early 2 region | Early 2 - 5' arm: chr1:112,533,062-112,535,191 |
| TetO array insertion in DT40 cells in Early 2 locus | Early2 5'arm REV | GGGGACAACTTTTGTATACAAAGTTGT CGGATCTCTCCTAAGCCACT | Primer flanked with attB5r sequence for 5'arm cloning in pDONR 221 P1-P5r, for the insertion of TetO array in Early 2 region | |
| TetO array insertion in DT40 cells in Early 2 locus | Early2 3'arm FOR | GGGGACAACTTTGTATAATAAAGTTG TCCCAGCAGGGCTTACTCT | Primer flanked with attB3 sequence for 3'arm cloning in pDONR 221 P3-P2, for the insertion of TetO array in Early 2 region | Early 2 - 3' arm: chr1:112,535,245-112,536,822 |
| TetO array insertion in DT40 cells in Early 2 locus | Early2 3'arm REV | GGGGACCACTTTGTACAAGAAAGCTGGGTA CTACAGGGCTTGTGGAGCAA | Primer flanked with attB2 sequence for 3'arm cloning in pDONR 221 P3-P2, for the insertion of TetO array in Early 2 region | |
| TetO array insertion in DT40 cells in Mid-late 1 locus | Mid-late1 5'arm FOR | GGGGACAAGTTTGTACAAAAAAGCAGGCT CCAAACCAGGCCACTCTTAGT | Primer flanked with attB1 sequence for 5'arm cloning in pDONR 221 P1-P5r, for the insertion of TetO array in Mid-late 1 region | Mid-late 1 - 5' arm: chr1:72,546,427-72,548,589 |
| TetO array insertion in DT40 cells in Mid-late 1 locus | Mid-late1 5'arm REV | GGGGACAACTTTTGTATACAAAGTTGT AGTCACTTGGCATAAATAAGAAGCC | Primer flanked with attB5r sequence for 5'arm cloning in pDONR 221 P1-P5r, for the insertion of TetO array in Mid-late 1 region | |
| TetO array insertion in DT40 cells in Mid-late 1 locus | Mid-late1 3'arm FOR | GGGGACAACTTTGTATAATAAAGTTG CTGAGCAGGAAGGGAAACGA | Primer flanked with attB3 sequence for 3'arm cloning in pDONR 221 P3-P2, for the insertion of TetO array in Mid-late 1 region | Mid-late 1 - 3' arm: chr1:72,548,595-72,550,662 |
| TetO array insertion in DT40 cells in Mid-late 1 locus | Mid-late1 3'arm REV | GGGGACCACTTTGTACAAGAAAGCTGGGTA CCATAGTGCAGACCTGGCAT | Primer flanked with attB2 sequence for 3'arm cloning in pDONR 221 P3-P2, for the insertion of TetO array in Mid-late 1 region | |
| TetO array insertion in DT40 cells in Mid-late 2 locus | Mid-late2 5'arm FOR | GGGGACAAGTTTGTACAAAAAAGCAGGCT ATGGCAGCAGAGCGTGCAGA | Primer flanked with attB1 sequence for 5'arm cloning in pDONR 221 P1-P5r, for the insertion of TetO array in Mid-late 2 region | Mid-late 2 - 5' arm: chr1:115,488,660-115,490,676 |
| TetO array insertion in DT40 cells in Mid-late 2 locus | Mid-late2 5'arm REV | GGGGACAACTTTTGTATACAAAGTTGT TAGGCAGCACCTGGTCCTGT | Primer flanked with attB5r sequence for 5'arm cloning in pDONR 221 P1-P5r, for the insertion of TetO array in Mid-late 2 region | |
| TetO array insertion in DT40 cells in Mid-late 2 locus | Mid-late2 3'arm FOR | GGGGACAACTTTGTATAATAAAGTTG GGTGGCGTGCATCAGTCAGA | Primer flanked with attB3 sequence for 3'arm cloning in pDONR 221 P3-P2, for the insertion of TetO array in Mid-late 2 region | Mid-late 2 - 3' arm: chr1:115,490,844-115,492,903 |
| TetO array insertion in DT40 cells in Mid-late 2 locus | Mid-late2 3'arm REV | GGGGACCACTTTGTACAAGAAAGCTGGGTA AGCTCTGTAAGGTCGCTCGC | Primer flanked with attB2 sequence for 3'arm cloning in pDONR 221 P3-P2, for the insertion of TetO array in Mid-late 2 region | |
| TetO array insertion in DT40 cells in Late 1 locus | Late1 5'arm FOR | GGGGACAAGTTTGTACAAAAAAGCAGGCT CAGTGAAACACAGGAGGAACA | Primer flanked with attB1 sequence for 5'arm cloning in pDONR 221 P1-P5r, for the insertion of TetO array in Late 1 region | Late 1 - 5' arm: chr1:70,521,406-70,523,649 |
| TetO array insertion in DT40 cells in Late 1 locus | Late1 5'arm REV | GGGGACAACTTTTGTATACAAAGTTGT TAACTCCAAGAACGATCACTGC | Primer flanked with attB5r sequence for 5'arm cloning in pDONR 221 P1-P5r, for the insertion of TetO array in Late 1 region | |
| TetO array insertion in DT40 cells in Late 1 locus | Late1 3'arm FOR | GGGGACAACTTTGTATAATAAAGTTG GGAAATGTCTTGAATCTCACAAGG | Primer flanked with attB3 sequence for 3'arm cloning in pDONR 221 P3-P2, for the insertion of TetO array in Late 1 region | Late 1 - 3' arm: chr1:70,523,730-70,525,866 |
| TetO array insertion in DT40 cells in Late 1 locus | Late1 3'arm REV | GGGGACCACTTTGTACAAGAAAGCTGGGTA ATGCCCACCAGTGTCCATAA | Primer flanked with attB2 sequence for 3'arm cloning in pDONR 221 P3-P2, for the insertion of TetO array in Late 1 region | |
| TetO array insertion in DT40 cells in Late 2 locus | Late2 (site 1) 5'arm FOR | GGGGACAAGTTTGTACAAAAAAGCAGGCT ACTTGTTGAGCCTTTATGGAGAAC | Primer flanked with attB1 sequence for 5'arm cloning in pDONR 221 P1-P5r, for the insertion of TetO array in the site 1 of Late 2 region | Late 2 (site 1) - 5' arm: chr1:177,934,197-177,936,192 |
| TetO array insertion in DT40 cells in Late 2 locus | Late2 (site 1) 5'arm REV | GGGGACAACTTTTGTATACAAAGTTGT CGGTGTTACAGAGGAGTAAACTGA | Primer flanked with attB5r sequence for 5'arm cloning in pDONR 221 P1-P5r, for the insertion of TetO array in the site 1 of Late 2 region | |
| TetO array insertion in DT40 cells in Late 2 locus | Late2 (site 1) 3'arm FOR | GGGGACAACTTTGTATAATAAAGTTG ACCTTATGCATTTCGTTCCTATGT | Primer flanked with attB3 sequence for 3'arm cloning in pDONR 221 P3-P2, for the insertion of TetO array in the site 1 of Late 2 region | Late 2 (site 1) - 3' arm: chr1:177,936,522-177,938,522 |
| TetO array insertion in DT40 cells in Late 2 locus | Late2 (site 1) 3'arm REV | GGGGACCACTTTGTACAAGAAAGCTGGGTA TAAGAAGAGAGATGGGGATCAAAC | Primer flanked with attB2 sequence for 3'arm cloning in pDONR 221 P3-P2, for the insertion of TetO array in the site 1 of Late 2 region | |
| TetO array insertion in DT40 cells in Late 2 locus | Late2 (site 2) 5'arm FOR | GGGGACAAGTTTGTACAAAAAAGCAGGCT GAGAGCAGCTAGTGGGAAGG | Primer flanked with attB1 sequence for 5'arm cloning in pDONR 221 P1-P5r, for the insertion of TetO array in the site 2 of Late 2 region | Late 2 (site 2) - 5' arm: chr1:177,929,394-177,930,911 |
| TetO array insertion in DT40 cells in Late 2 locus | Late2 (site 2) 5'arm REV | GGGGACAACTTTTGTATACAAAGTTGT CTTCAGGTACGCAGGGATGA | Primer flanked with attB5r sequence for 5'arm cloning in pDONR 221 P1-P5r, for the insertion of TetO array in the site 2 of Late 2 region | |
| TetO array insertion in DT40 cells in Late 2 locus | Late2 (site 2) 3'arm FOR | GGGGACAACTTTGTATAATAAAGTTG GGATGTCACCCACAAGATCAG | Primer flanked with attB3 sequence for 3'arm cloning in pDONR 221 P3-P2, for the insertion of TetO array in the site 2 of Late 2 region | Late 2 (site 2) - 3' arm: chr1:177,931,024-177,933,038 |
| TetO array insertion in DT40 cells in Late 2 locus | Late2 (site 2) 3'arm REV | GGGGACCACTTTGTACAAGAAAGCTGGGTA CGGGGATAGAAGTGGACACAA | Primer flanked with attB2 sequence for 3'arm cloning in pDONR 221 P3-P2, for the insertion of TetO array in the site 2 of Late 2 region | |
| **Cell line construction** | | | | |
| Test of TetO array insertion in Early 1 locus | 3' Early1 site Reverse | GTAACATGGAATCCTGCCTTGA | Test of targeted TetO insertion and of Blasticidin resistance cassette excision in Early 1 region | chr1: 91,746,107-91,746,128 |
| Test of TetO array insertion in Early 2 locus | 3' Early2 site Reverse | AGGGAATACCTTCCACGAGTCT | Test of targeted TetO insertion in Early 2 region | chr1: 112,536,855-112,536,876 |
| Test of TetO array insertion in Early 2 locus | 3'arm Early2 Reverse | CACATGGGGATGGATTTAGG | Test of Blasticidin resistance cassette excision in Early 2 region | chr1:112,535,684-112,535,703 |
| Test of TetO array insertion in Mid-late 1 locus | 3' Mid-late1 site Reverse | AGGACGGAAAAGAGGCAAAGA | Test of targeted TetO insertion and of Blasticidin resistance cassette excision in Mid-late 1 region | chr1:72,550,854-72,550,874 |
| Test of TetO array insertion in Mid-late 2 locus | 3' Mid-late2 site Reverse | GTTACATGGCCAAGGTTGCT | Test of targeted TetO insertion in Mid-late 2 region | chr1:115,493,100-115,493,119 |
| Test of TetO array insertion in Mid-late 2 locus | 3'arm Mid-late2 Reverse | CATCACAGAGGGAGATGCAA | Test of Blasticidin resistance cassette excision in Mid-late 2 region | chr1:115,490,941-115,490,960 |
| Test of TetO array insertion in Late 1 locus | 3' Late1 site Reverse | ACTATTGTACCCCTCCCTGTTG | Test of targeted TetO insertion in Late 1 region | chr1:70,526,230-70,526,251 |
| Test of TetO array insertion in Late 1 locus | 3'arm Late1 Reverse | TGGGAGTGGGGTATTGGTAA | Test of Blasticidin resistance cassette excision in Late 1 region | chr1:70,524,054-70,524,073 |
| Test of TetO array insertion in Late 2 locus | 3' Late2 (site 1) site Reverse | ATCTCTGCCTTCAAACCTTCAG | Test of targeted TetO insertion and of Blasticidin resistance cassette excision in site 1 of Late 2 region | chr1: 177,938,826-177,938,847 |
| Test of TetO array insertion in Late 2 locus | 3' Late2 (site 2) site Reverse | CATTCAACATTTGTCTGCTTTTCT | Test of targeted TetO insertion in site 2 of Late 2 region | chr1:177,933,211-177,933,234 |
| Test of TetO array insertion in Late 2 locus | 3'arm Late2 (site 2) Reverse | TCCAGATTTGTATTTGTCACTTAACC | Test of Blasticidin resistance cassette excision in site 2 of Late 2 region | chr1:177,931,930-177,931,955 |
| Test of TetO array insertion | TetO Forward 1 | TAGAGAAGATGGGGGCCTCT | Test of Blasticidin resistance cassette excision (in Early 1, Early 2, Mid-late 1, Mid-late 2, Late 1 and site 1 of Late 2 regions) | - |
| Test of TetO array insertion | TetO Forward 2 | TCATGGCTTGTTATGACTGTTTTT | Test of Blasticidin resistance cassette excision (in site 2 of Late 2 regions) | - |
| Test of TetO array insertion | Bls Forward 1 | CCCCCTGAACCTGAAACATA | Test of targeted TetO insertion in Early 1, Early 2, Mid-late 1, Mid-late 2, late 1 and site 1 of Late 2 regions | - |
| Test of TetO array insertion | Bls Forward 2 | TGCATTCTAGTTGTGGTTTGTCC | Test of targeted TetO insertion in site 2 of Late 2 regions | - |
| **Timing assays** | | | | |
| Timing assays in Early 1 locus | Without Early1 FOR | AGAAAGAACACGGCAGCCTA | qPCR in Timing assays with Early 1 cell line (Without Early 1) | Without Early 1: chr1:91,744,475-91,744,598 |
| Timing assays in Early 1 locus | Without Early1 REV | CCCCCTTCTCTGTCTGTCAA | qPCR in Timing assays with Early 1 cell line (Without Early 1) | |
| Timing assays in Early 1 locus | Both Early1 FOR | TCGTACAGGGCCAGTCTTTCT | qPCR in Timing assays with Early 1 cell line (Both Early 1) | Both Early 1: chr1:91,751,582-91,751,703 |
| Timing assays in Early 1 locus | Both Early1 REV | CTGCGCCATATCAAGGCTAC | qPCR in Timing assays with Early 1 cell line (Both Early 1) | |
| Timing assays in Early 2 locus | Without Early2 FOR | TCTCCAAAAAGTGGCTTAGGAG | qPCR in Timing assays with Early 2 cell line (Without Early 2) | Without Early 2: chr1:112,535,163-112,535,315 |
| Timing assays in Early 2 locus | Without Early2 REV | CATCTCCCCTTCACAGCATC | qPCR in Timing assays with Early 2 cell line (Without Early 2) | |
| Timing assays in Early 2 locus | Both Early2 FOR | GAGGAAGAGGGGAGGGAGAC | qPCR in Timing assays with Early 2 cell line (Both Early 2) | Both Early 2: chr1:112,530,432-112,530,540 |
| Timing assays in Early 2 locus | Both Early2 REV | GTGAGCTTGGTGTGGAGTTTG | qPCR in Timing assays with Early 2 cell line (Both Early 2) | |
| Timing assays in Mid-late 1 locus | Without Mid-late1 FOR | GGCTTCTTATTTATGCCAAGTGA | qPCR in Timing assays with Mid-late 1 cell line (Without Mid-late 1) | Without Mid-late 1: chr1:72,548,565-72,548,659 |
| Timing assays in Mid-late 1 locus | Without Mid-late1 REV | TGCATCAGCAATGAGCAAAC | qPCR in Timing assays with Mid-late 1 cell line (Without Mid-late 1) | |
| Timing assays in Mid-late 1 locus | Both Mid-late1 FOR | TCTTCCAACCTAAGCCACTCTG | qPCR in Timing assays with Mid-late 1 cell line (Both Mid-late 1) | Both Mid-late 1: chr1:72,553,336-72,553,456 |
| Timing assays in Mid-late 1 locus | Both Mid-late1 REV | TGCATATTTTCCCCGAGTCTT | qPCR in Timing assays with Mid-late 1 cell line (Both Mid-late 1) | |
| Timing assays in Mid-late 2 locus | Without Mid-late2 FOR | TTGGCTCTAGCTTGCACTCA | qPCR in Timing assays with Mid-late 2 cell line (Without Mid-late 2) | Without Mid-late 2: chr1:115,490,690-115,490,817 |
| Timing assays in Mid-late 2 locus | Without Mid-late2 REV | GCAAAAGCTGGCTGAATTTT | qPCR in Timing assays with Mid-late 2 cell line (Without Mid-late 2) | |
| Timing assays in Mid-late 2 locus | Both Mid-late2 FOR | TGGCCTGTCAAAGAGGAGAG | qPCR in Timing assays with Mid-late 2 cell line (Both Mid-late 2) | Both Mid-late 2: chr1:115,481,076-115,481,175 |
| Timing assays in Mid-late 2 locus | Both Mid-late2 REV | GCCAGCCCTACTCAGACAAA | qPCR in Timing assays with Mid-late 2 cell line (Both Mid-late 2) | |
| Timing assays in Late 1 locus | Without Late1 FOR | TGGAGAAAGGAGGGCTTTTTA | qPCR in Timing assays with Late 1 cell line (Without Late 1) | Without Late 1: chr1:70,523,661-70,523,767 |
| Timing assays in Late 1 locus | Without Late1 REV | GCCTCTTCAGTGTCCCTTGT | qPCR in Timing assays with Late 1 cell line (Without Late 1) | |
| Timing assays in Late 1 locus | Both Late1 FOR | TGTACTTCTCTGTGGACATGCA | qPCR in Timing assays with Late 1 cell line (Both Late 1) | Both Late 1: chr1:70,518,727-70,518,810 |
| Timing assays in Late 1 locus | Both Late1 REV | TGGCACAGAGGACAGGTAAGA | qPCR in Timing assays with Late 1 cell line (Both Late 1) | |
| Timing assays in Late 2 locus | Without Late2 FOR | CTTTGAGAGCATGTGGTAATACGG | qPCR in Timing assays with Late 2 cell line (Without Late 2) | Without Late 2: chr1:177,936,263-177,936,368 |
| Timing assays in Late 2 locus | Without Late2 REV | TACTGTATCGAGGGGTGGGAGTAG | qPCR in Timing assays with Late 2 cell line (Without Late 2) | |
| Timing assays in Late 2 locus | Both Late2 FOR | GGAGAGAAAGCGGTGATGTG | qPCR in Timing assays with Late 2 cell line (Both Late 2) | Both Late 2: chr1:177,941,044-177,941,153 |
| Timing assays in Late 2 locus | Both Late2 REV | GGACCTGCTGCATGATTGTT | qPCR in Timing assays with Late 2 cell line (Both Late 2) | |
| Timing assays | With TetO FOR 1 | GCGGTTTTCATGGCTTGT | qPCR in Timing assays (With TetO) (primer pair: With TetO FOR1/With TetO REV 1) | - |
| Timing assays | With TetO REV 1 | CGTTGCTGCTCCATAACATC | qPCR in Timing assays (With TetO) (primer pair: With TetO FOR1/With TetO REV 1) | - |
| Timing assays | With TetO FOR 2 | TCAGGATAGGAGGGGGCTAC | qPCR in Timing assays (Witrh TetO) (primer pair: With TetO FOR 2/With TetO REV 2) | - |
| Timing assays | With TetO REV 2 | TCACTGATGGGATGGCTCTT | qPCR in Timing assays (Witrh TetO) (primer pair: With TetO FOR 2/With TetO REV 2) | - |
| Timing assays | With TetO FOR 3 | CGAGGGGGTCCCTATCAGT | qPCR in Timing assays (With TetO) (primer pair: With TetO FOR 3/With TetO REV 3) | - |
| Timing assays | With TetO REV 3 | AGCTGGCCCCATTCTCTATC | qPCR in Timing assays (With TetO) (primer pair: With TetO FOR 3/With TetO REV 3) | - |
| Timing assays, control of FACS sorting | Early control 1 FOR | GACCTCCCCTCGTCTCCC | qPCR in Timing assays, control of very Early timing (Early Control 1) | Early Control 1: chr1:112,227,405-112,227,557 |
| Timing assays, control of FACS sorting | Early control 1 REV | CGTCACGGTCGGGGTTAGC | qPCR in Timing assays, control of very Early timing (Early Control 1) | |
| Timing assays, control of FACS sorting | Early control 2 FOR | TGCGGAAATCACCAGGGAGT | qPCR in Timing assays, control of early timing (Early Control 2) | Early Control 2: chr1:194,564,113-194,564,243 |
| Timing assays, control of FACS sorting | Early control 2 REV | AGTCTAAGCTGCCTCGGGG | qPCR in Timing assays, control of early timing (Early Control 2) | |
| Timing assays, control of FACS sorting | Late control 3 FOR | TCAGTTGTTACGAGACACTTAGCT | qPCR in Timing assays, control of Late timing (Late Control 3) | Late Control 3: chr1:114,319,423-114,319,586 |
| Timing assays, control of FACS sorting | Late control 3 REV | AGGCAACAGCCAGCAAC | qPCR in Timing assays, control of Late timing (Late Control 3) | |
| Timing assays, control of FACS sorting | Mitochondria FOR | CATCCCATGCATAACTCCTG | qPCR in timing assays, normalization of the FACS fractions | chrM:541-731 |
| Timing assays, normalization of the FACS fractions | Mitochondria REV | GTAGTCCAGGCTTCACTTGA | qPCR in timing assays, normalization of the FACS fractions | |
